# Evolutionary bursts drive morphological novelty in the world’s largest skinks

**DOI:** 10.1101/2024.06.27.600807

**Authors:** Ian G. Brennan, David G. Chapple, J. Scott Keogh, Stephen Donnellan

## Abstract

Animal phenotypes evolve and diverge as a result of differing selective pressures and drift. These processes leave unique signatures in patterns of trait evolution, impacting the tempo and mode of morphological macroevolution. While there is a broad understanding of the history of some organismal traits (e.g. body size), there is little consensus about the evolutionary mode of most others. This includes the relative contribution of prolonged (Darwinian gradualist) and episodic (Simpsonian jump) changes towards the evolution of novel morphologies. Here we use new exon-capture and linear morphological datasets to investigate the tempo and mode of morphological evolution in Australo-Melanesian Tiliquini skinks. We generate a well-supported time-calibrated phylogenomic tree from ∼400 nuclear markers for more than 100 specimens including undescribed diversity, and provide unprecedented resolution of the rapid Miocene diversification of these lizards. By collecting a morphological dataset that encompasses the lizard body plan (19 traits across the head, body, limb, and tail) we are able to identify that most traits evolve conservatively but infrequent evolutionary bursts result in morphological novelty. These phenotypic discontinuities occur via rapid rate increases along individual branches, inconsistent with both gradualistic and punctuated equilibrial evolutionary modes. Instead, this ‘punctuated gradualism’ has resulted in the rapid evolution of blue-tongued giants and armored dwarves in the ∼20 million years since colonizing Australia. These results outline the evolutionary pathway towards new morphologies and highlight the heterogeneity of evolutionary tempo and mode, even within individual traits.

## Introduction

Great variations in organismal morphology are expected to accumulate over long periods of time and reflect the varied requirements of different species^1,2^. Some organisms are very small and some are very large, others are brightly colored and others cryptic, and through drift and selection, traits respond and diverge from common forms. Despite the dramatic variety we see in nature, on deep timescales, measures of these differences (variance, disparity, *et al.*) often fail to exceed the expectation of a neutral model where the accumulation of variation is the result of a random evolutionary walk through time (Brownian Motion)^3,4^. In other words, Earth’s flora and fauna realize only a fraction of all possible shapes and sizes^5^. This diversity undershoot suggests bounds on phenotypes as a result of genetic, functional, or developmental constraints^6,7^. If constant and gradual processes are unlikely explanations for the accumulation of trait diversity at deep timescales, then how important are heterogeneous processes at shallower ones?

When traits are assumed to evolve under an unbiased random walk (Brownian Motion), trait variance (*v*) is proportional to evolutionary rate (*σ*^2^), and the accumulation of variance is dictated by elapsed time (*t*; so *v*=*σ*^2^*t*). This exploration of trait space (diffusion) is characterized by trait change along branches drawn from a normal (Gaussian) distribution, and assumes a flat adaptive landscape^8,9^. Given observed trait limits, correlations among traits, and the “clumpiness” of extant morphological diversity, this seems an unlikely expectation^2,10^. In practice, morphological evolution is often concentrated on one or a small number of major axes. The primary axis serves as a ‘line of least resistance’^11^ providing a pool of variation on which selection can act. However evolution is not limited to the major axis, and trajectories along minor axes may also pave a path towards new phenotypes. These competing processes are sometimes called “elaboration” (*along* major axis) and “innovation” (*away* from major axis)^12,13^. However their relative contributions towards organismal macroevolution and the evolution of novel forms has been largely overlooked^13–15^.

Almost 80 years ago G.G. Simpson suggested novelty arose by rapid “jumps” to new adaptive zones across an uneven landscape^16^. To explore this landscape, one potential path requires relaxing our assumption that traits evolve consistently across lineages—changing the *mode* of evolution. Proposed alternative evolutionary models such as pulsed or punctuated processes account for rapid jumps in just such a way—by deviating only in the evolutionary *mode*—leaving variance to accumulate as under BM. Another pathway however requires relaxing our assumption that traits evolve via constant rates through time—changing the *tempo* of evolution. These relaxed-rate methods are implemented in Variable Rates or Multi-Regime models of trait evolution. Traditionally, jumps have been difficult to identify without fossil data. This is due in part to the limited information content at internal nodes of a phylogenetic tree, however, even in fossil studies debate has continued about gradual versus pulsed/punctuated evolution^17,18^. Recent advances in phylogenetic comparative methods however, provide the tools to distinguish between gradual and punctuated evolutionary modes, and suggest that pulsed evolution and rate heterogeneity may be common processes^19–24^. Investigating these patterns and processes in extant systems provides a potentially valuable comparison against the paleontological foundation on which many of these ideas were built.

When studying the morphological evolution of any empirical group differences in evolutionary mode and tempo are potentially compounded by mosaic evolution—which suggests that processes may vary widely across individual traits and the modules they make up. So when looking for macroevolutionary shifts towards new phenotypes, changes may be temporally, phylogenetically, or regionally (across the body) heterogeneous. Just a century ago these complications seemed impossible to address: “To select a few of the great number of structural differences for measurement would be almost certainly misleading; to average them all would entail many thousands of measurements for each species or genus compared”^25^. Modern comparative methods however, have made these comparisons possible^26–28^. Here we investigate the relative roles these sources of variation play, by focusing on the Tiliquini skinks. Tiliquines consist of ∼62 species with varied ecologies, high levels of sociality^29,30^, and highly imbalanced biogeographic richness. The majority of species (∼48 spp.) are endemic to Australia where their distributions span the continent’s highest alpine peaks and most inhospitable deserts. To survive in such varied climes, tiliquines have diverged into herbivorous giants that roam the treetops, spiky socialites that live communally in rock crevices or complex burrows, and elongate long-lived slow-movers that wander across open lands. These ecotypes have diagnosable morphologies, and the modest species richness of the group allows high-dimensional macroevolutionary study to be computationally tractable. Additionally, tiliquines are ideally suited for studies of the tempo and pace of morphological evolution because they have been suggested to show a morphological gradient with highly derived phenotypes nested deeply within a generally conserved body plan^30,31^.

But to what degree are these morphological deviations *remarkable*, and what tempo and mode have they evolved by? To answer these questions we started by generating an exon-capture dataset for Tiliquini skinks and reconstructing the phylogenetic relationships of this group. To look at morphological evolution we collected an extensive phenotypic dataset of 19 linear measurements which summarize broad axes of variation across the head, body, limbs, and tail. Finally we incorporate these data into a multivariate framework for comparing evolutionary rates and disparity of traits (and the modules they compose) relative to a neutral BM model of evolution. From this we are able to show that (1) most traits show heterogeneous—not neutral or incremental—evolutionary histories, (2) evolutionary bursts are temporally and phylogenetically distributed but uncommon, and (3) these jumps result in morphological novelty that can exceed uncorrelated trait expectations.

## Results

### Phylogenetic Analyses and Divergence Dating

We present a well-resolved phylogeny that provides unprecedented resolution of Tiliquini skinks representing all genera and 63 taxa (102 samples) across this continental lizard radiation. Concatenated (Fig.S17) and coalescent-consistent species trees (Fig.1) return broadly similar topologies with the exception of a handful of extremely short internal branches (Fig.S18). Discordant branches fall within the anomaly zone^32^ in which concatenation is likely to be misled by the contribution of a large number of anomalous gene trees. In support of this, these nodes as represented in our coalescent species tree generally have low gene concordance factors (gCFs) and equivocal site concordance factors (sCFs), but are strongly supported by individual topology tests (see Supplementary Materials Topology Tests). Of these contentious nodes, two have taxonomic implications. *Lissolepis* is not recovered as monophyletic, with *L. coventryi* estimated as sister to *Liopholis*, and *L. luctuosa* more closely related to the remaining tiliquines (*Cyclodomorphus*, *Tiliqua*, *Bellatorias*, *Egernia*). We also find *Cyclodomorphus* to be non-monophyletic, with the *C. gerrardii* group more closely allied with *Tiliqua* than with the *C. maximus* group.

**Figure 1.**
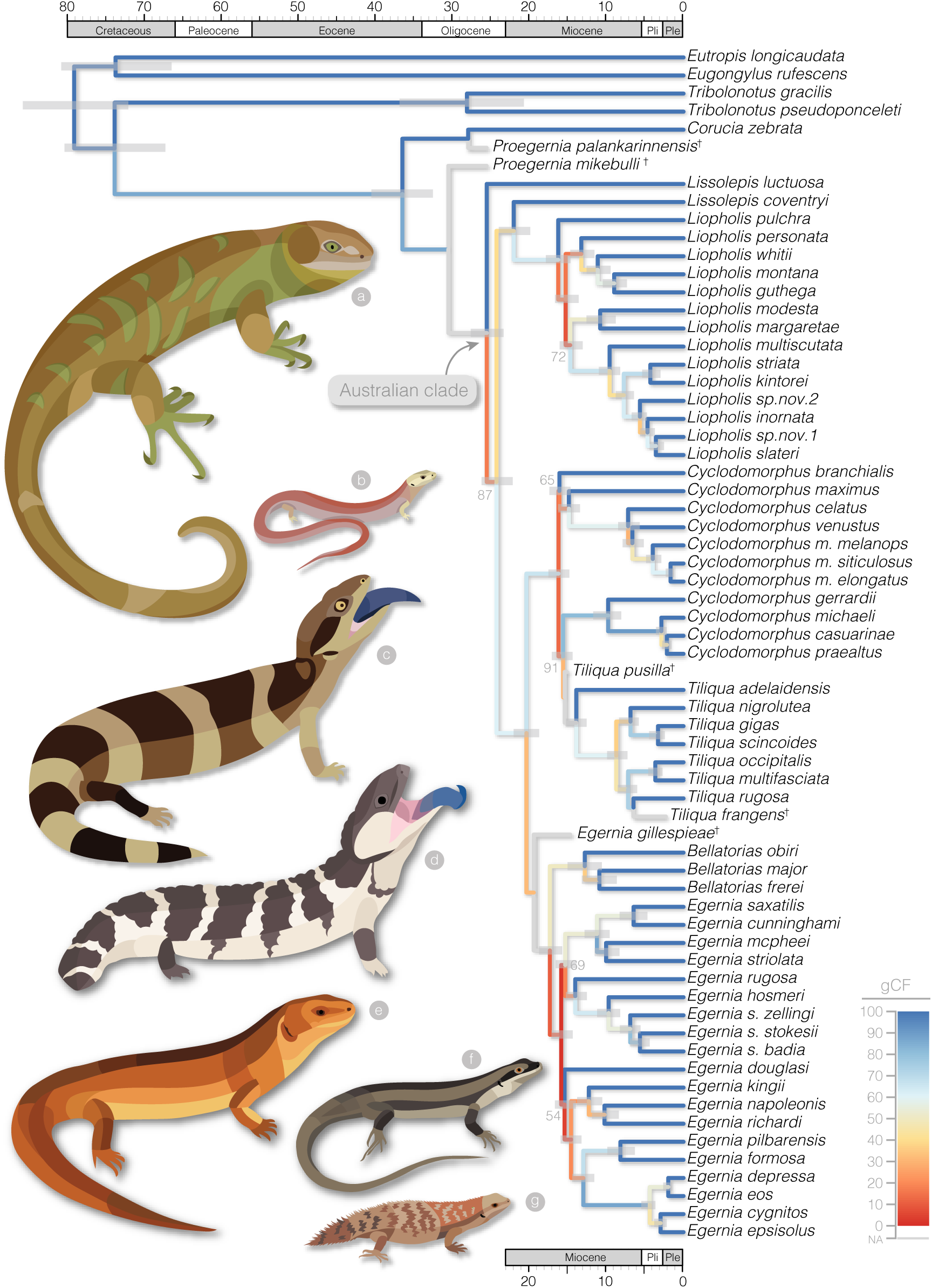
Species tree estimated with ASTRAL and time-calibrated with MCMCtree shows late Oligocene or early Miocene divergences of most major Australian Tiliquini skinks, in contrast to a Cretaceous divergence from *Tribolonotus* and late Eocene divergence from the monotypic *Corucia*. Local posterior supports for all nodes are 100, except where indicated by grey numbers at nodes. Branches are colored according to gene concordance factors (gCF)—the proportion of gene trees which support the given bifurcation—to highlight areas of discordance among genetrees. For illustrative purposes fossil taxa (*†*) have been placed in the dated molecular tree following a combined evidence analysis in BEAST. Animals illustrate the diversity of size and shape across the Tiliquini: (a) *Corucia zebrata*, (b) *Cyclodomorphus michaeli*, (c) *Tiliqua occipitalis*, (d) *Tiliqua rugosa*, (e) *Egernia rugosa*, (f) *Egernia striolata*, (f) *Egernia depressa*.

Branches that define many of the inter-generic relationships among Australian tiliquines are the result of a series of rapid speciation events occurring 15–21 mya. Most of these branches are shorter than a million years long, and several of which are less than 500k years long. These rapid divergences contrast with the deep splits between Australian tiliquines and their sister taxon *Corucia* (30 mya; 95% HPD: 27–35 mya), as well as the preceding split from *Tribolonotus* (60 mya; 95% HPD: 55-64 mya) (Fig.1,S1).

### Phenotypic Analyses

Modularity and integration model selection in *EMMLi*^33^ identified a four module model with separate within module correlations (intramodule integration varies among modules) and separate among module correlations (intermodule integration varies among modules) as the best fit to our data.

15 of 19 morphological traits are best fit by variable rate or pulsed models (Fig.S15) as well as the summary trait encapsulating size. The abundant preference for heterogeneous evolutionary models encouraged us to focus on the evolution of traits and modules under the Variable Rates model for all further analyses. The major axis of morphological variation (elaboration; PC1) across the Tiliquini and across each genus is primarily explained by tail length (Fig.S29—S31). The major axis of morphological innovation (PC2) varies when considering the clade as a whole (interlimb length) or individual genera (primarily size and/or interlimb length). The relationship between elaboration and innovation, however, varies among genera. We visualize this through varied slopes between our first two PC axes, highlighting different paths to phenotypic novelty (Fig.2).

**Figure 2:**
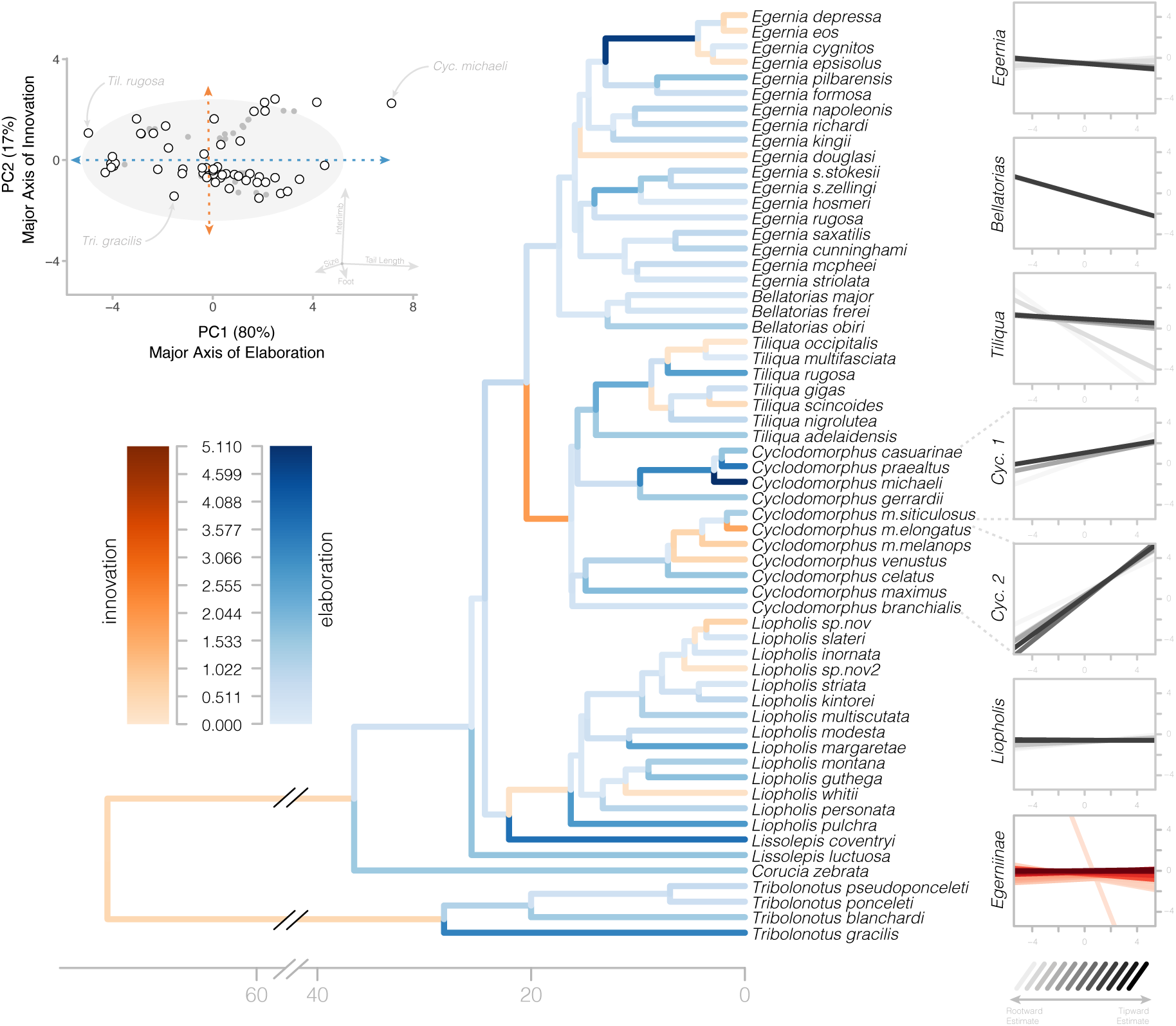
Major and minor axes of morphological change allow us to identify periods of elaboration (along PC1–blues) and innovation (along PC2–oranges) in Tiliquini skinks. Top left–biplot of first two principal components show the distribution of observed species (white circles with black outlines) and estimated ancestors (grey circles) along the major axis of elaboration and innovation. Colored branches on the tree at center indicate the primary direction of morphological change from ancestor to descendant node. Blue branches indicate principally elaborative change, while orange branches indicate principally innovative change. Color saturation (light to dark) indicates the Euclidean distance travelled in PC space along that branch. At right we visualize the varied relationships between these axes among groups. For each clade (genera and group as a whole) we plot the evolution of the relationship between elaborative and innovative axes through time from the root of the clade (lightest regression) to the tips (darkest regression). Regression plots highlight the varied patterns of subclades, including strong conservativism of *Liopholis* and novelty of *Cyclodomorphus*.

Comparing the slopes of disparity accumulation curves highlights periods of significant morphological conservatism (tail module 20–15 mya) and expansion (body and limb modules 18–14 mya; tail module 3 mya–present). Periods of expansion generally—but not always— coincide with periods of increased mean evolutionary rates (Fig.3). Little information can be extracted from the early evolution of this group (75-30 mya), as estimates along bare branches leading to *Tribolonotus* and *Corucia* likely do not reflect the evolution of extinct unsampled lineages along these edges. Discretizing morphological change as primarily elaborative or innovative allows us to identify that both processes happen at clade and species-level scales (Fig.2).

**Figure 3:**
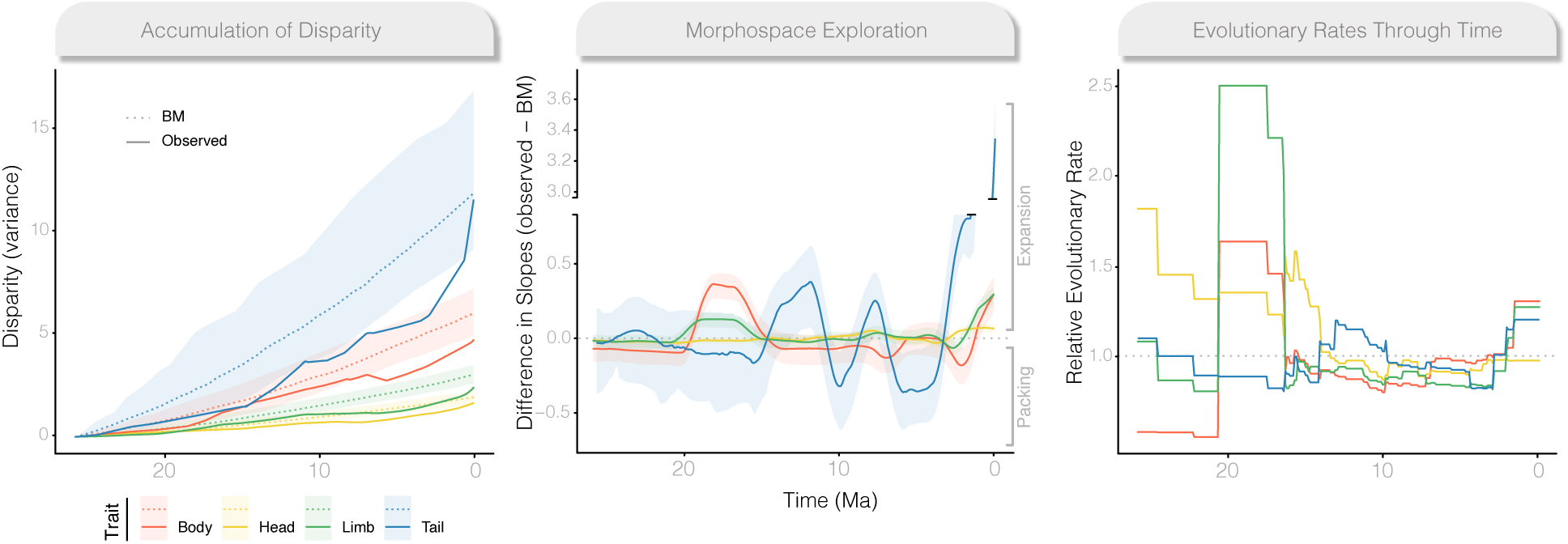
Evolutionary trajectories vary widely across modules and through time. (left) Accumulation of multivariate disparity (as variance) through time for each module (solid lines) compared to Brownian Motion (dotted lines). (center) The comparison of slopes of each module to BM highlights periods of morphological expansion (values greater than 0) and conservatism (values less than 0). (right) Evolutionary rates across modules are highly heterogeneous (see three different scales for y axis), showing periods of temporal variability, as well as high variances within modules and among traits (Fig.S24–S26). Figures represent analyses of the Australian clade alone to avoid bias in estimated rates and diversity as a result of long bare branches leading to *Corucia* and *Tribolonotus*. For comparison with patterns estimated from data including these taxa, see Fig.S32.

Averaging rates however, hides pulses of extreme rate variation (Fig.4). We identified pulses by isolating branches with at least twice the background evolutionary rate (mean scalar *r ≥* 2) that were shifted in at least 70% of the sampling posterior. Across 19 focal traits, roughly 13% branches exhibited a significant rate pulse, with more than 3% showing major shifts in evolutionary rate (mean scalar *r* > 10) (Fig.S12). Major shifts were primarily concentrated in head width, interlimb length, tail length and width, and upper arm length. In modules, nearly 16% of branches exhibited significant rate pulses, with 2% showing major shifts, concentrated in tail and body modules. Many rate increases are concentrated in the *Cyclodomorphus*–*Tiliqua* clade, with particularly rapid rates among *Cyclodomorphus michaeli*, *casuarinae*, and *praealtus*. Body and tail traits also commonly show rate pulses in the crevice dwelling *Egernia stokesii* and *depressa* clades. Under relaxed-rate models rate pulses transfer into bursts in morphological change. We are able to visualize these morphological jumps along individual branches (Fig.6), highlighting instances which lead to novel phenotypes.

**Figure 4.**
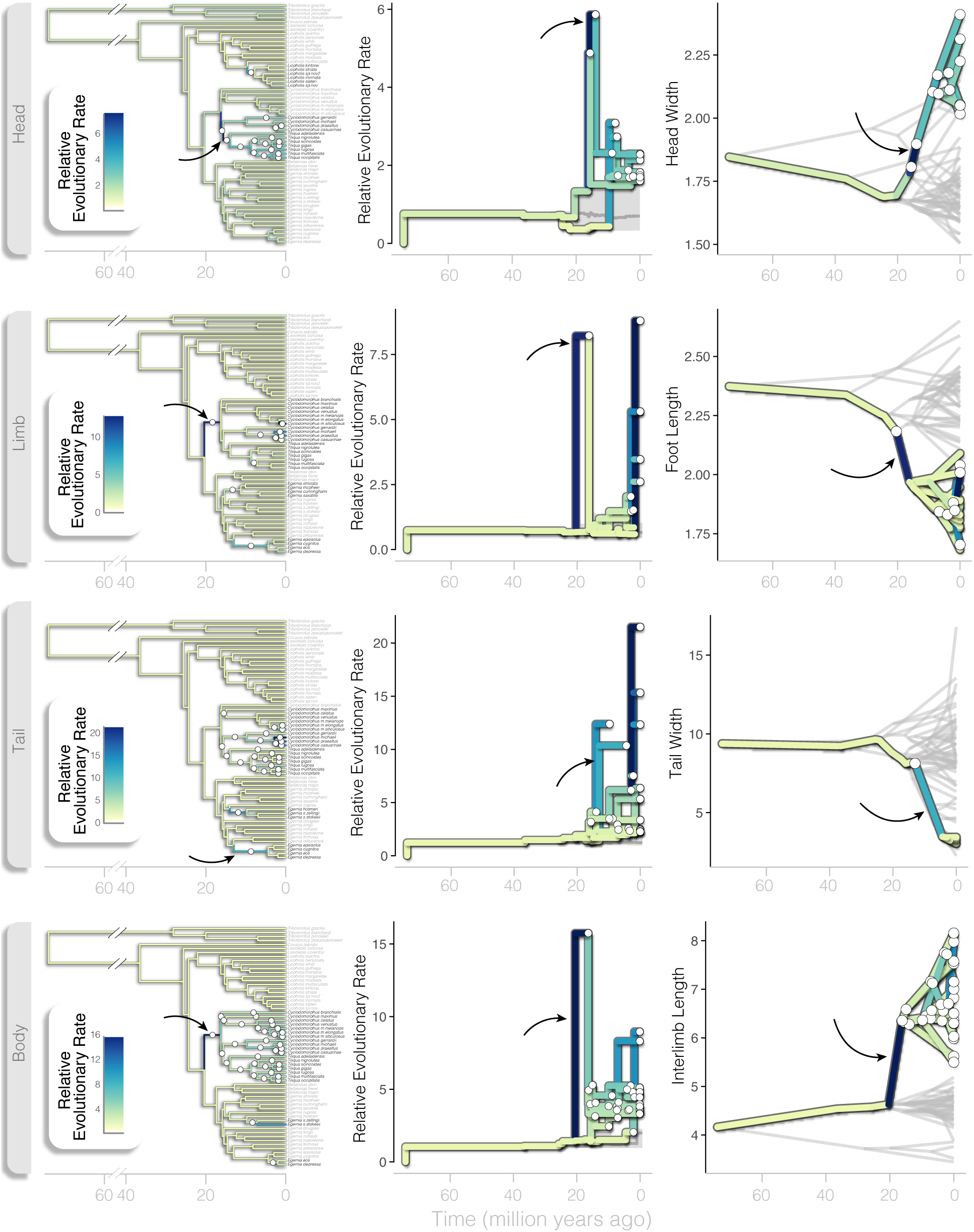
Bursts in rates of phenotypic evolution are distributed across the Tiliquini tree and exhibit strong departures from background rates. Rows represent morphological modules. In all plots branch colors correspond to estimated multivariate evolutionary rates, with significant rate changes noted by white circles at nodes or along branches. (left column) Tiliquini species trees highlighting the location of multivariate rate pulses. (center column) Branch rate trajectories plotted from the root node to nodes that show significant rate shifts (estimated rate scalar *r* > 2). Solid grey envelope contains 95% quantiles of background evolutionary rates (estimated rate scalar *r* < 2) with the mean plotted in dark grey. (right column) Phenograms of an individual trait from each module showing the evolution of extreme phenotypes driven by bursts in evolutionary rate. In each row, a black arrow highlights a single branch of interest across all three plots.

Simulations under uncorrelated and correlated Brownian Motion show differences in the accumulation of trait combinations. This highlights how evolution can be biased along particular axes (Fig.5, S6). Empirical traits generally conform to BM expectations, however extreme phenotypes exceeding BM predictions have evolved in the feet of *Tiliqua* and tails of some *Egernia* (Fig.5, S19–S23).

**Figure 5:**
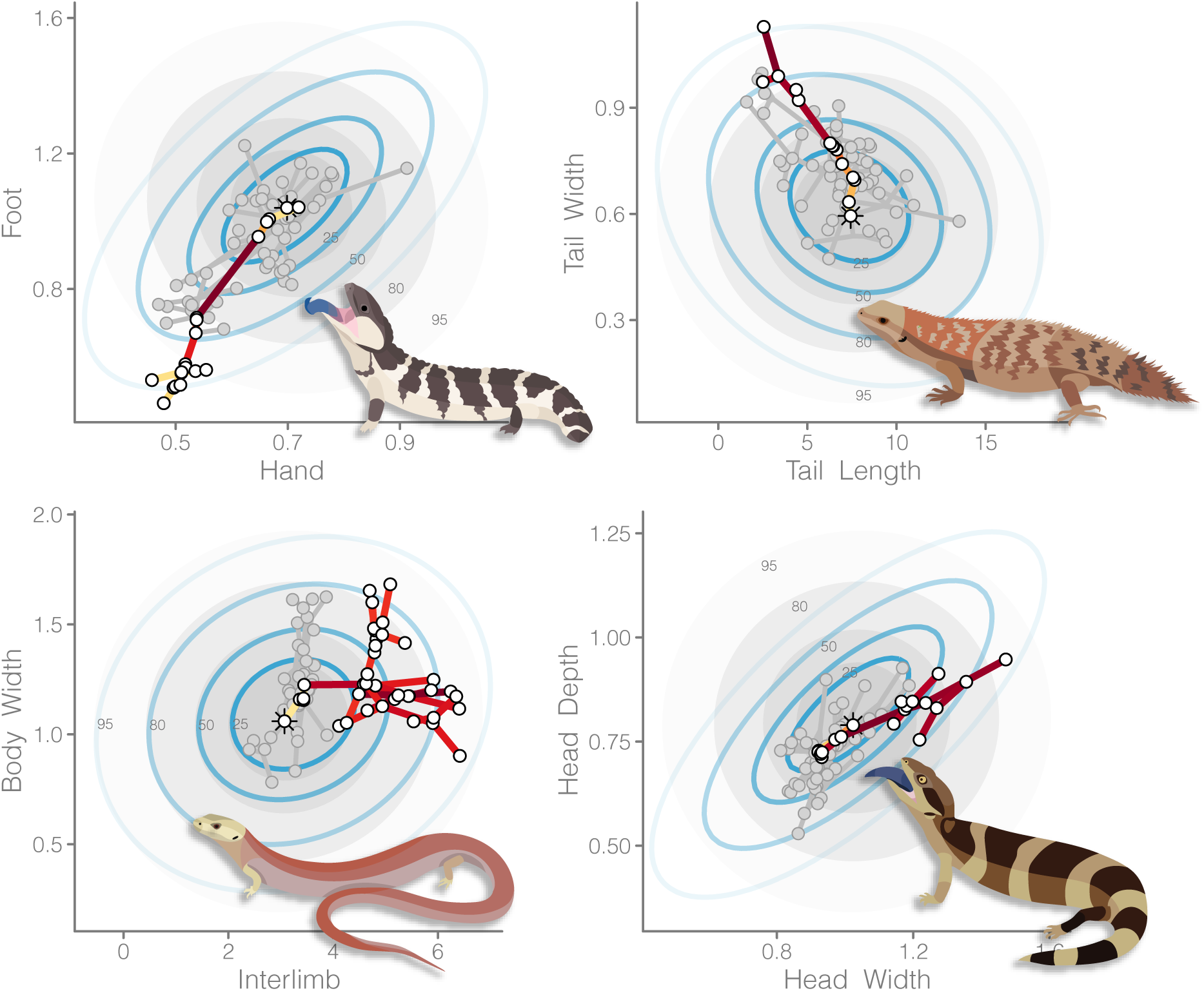
Novel morphologies exceed expectations of uncorrelated trait evolution. Bivariate plots show a phylomorphospace (species points connected by a phylogeny) of the Tiliquini. Each plot isolates a single clade within the group to highlight morphological extremes. Branches of the highlighted clade are colored according to evolutionary rates, with remaining species and branches in grey. In the background are ellipses containing 25/50/80/95% of traits simulated under empirical rates for uncorrelated (grey) and correlated (blue rings) Brownian Motion. Highlighted clades are (clockwise from top left: *Tiliqua*; *Egernia stokesii–E.hosmeri*, *Tiliqua rugosa–T.gigas*, *Cyclodomorphus–Tiliqua*).

**Figure 6.**
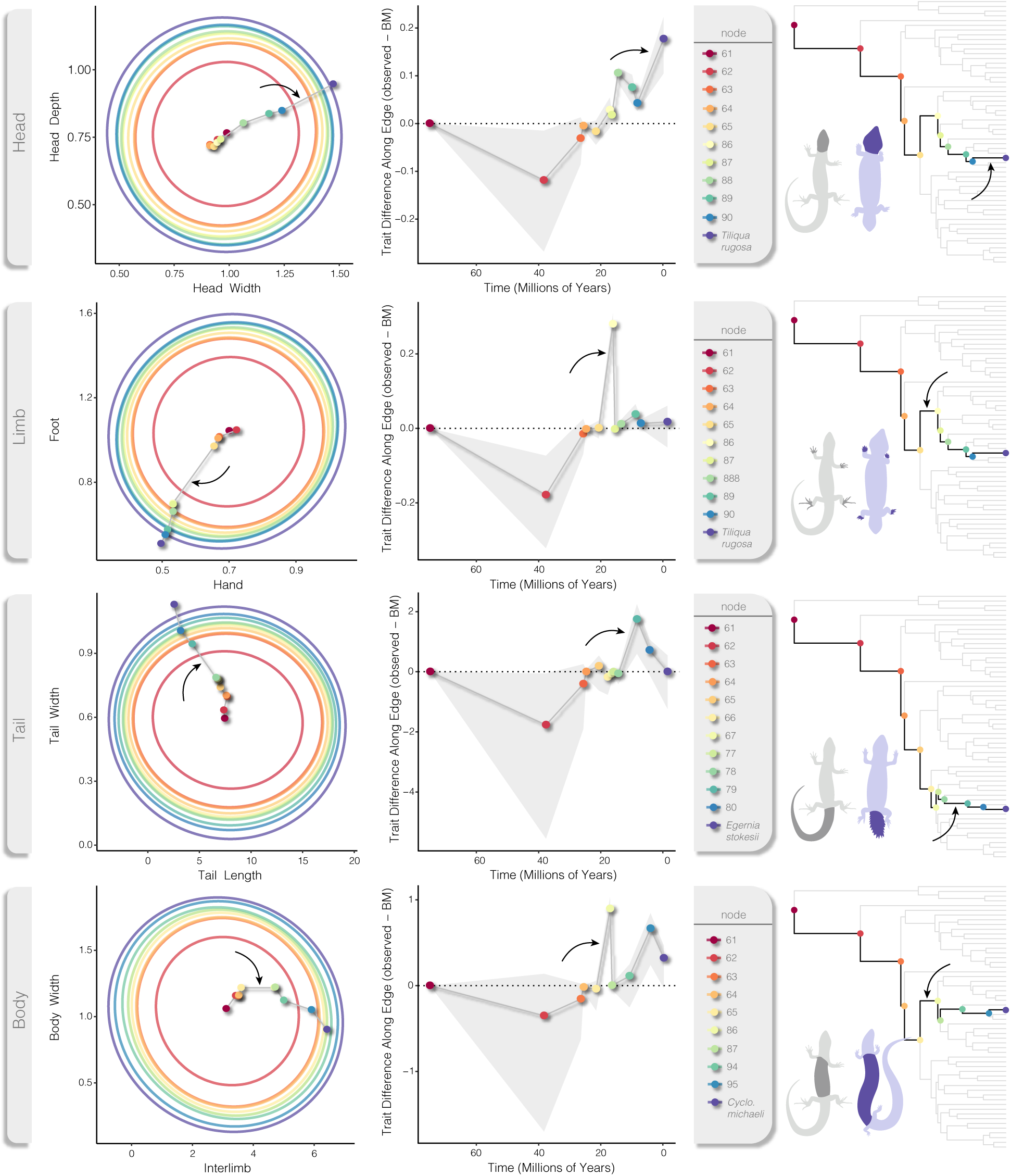
Phenotypic jumps are visible as greater than expected trait changes along branches, as estimated under the Variable Rates model and compared to Brownian Motion. Rows correspond to morphological modules, noted in grey at far left. Both circular (left column) and line plots (center column) show the evolutionary trajectory of focal traits from the root of the Tiliquini tree (node 57, dark red) to a single tip (dark purple) that has exhibited one or more rate bursts. (left) Large colored rings represent the simulated distribution of trait values under uncorrelated Brownian Motion at each node along the root-to-tip trajectory. Small colored points represent observed and ancestral trait values as estimated under Variable Rates as they traverse the path root-to-tip connected by a grey line. (center) The line plot indicates the trait difference of nodes as (observed - simulated) values with 95% quantiles shown in light grey. Nodes that fall above the dotted line (0) show greater than expected trait change along the branch that leads to that node, visible as greater distances between points in the plots at left. Nodes falling below the dotted line show less trait change than expected. Trees at right serve as a guide for the root-to-tip path of each example, with nodes colored accordingly. Examples of divergent morphologies are illustrated (purple) against a more ‘typical’ scincid body plan (*Liopholis*–grey). Black arrows in each row indicate the greatest jump in phenotype compared to expectations. References to node numbers can be seen in Fig. S28.

## Discussion

The variety of organismal forms provides a rich source of data for macroevolutionary biologists investigating the timing and accumulation of morphological diversity. Whether morphological disparity has accumulated early^1^, uniformly^34^, or via intermittent pulses^2,20,23^ remains a contested topic at varied evolutionary scales. Here we investigate the morphological evolution of the tiliquine skink body by looking across time, phylogeny, and body regions to better understand the modes by which traits evolve. We provide evidence for Tiliquini skinks that most traits evolve conservatively, but morphological novelty accumulates through extreme punctuations—jumps into new trait space. Our findings are consistent with evidence from the fossil record that major transitions to new phenotypes may occur over short intervals^14–16^. In Tiliquini skinks these jumps are uncommon (0.003–0.05 jumps/my), do not follow an established morphological order^35^, and can be nested to develop new trait combinations.

### Phylogenomics of the Tiliquini

Skinks are one of the most species-rich reptile groups, making up roughly 25% of all lizard diversity (1700+ species). The largest and most impressive skinks belong to the tribe Tiliquini which are primarily distributed across Australia with a small number of species found in Indonesia, Papua New Guinea, and the Solomon Islands. Despite their importance as models of reptilian sociality^29,36^ and recent fossil discoveries^31,37,38^, phylogenetic hypotheses for the Tiliquini have relied on a handful of molecular markers and limited species sampling^30,39,40^. Our exon capture dataset provides a well supported estimate of the relationships among all eight Tiliquini genera and 85% of described species. In agreement with previous estimates we recover the Tiliquini as members of the Lygosominae alongside the Sphenomorphini, Eugongylini, and Mabuyini (Fig.S1), suggesting an Asian origin for the subfamily^41^.

Living tiliquines are divided into three clades, comprising the enigmatic Crocodile Skinks *Tribolonotus*, the monotypic Solomon Islands endemic *Corucia*, and an Australian radiation. Splits among these groups are old (∼60 & ∼30 mya), and followed by the rapid Miocene divergence of all Australian Tiliquini genera (23–15 mya). These rapid speciation events result in short internal branches that prove difficult to resolve on a per-locus basis (Fig.1,S27). However, most of these difficult nodes are resolved by leveraging our ∼400 loci in ASTRAL, investigating summary statistics, and applying topology tests. Our new phylogenetic hypotheses of these nodes has necessitated that we propose taxonomic changes for two genera, which we provide in the attached Appendix (Taxonomic Implications and Changes). Some splits remain intractable however. The branching patterns among major *Egernia* clades, and the series of splits among *Lissolepis luctuosa*, *Liopholis*, and the remaining Australian tiliquines have splits so short (300–400ky) that they exist at the limits of phylogenetic reconstruction. Regardless of resolution of these difficult branches, the radiation of Australian Tiliquini into divergent ecologies and morphologies happened rapidly and in concert across open landscapes, closed forests, deserts, and mountain peaks.

### Evolving Novel Phenotypes by Bursts

Earth’s incredible biodiversity of forms and functions have evolved over hundreds of millions of years. Growing evidence suggests that much of this diversity accumulated via short periods of rapid phenotypic change, not by incremental divergence^23^. If this is true and pulsed morphological evolution is common, then it is important to understand how it contributes to the development of morphological novelty and diversity. To do this we identified how pulses are distributed through time, across the phylogeny, and among morphological axes. We provide evidence of highly heterogeneous patterns of morphological trait evolution in Tiliquini skinks, with truly novel phenotypes resulting from rapid episodic changes.

Much of modern macroevolutionary thinking relies on Simpson’s (1944) idea of adaptive zones. The varied adaptive landscape accumulates diversity as lineages traverse into new adaptive zones that are centered around fitness peaks. Lineages move into new zones by evolutionary jumps—rapid movement across suboptimal space. In comparative studies the idea of a multipeak morphological landscape is often described by multi-optima OU models^44–46^. While these models account for the clustering of species around discrete trait values (evolutionary ‘clumpiness’) they do not explain well the process of movement across the landscape to occupy new peaks. Our analysis of the tiliquine lizard body plan provides evidence that pulses in evolutionary rate and coincident phenotypic change are common across many morphological traits (preferred in 15 of 19 traits), but are likely rare events relative to slower background evolution. This suggests that large distances of trait space can be quickly traversed to reach new morphological realms, and these realms are likely wide. When comparing modeled empirical data to gradualist simulations, jumps are observable as large trait changes along individual branches (Fig.6). However, the accumulation of morphological disparity does not appear to be dictated solely by rare jumps, or to occur explicitly at speciation, as in the punctuated equlibrium model of Eldredge & Gould^47^. Instead, we find broad support for a model in which the background process of evolution by a random walk (Brownian Motion) is punctuated by bursts in evolution that result in jumps to new adaptive zones. We consider this process more akin to the “punctuated gradualism” model of Malmgren et al. (1983)^48^. While we recognize that our inferred process is not entirely consistent with the original description which proposed punctuations between evolutionary *stasis* and gradualism, we suggest that these distinctions may instead be indicative of similar processes occurring at different scales along the micro-to-macroevolutionary continuum. At both scales, a heterogeneous mode driven by an increase in evolutionary rates facilitates the evolution of novel and divergent morphologies (Fig.4)—in the case of these lizards, this occurs against a background morphological diffusion process. In this way we attempt to find common ground for both Darwinian and Simpsonian evolution.

In the Variable Rates model branch-specific shifts are indicative of an increase in evolutionary rate but remain a parameter of a random walk (Brownian) process. So, increasing the evolutionary rate suggests an increase in the *evolvability* and ultimately the potential variance of a lineage^49^. An alternative interpretation of our identification of rate bursts could instead be a directional (biased) trend towards distinct trait values. A recently designed method for addressing this—the *fabric* model of *BayesTraits*^50^—proposes that excess rate estimates could instead be absorbed by a biased walk to new trait space. Importantly, this does not fundamentally change the outcome that evolution has veered into a new morphological lane, but it does suggest a difference in the evolutionary mode. Whereas a rate pulse suggests a rapid, random exploration of trait space that lands in a new zone, directional trends elicit a guided evolutionary walk (e.g. via selection) towards a new area of trait space. Because a guided walk is directional the evolutionary rate need not be rapid. Regardless, the jump in trait value along a branch, and evolution of a new phenotype remains the same. This provides an appealing explanation for scenarios where selection might be particularly effective, such as in small populations or those with high trait variances.

Importantly, inferred phenotypic jumps happen along both the primary morphological axis—where they exaggerate existing variance (elaboration)—**and** along minor morphological axes—where they develop novel trait combinations (innovation). Along both major and minor axes relatively uncommon but major discontinuities result in unusual and novel morphologies, as seen in birds^51^ and fish^52^. In Tiliquini skinks, the most obviously novel phenotypes belong to the Bluetongue lizards *Cyclodomorphus* and *Tiliqua*. Bluetongues concurrently underwent dramatic shifts in limb and body modules, extending the length of the body and shortening the limbs. Subsequent jumps in head and tail modules resulted in further temporally staggered morphological bursts. These nested morphological pulses gave rise to broad heads and bodies, dwarfed extremities and stumpy tails in *Tiliqua*, highlighting that truly new morphologies can arise from rapid exploration of multiple body axes. However, some of the most dramatic changes have happened on the shortest timescales. In the *Cyclodomorphus michaeli* clade (*C.casuarinae*, *C.michaeli*, *C.praealtus*) which we estimate at less than 4 million years old, we identify jumps in tail, limb, and body traits. These result in the rapid shortening of the interlimb, tail, and leg lengths in *C.praealtus*, and lengthening of these traits in *C.casuarinae* and *C.michaeli*. Interestingly, these changes are potentially driven not by mechanistic morphological reasons but instead by physiological ones. *C.praealtus* lives at high elevations where shortened extremeties may be advantageous to maintain thermal mass following Allen’s Rule^53^.

Occasionally independent evolutionary trajectories arrive at similar adaptive zones, a process usually called morphological convergence. We identify convergence in the spiny crevice-dwelling clades of *Egernia* (*E. stokesii* and *E. depressa*). These clades have undergone rapid shortening and widening of the tail from non-armored, long-tailed ancestors (Fig.4). Amazingly, these clades have both arrived at this novel phenotype via rate increases of roughly ten times the background rate. The repeated evolution of a distinct phenotype like highly modified tails suggests strong selective pressures shaping some morphological axes^54^. Another example of the strength of selection concerns the only clade of primarily crepuscular and nocturnal tiliquines the *Liopholis kintorei*–*inornata* group. This clade is broadly distributed across arid and semi-arid Australia, living in burrows they dig in loose soil or sand. However, the transition of these skinks towards a nocturnal lifestyle does not appear to have driven exceptional morphological differentiation from other *Liopholis*, except for the size of their eyes. The transition in diel activity is coincident with a jump in eye diameter and even the evolution of vertical pupils in the Night skink *Liopholis striata* (Fig.S12). Despite the evolution of novel morphologies by phenotypic bursts, truly exceptional morphologies (those exceeding Brownian expectations) are rare. In a few cases, such as the feet of some *Tiliqua* and the tails of some *Egernia*, contemporary trait values fall outside what we expect under an uncorrelated random walk model. These instances provide the strongest cases for active selection driving the evolution of phenotypes (Fig.5).

### Conclusions

Animal bodies are made up of many morphological traits which in their myriad combinations contribute to Earth’s amazing biodiversity. Our study of gross lizard morphology provides evidence that trait evolution is heterogeneous, and is structured phylogenetically, temporally, and across the body-plan. This adds to a growing body of evidence that morphological traits diverge through evolutionarily discontinuous processes. These discontinuities contribute substantially towards the evolution of novel morphologies, and we suggest that this is may be a common process in morphological diversification across animals that helps reconcile Darwinian and Simpsonian evolution.

## Acknowledgments

A considerable thank you to the curators and staff of the many Australian museums and databases for access to tissues and locality data that made this work possible. We greatly appreciate the assistance of Emily Lemmon Moriarty, Alan Lemmon, and Michelle Kortyna in generating AHE data for this project. We thank Mark Hutchinson, Kailah Thorn, and Mike Lee for sharing their expertise in Tiliquini skinks, and colleagues in Natalie Cooper’s lab at the NHM for their insight. JSK, SCD, and DGC thank the Australian Research Council for ongoing support, including grants LP170100012 and FT200100108 to DGC. IGB is supported by a Marie Skłodowska-Curie Fellowship funded by the European Commission. We appreciate the constructive assessments of three anonymous reviewers who challenged us to improve this research.

## Author Contributions

Conceptualization, IGB, JSK, SD; sampling, JSK, SD.; analysis, IGB; funding acquisition/resources, JSK, SD, DCG; data curation, DCG, IGB; writing—original draft, IGB; writing—reviewing & editing, all authors.

## Competing Interests

The authors recognize no conflicts of interest, either direct or indirect, that might bias the conclusions, implications, or opinions stated in this work.

## Materials and Methods

### Data and code availability

All data and code necessary to replicate the results and figures of this study are available on GitHub at www.github.com/IanGBrennan/Tiliquini.

### Taxon Sampling

We assembled an exon-capture dataset across 77 Tiliquini skinks representing 53 of 62 currently recognized species, with a focus on Australian taxa (47 of 48 spp.). This sampling covers all 8 genera, as well as many recognized subspecies (Table S1). We included outgroup representatives from all subfamilies and most tribes, generated as part of broader look at squamate phylogenetics^41^.

### Molecular Data Collection and Processing

Genomic loci were targeted and sequenced using the Anchored Hybrid Enrichment approach^55^, and resulted in 379 loci (average coverage 367 loci, min = 362, max = 375) totaling ∼600 kbp per sample (Fig.S3). We downloaded the complete CDS of *Anolis carolinensis* and used a reciprocal blast to identify unique exonic targets in our AHE dataset (n=358). We identified 25 targets as separate exons of shared genes (APOB, DSP, DST, FAT4, HIVEP2, IRS1, KAT6B, PLXNA1, RBM15, SHANK2, USF3, ZNF536). We then downloaded additional squamate genomes as well as *Sphenodon* and *Gallus* and extracted orthologous targets. Rough alignments were compiled using MAFFT^56^ and refined into correct reading frame using MACSE^57^, alignment parameters are specified below. Per locus information content is summarized in .

Input MAFFT/MUSCLE alignments were refined with MACSE using the following call:

~~~
$ java -jar /Applications/macse_v2.03.jar \
 -prog refineAlignment -align [rough alignment path] \
 -optim 1 -local_realign_init 0.1 -local_realign_dec 0.1 \
 -fs 10 -stop 10
~~~

Additional samples were added to MACSE alignments using MAFFT:

~~~
$ /Applications/mafft-mac/mafft.bat \
 --quiet --add <input_alignment> \
 --keeplength --reorder <existing_alignment> \
 > <combined_alignment>
~~~

Valuable statistics of our molecular data are summarized in supplementary files *Tiliquini_Summary_PerLocus.txt*, *Tiliquini_Summary_PerSample.csv* and Fig.S4. For brevity, we provide some basic statistics below:

- Average alignment length was 1884 bp (min=332; max=2708).
- Average site coverage was 1837 (+/-930).
- Average number of reads per sample was 6,465,416 (min=340,302; max=18,867,860)
- Average percent missing data per locus was 8% (min=0.93%; max=19%)

### Morphological Data Collection

We collected 21 linear measurements for 61 Tiliquini skink species from 650 museum samples (average 10 per spp., min.=2, max.=134). These linear measurements aimed to capture the gross morphology of the lizard body plan and are distributed across the head, body, limbs, and tail (Fig.S5). From the initial 21 measurements some were further divided (e.g. snout-vent length) or dropped to arrive at a set of 19 non-overlapping morphological traits. Definitions of trait measurements and adjustments are summarized in Table S3 and Table S4.

### Phylogenetic Analyses

We reconstructed individual genealogies for our exon-capture data (n=379) under maximumlikelihood in IQ-TREE 2^58^, allowing the program to assign the best fitting substitution model using ModelFinder^59^, then perform 1,000 ultrafast bootstraps^60^. We then estimated the species tree using the shortcut coalescent method ASTRAL III^61^, with IQ-TREE gene trees as input. For comparison we estimated a locus-partitioned concatenated species tree in IQ-TREE and investigated differing species tree topologies by delimiting the anomaly zone. To assess the likelihood that discordant branches fall into the anomaly zone, where concatenation may be positively misled, we identified anomalous branches using *Anomaly Finder* ^32^.

To estimate divergence times among taxa we applied a series of fossil and secondary calibrations in MCMCtree^62^ and as outlined in Table S2 and informed by a combined-evidence morphology and molecular dating estimation in BEAST^63^.

We estimated individual genetrees (n=379) under maximum-likelihood in IQ-TREE 2^58^, allowing the program to assign the best fitting substitution model using ModelFinder^59^, then perform 1,000 ultrafast bootstraps^60^. We then estimated the species tree using the shortcut coalescent method ASTRAL III^61^, with IQ-TREE gene trees as input.

For each exonic locus we started by estimating raw genetic distances from alignments to use as a proxy for evolutionary rate. We removed the fastest 5% and slowest 5% of loci to avoid issues with extreme rate heterogeneity then partitioned the remaining loci into three partitions using AMAS^64^ and removed third codon positions.

~~~
python3 /Applications/AMAS-master/amas/AMAS.py concat
-d dna -f fasta -t GeneClump1_CP12.fasta -n 12 -i …
-u fasta -c 8 -t GeneClump1_CP12.fasta -p GeneClump1_CP12_parts.txt
~~~

### BEAST: Divergence Dating

To incorporate fossil information in a cohesive way we modified the combined-evidence BEAST xml file of Thorn et al.^38^ to include our phylogenomic data and additional fossil taxa. BEAST analyses are often model and parameter rich, so we outline the major assumptions and methodological decisions below, and direct the reader to the XML file for complete annotations of the model, priors, data, and settings.

We started by replacing the existing multilocus mitonuclear molecular partitions with our three AHE partitions described above. Due to the computational demands of full Bayesian methods such as BEAST we were forced to downsample the molecular matrices to allow appropriate MCMC sampling and mixing. The complexity of BEAST analyses also meant we chose not to expand the species sampling dramatically. This helped us to avoid creating a gappy morphological matrix and tree reconstruction error resulting from concatenation as shown in section Anomaly Zone, and allowed more direct comparison with the results of Thorn et al. (2021). We randomly retained 50,000 sites for each of the three partitions and applied a GTR model to each. We removed the continuous morphological data which had shown to cause spurious results^38^ and added in the newly redescribed Plio/Pleistocene species *Tiliqua frangens* to the discrete morphological data. To the discrete morphological data we fit the Mkv model with correction for non sampled constant characters. Polymorphic discrete morphological data were treated as coded (e.g. (0,1)), rather than as unknown (?). We applied separate uncorrelated relaxed clocks for the molecular and discrete morphological datasets with wide lognormal priors (*x̄*=0.001, 0.01; *sd*=0.33) based on exploratory analyses. Fossil tips were dated using a tip-dating strategy informed by their stratigraphic information and the morphological data, which in turn informed node ages in conjunction with the molecular clock model. These elements were combined under the birth-death serial sampling (BDSS) model of Stadler^65^. While there will always be debate regarding model and tree prior choice, we feel strongly that the tip-dating methodology removes subjectivity regarding node calibrations and more completely uses fossil information. Additionally, the BDSS model allows for a mechanistic explanation of tree characteristics resulting from parameterized speciation, extinction, and sampling processes^66^.

We ran four independent runs for 100 million generations, inspected log files for convergence and stationarity in TRACER, combined runs with LOGCOMBINER, and generated a maximum clade credibility tree with median node heights using TREEANNOTATOR (Fig.S7). Phylogenetic analyses of Tiliquini skinks based on morphology alone can generate unreliable topologies^31^ but are buffered by the inclusion of molecular data. Our combined-evidence tree is mostly consistent with our molecular tree, but molecular and morphological partitions exhibit some differences in behavior. The morphological partition in particular appears to show extreme rate variation among taxa, particularly along the lineage leading to *Cyclodomorphus* and *Tiliqua* (Fig.S8). We anticipate that this rate variation may inflate estimated ages, and as such used the estimated 95% confidence intervals to generate wide Skew-T priors for our subsequent MCMCTree analysis.

### MCMCTree: Divergence Dating

For each partition we ran MCMCTree with usedata = 3 to get the approximate likelihoods and branch lengths using *baseml*^67^, then concatenated the out.BV files together. We then ran four replicate *mcmctree* analyses on the gradient and Hessian (in.BV file; usedata = 2), each for 20k burnin generations before collecting 20k samples at a sampling frequency of 100 generations (2,020,000 total generations). We compared mcmc files for stationarity and convergence (ESS of all parameters > 300), combined them using logCombiner, and used this combined mcmc file to summarize divergence times on our tree (*print = -1* in .ctl file). To investigate our priors we ran an additional analysis usedata = 0 to run explicitly from the prior calibrations and determine our effective priors for comparison against our posterior age estimates. We then plotted the applied priors (Table S2), against effective priors (priors as a result of multiple interacting priors from usedata = 0), and posterior estimates to ensure the appropriate behavior of the MCMCtree analyses (Fig.S9). Our summarized MCMCtree output is show in Figure S1.

To take advantage of all available morphological and phylogenetic information we incorporated two additional *Tribolonotus* species into our dated tree used for trait evolution analyses. *T. blanchardi* and *T. ponceleti* were added to our MCMCTree output using *bind.tip* in *phytools*, based on the topology and ages estimated from a recent phylogenomic investigation of Squamata by Title & Singhal et al. (2024).

### Phenotypic Analyses

The below methodology is accompanied by a series of scripts found in the associated GitHub repository /Scripts which will run on the included/Data files. These steps can be followed chronologically and scripts are indicated as such `<Number>_<Process>.Re.g. 00_Data_Preparation.R.`

#### Defining Modules

Our interest is in identifying the tempo and mode of evolution that produces morphological novelty and so we focus on the dynamics of morphological diversification in the Tiliquini radiation. Our approach is based on a novel dataset that summarizes the lizard body plan. We started by generating mean trait values per species (Scripts/00_Data_Preparation.R). To remove the effect of size on individual traits (allometry) we then calculated the geometric mean of all traits by species and used this to transform trait measurements into log-shape ratios. This also provided an additional trait *size*. To identify independently evolving morphological modules we designed *a priori* hypotheses as model inputs for the package *EMMLi*^33^. We provided six general hypotheses for model comparison with the most specialized model allowing traits of the head, limbs, body, and tail to evolve as independent modules, and the most restrictive null model lumping all traits into a single module. (1) a four-module model (m1:*Body_Tail_Limbs_Head*) in which each of the major body regions is isolated as an independent module; (2) a threemodule model (m2:*BodyHead_Tail_Limbs*) where traits of the head and body are combined into a single module; (3) a two-module model (m3:*BodyHeadTail_Limbs*) in which traits of the head, body, and tail are combined into a single module; (4) a three-module model (*Body_Head_TailLimbs*) where tail and limb traits make up a single module; and (5) a two-module model (m4:*BodyHead_TailLimbs*) where body and head traits are one module and tail and limb traits another. We compared these to one another and to a null model where all traits exist as part of a single module (m0). For model specifications shown see Table S5 and Fig.S11. *EMMLi* also allowed us to compare models in which the correlation coefficient among modules and among traits within modules is either similar or different (see Goswami & Finarelli Fig.2). Once we established the preferred model (head, body, limbs, tail as separate modules with differing inter– and intramodule correlations) we split our traits into module-specific datasets.

#### Comparative Trait Evolution

To identify the evolutionary mode of individual morphological traits we fit a series of models ranging from a basic unbiased random walk to entirely punctuated. These allowed phenotypes to evolve through incremental change (Brownian Motion—BM), incremental change around an optimum (Ornstein Uhlenbeck), decreasing change with time (Early Burst)—akin to an adaptive radiation scenario, two “pulse” or “jump” models which allow change in instantaneous bursts (Jump Normal, Normal Inverse Gaussian), and a Variable Rates model in which rates of change vary across individual branches of the tree. In principle the Variable Rates model is a multirate BM model conceptually similar to the uncorrelated relaxed clock model of timetree estimation. The number of different evolutionary rates (*sigma*) can vary from one (constant rate) to *n*—where *n* is the number of branches in the tree—and is determined by the reversible jump MCMC algorithm. For consistency we fit the BM, OU, EB, JN, and NIG models in *pulsR*^23^. We fit the VR model in *BayesTraits V4* ^68^ and processed the output using the standalone *PPPostProcess* software. We compared model fit by AIC scores. *BayesTraits* implements an MCMC algorithm so to estimate a maximum likelihood for the VR model we transformed the input tree by the estimated median rate scalar, then fit the observed data to the transformed tree using BM in *pulsR*. We obtained an AIC value by penalizing the likelihood for each scaled branch of the transformed tree with mean scalar *r* greater than two, in addition to the estimation of the rate parameter and root state. We then calculated AICw for each model and identified a preferred model if its AICw was greater than twice the next best model.

We fit the Variable Rates model via the comparative methods software *BayesTraits V4* which implements a reversible-jump MCMC sampler. We began by fitting the VR model (burnin 10 million generations, 110 million total generations) to each individual trait and summarizing the output to check for convergence with *Scripts/processBTVarRates.R* verifying that all parameters had reached effective sample sizes (ESS) >200. We summarized model parameters and used these to provide informed priors (*Data/BayesTraits/…*) for full analyses of individual modules (multi-trait). To verify model fit and consistency among runs we ran four separate runs for each module (burnin 100 million generations, 200 million total generations). Individual model outputs *. . . VarRates.txt* were summarized using the *PPPost-Processor* script. We then summarized runs for each module accepting shifts that occurred in >70% of the sampled posterior in **all four** runs (01_Process_BayesTraits_Output.R). Trees were transformed, colored, and plotted with evolutionary rate and shift information using custom scripts (02_VarRates_Trees.R, see Figure 4 left column)

#### The Influence of Fossil Information

To investigate the effect fossil taxa may have on our evolutionary inferences we analyzed a dataset of head length of extinct and extant Tiliquini species by fitting the Variable Rates model. We used the age and relationships of extinct species as inferred in our BEAST analysis and added these taxa to our MCMCTree timetree. Most fossil tiliquines are highly fragmentary and limited to skull elements, so we focused on the trait “head length” which in extant taxa was measured from the nose tip to the anterior of the ear opening, and for extinct taxa was measured as the full extent of the mandible length. These measurements are comparable in this group of skinks. Results are qualitatively identical with the same number and positions of shifts (Fig.S13).

#### Temporal Trends in Trait Evolution

To understand the temporal and phylogenetic heterogeneity of evolution we estimated ancestral states for each trait under the rate heterogeneous VR model by optimizing Brownian Motion on the VR rate-transformed trees in *phytools*^69^. We transformed branch lengths according to the mean rate scalar estimated by BayesTraits (*mean.scalar.trees*), using plotting_BayesTraits.R.We then extrapolated trait values linearly along branches given start and end values at nodes and a constant evolutionary rate. We did this from the root to the tips in 0.1 million year windows across all branches 03_Extract_Ancestral_Traits.R.

To summarize the standing morphological variation across the same temporal windows we calculated disparity as both the variance and the average squared Euclidean distance among all pairs of contemporaneous taxa 04_Disparity_Through_Time_TRAITS.R. Similarly we extracted the mean evolutionary rate in 0.1 million year windows for each trait and module 05_Rate_Through_Time_TRAITS.R. To determine if our observed patterns follow a null (BM) expectation of the accumulation of disparity through time we simulated univariate and uncorrelated and correlated multivariate datasets for each trait and module applying parameter estimates from observed data for theta, sigma, and covariance. We carried out the same disparity and rate through time extraction methods in 0.1 million year windows. To understand the relative contribution of niche expansion and niche packing to the accumulation of disparity through time we compared slopes of the accumulation of variance of observed traits and modules to simulated data. We then plotted the trends in variance, rate, and slopes 06_Disparity_Rate_Through_Time_MODULES.R.

We further developed a series of visualizations to assist in understanding the timing, frequency, distribution, and influence of inferred morphological rate shifts. To visualize the relationship between shifts and their morphological consequences we built Traitgrams (Figure 4, right column) with edges colored according to their evolutionary rate 07_Rate_Trajectory_Traitgram.R. To highlight the magnitude of these shifts relative to the background evolutionary rate we developed Rategrams (Figure 4, center column) which also serve to show the temporal separation and clustering of shifts 08_Rate_to_Node.R. To further compare the paths of empirical and Brownian evolutionary change and morphospace expansion we plotted bivariate trait change from the root to a specified tip 09_Sim_to_Node.R. In order to visualize the accumulation of trait variation under Brownian Motion we simulated 500 datasets per trait with parameters informed by empirical model fits, then plotted an empirical path atop this. These plots highlight the relatively even accumulation of variation under BM, whereas great trait changes are often partitioned onto individual edges in our preferred heterogeneous model (Figure 6, left column). To emphasize the discrepancy of trait change along individual edges, we plotted the difference in empirical and simulated trait change occurring between ancestor and descendant nodes (Figure 6, center column) 10_Distance_Btwn_Nodes.R. These plots illustrate that while trait change along many branches can be explained by a global background constant-rate Brownian process, some branches depart significantly from these expectations.

#### Innovation and Elaboration

To summarize the major avenues of morphological change we started with our dataset of all traits including estimated trait values at ancestral nodes which were inferred using our rate-transformed BayesTraits trees. We then ran PCA on (1) all traits jointly across all species and across (2) individual modules and (3) clades separately, then fit linear models to the first 2 PC axes (always accounting for *≥* 90% of variance). This allowed us to identify the major axes of elaboration (PC1) and innovation (PC2) following the language of Endler^12^ and Guillerme et al.^13^ 11_Innovate_Elaborate.R. We then identify change between parent and child node pairs as primarily elaborative or innovative by estimating the angle/slope of change of the line connecting them in Euclidean space (Fig.S16) and indicated by color. For branches which exhibit both elaborative and innovative change (likely most branches exhibit some combination of both), the primary trend is a discrete binary state described by the angle of change (innovation: >45*^◦^*<135*^◦^*, >225*^◦^*<315*^◦^*; elaboration: >135*^◦^*<225*^◦^*, >315<360*^◦^*). Further, we identify the strength of this change as the distance between the points and indicated by color saturation. Adjusting the threshold for distinguishing between elaboration/innovation (e.g. to a more conservative coding as in Fig.S16), does not qualitatively change the inferences of periods of primarily elaborative or innovative change.

## Supplementary Material

Supplementary material included below consists of additional figures, tables, and extended methods to complement the main text.

**Table S1.**
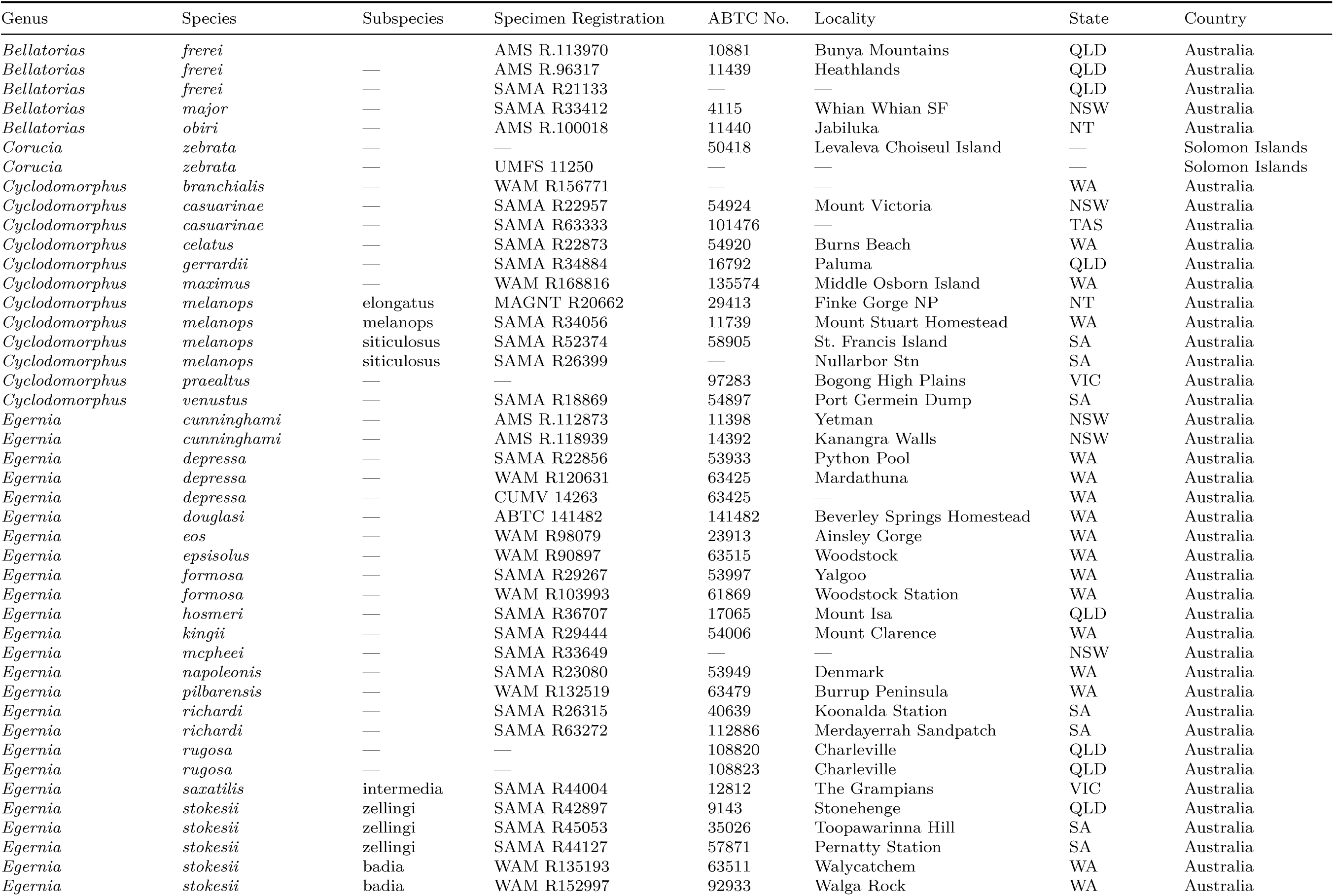
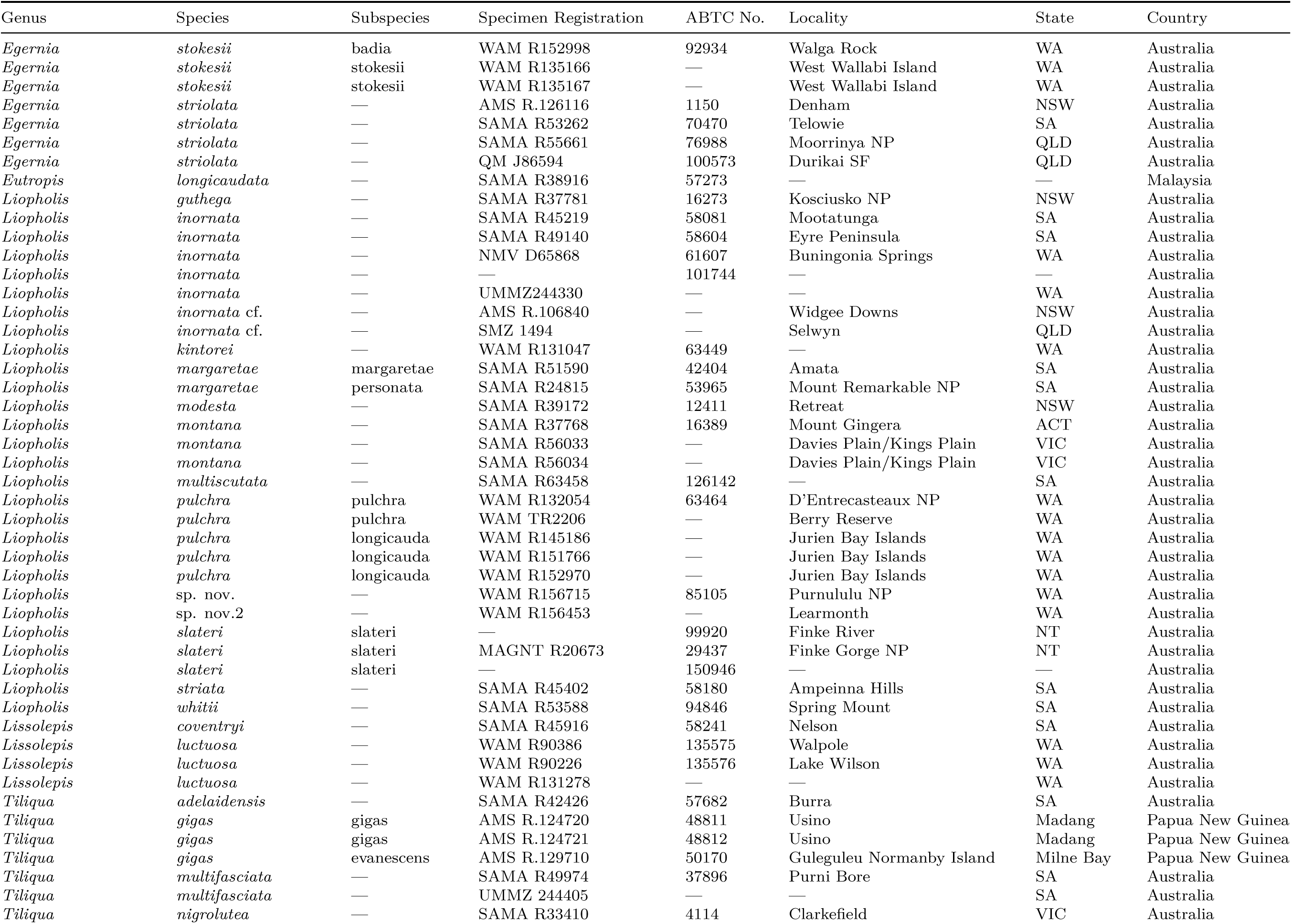
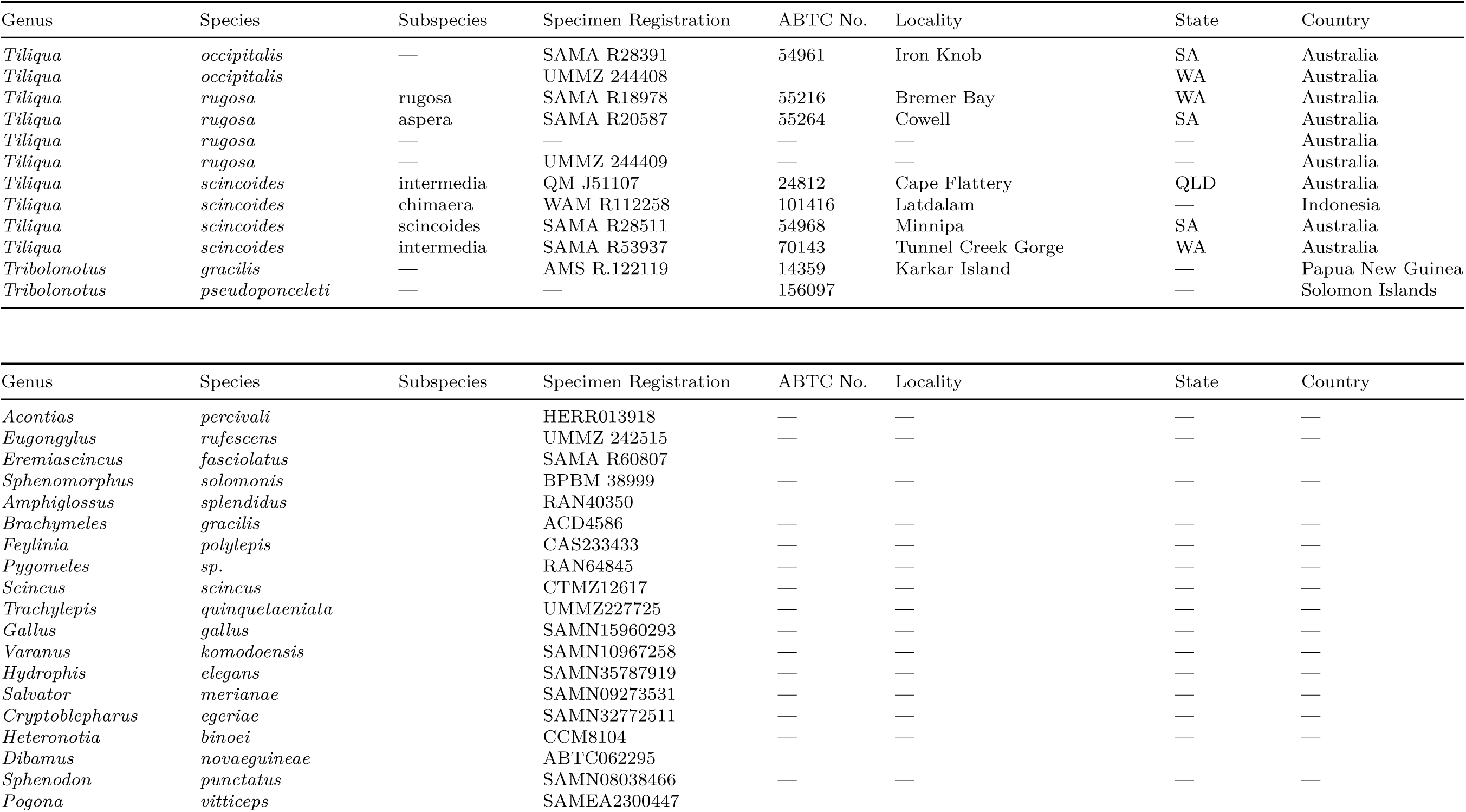
Taxon sampling: Ingroup Tiliquini sampling is listed in upper table, with outgroup sampling following.

**Table S2.**
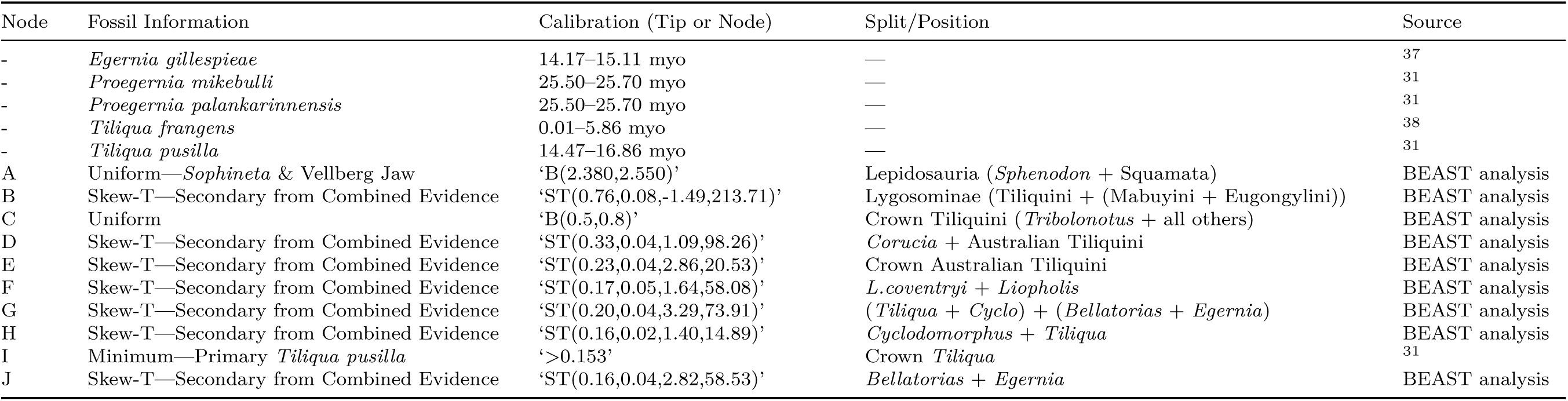
Fossil ages and divergence dating calibrations as implemented in BEAST and MCMCtree. The root divergence between lepidosaurs (*Sphenodon* + squamates) is based on the fossil taxa *Sophineta cracoviensis* (Evans & Bialynicka, 2009), *Megachirella wachtleri* (Renesto & Posenato, 2003), and the Vellberg Jaw (Jones et al., 2013). These taxa represent stem squamates and rhynchocephalians, and so provide a soft lower bound on the crown divergence of Lepidosauria, with a soft upper bound provided by *Protosaurus speneri* (von Meyer, 1832). Fossil *Egernia*, *Proegernia*, and *Tiliqua* samples were implemented as tip calibrations in the combined evidence BEAST analysis shown in Fig.S7. Node calibrations A–J implemented in our MCMCtree analysis are shown in Fig.S1. All node priors as applied in MCMCtree are soft, allowing estimated ages to pull beyond priors if driven by the data.

**Figure S1:**
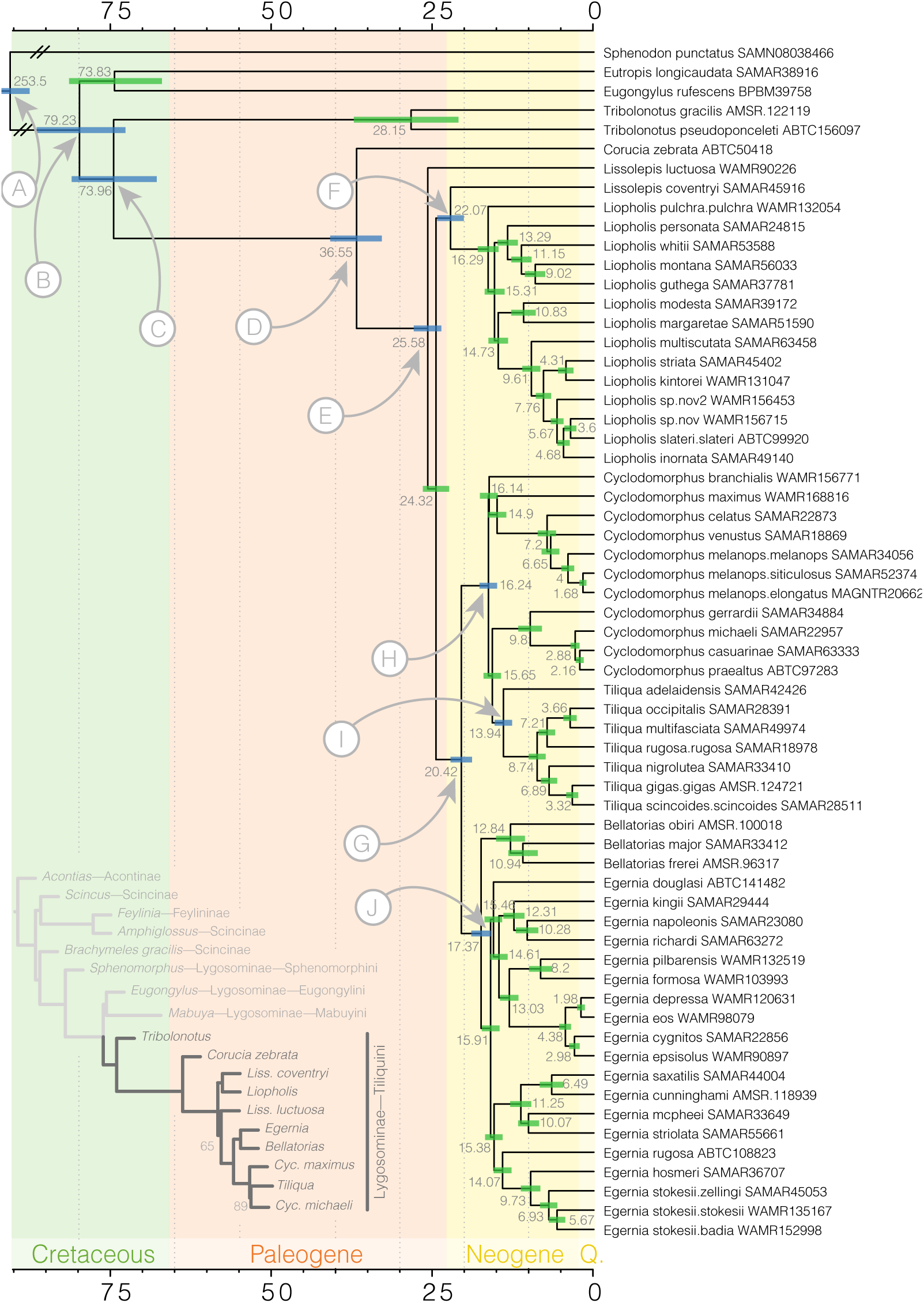
Reduced sampling Tiliquini species tree estimated with ASTRAL from IQTREE2 genetrees and time-calibrated with MCMCtree. Shaded bars at nodes indicate 95% confidence estimates on ages. Nodes labelled by letters A–F correspond to fossil calibrations listed in Table S2. Inset tree depicts the scincid phylogeny estimated from AHE data showing the placement of the Tiliquini among the Lygosominae.

**Figure S2:**
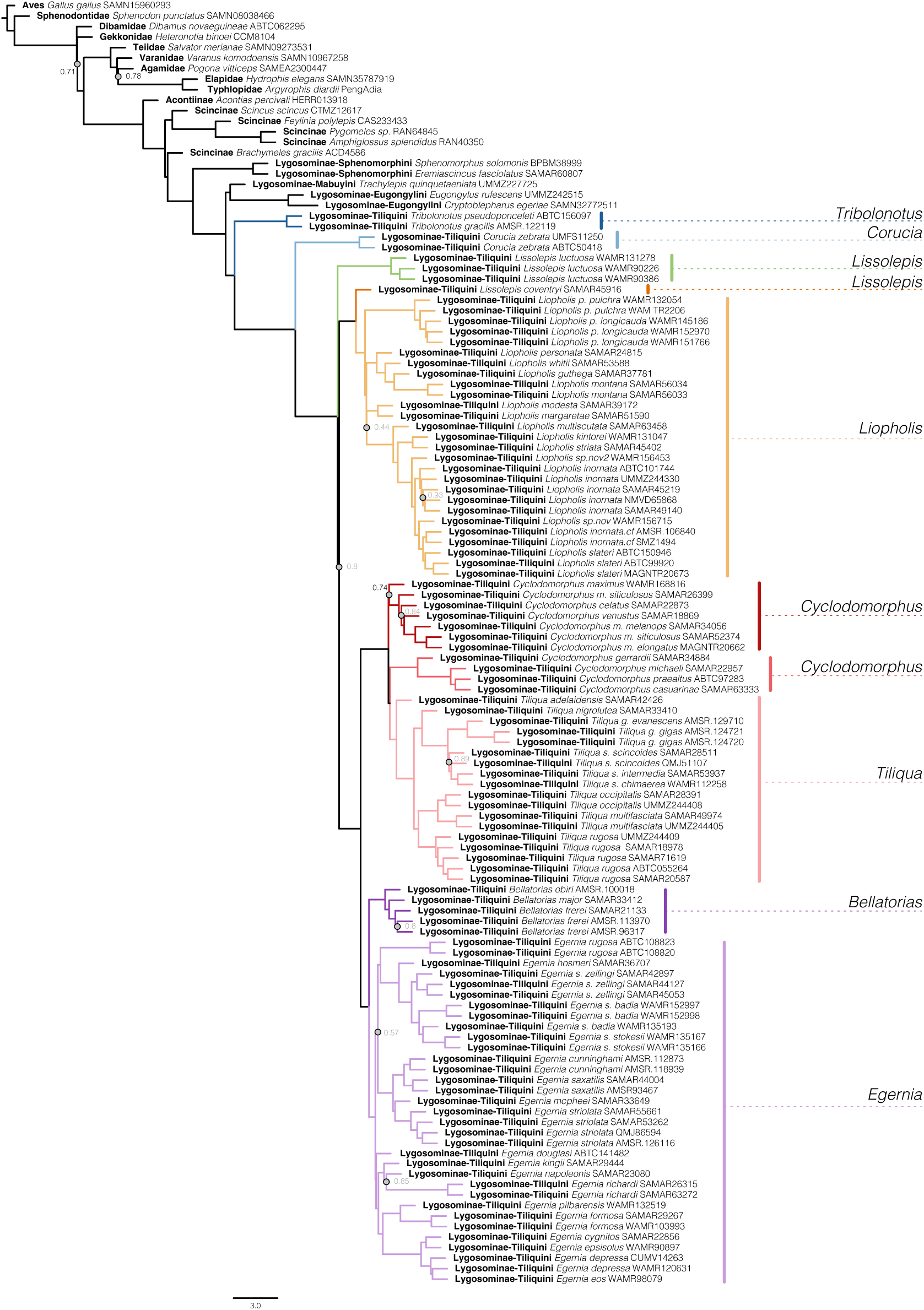
Fully sampled Tiliquini species tree estimated with hASTRAL from 379 IQTREE2 genetrees. Presentation of this full tree highlights intraspecific sampling. Support values (local posterior probabilities) are 1 for all nodes except those marked by a grey circle and text.

### Alignment Specifications

#### Investigating Data Completeness and Informativeness

**Figure S3:**
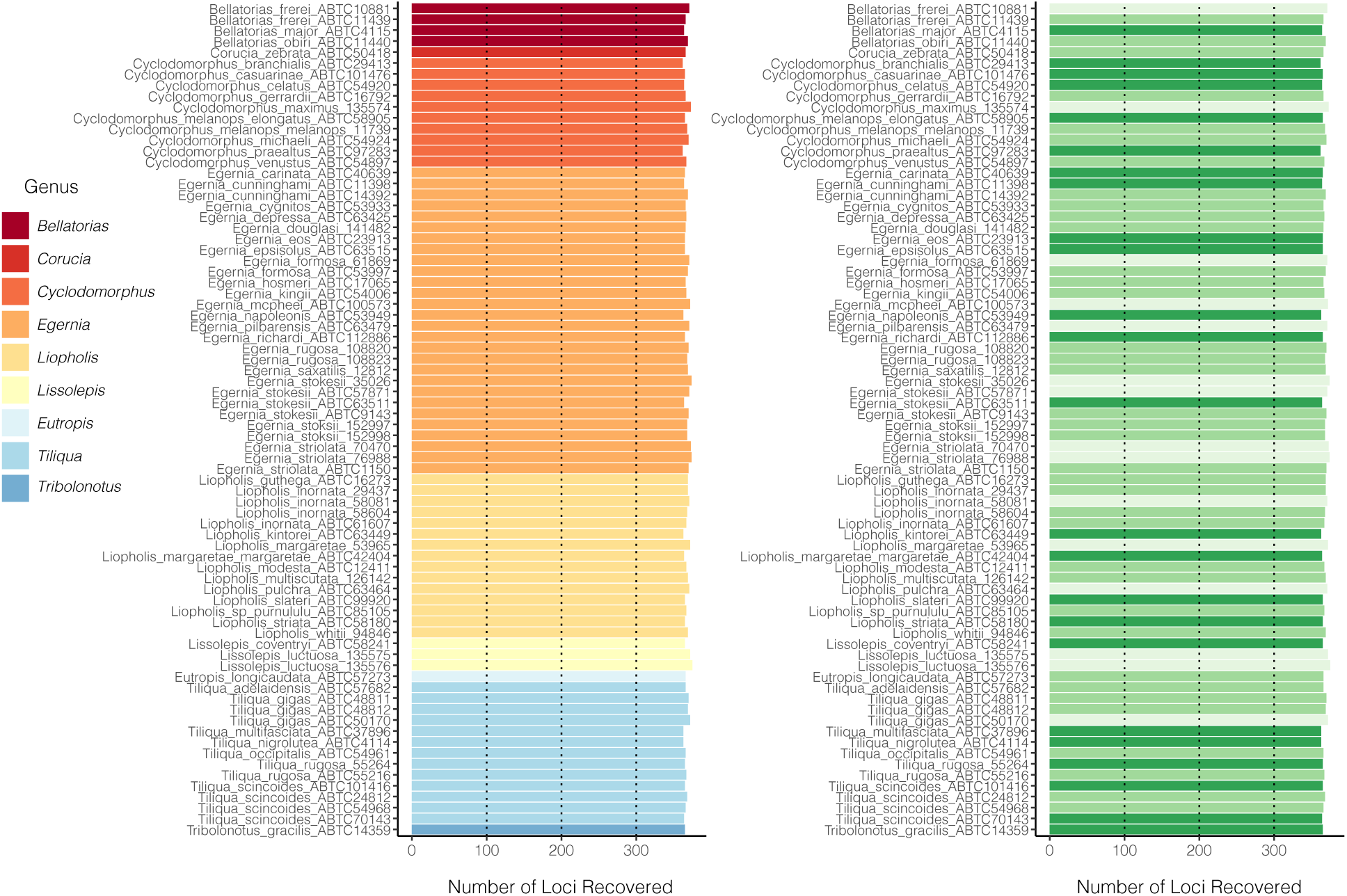
Number of loci recovered per sample for all Tiliquini and outgroup taxa included in the molecular data. Samples are colored by Genus (left) and coverage (right).

**Figure S4:**
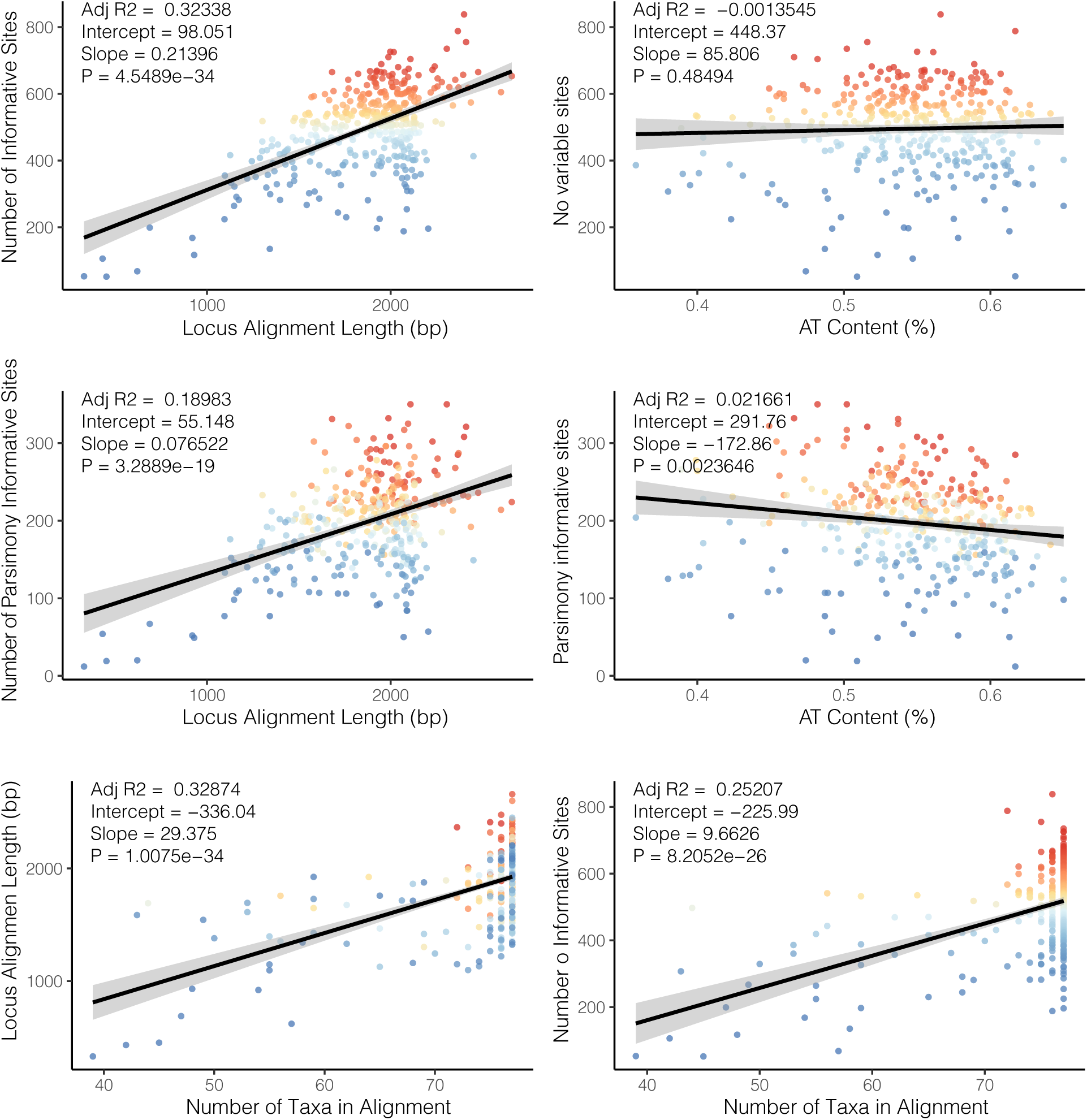
Plots of individual locus completeness and informativeness. Points are colored according to number of informative (variable) sites (blue—few, red—many). Top row shows the number of variable sites in each alignment as a function of alignment length and AT content. The middle row shows the number of parsimony informative sites as a function of alignment length and AT content. The bottom row shows alignment length and number of variable sites as a function of completeness.

### Morphological Measurements

**Figure S5:**
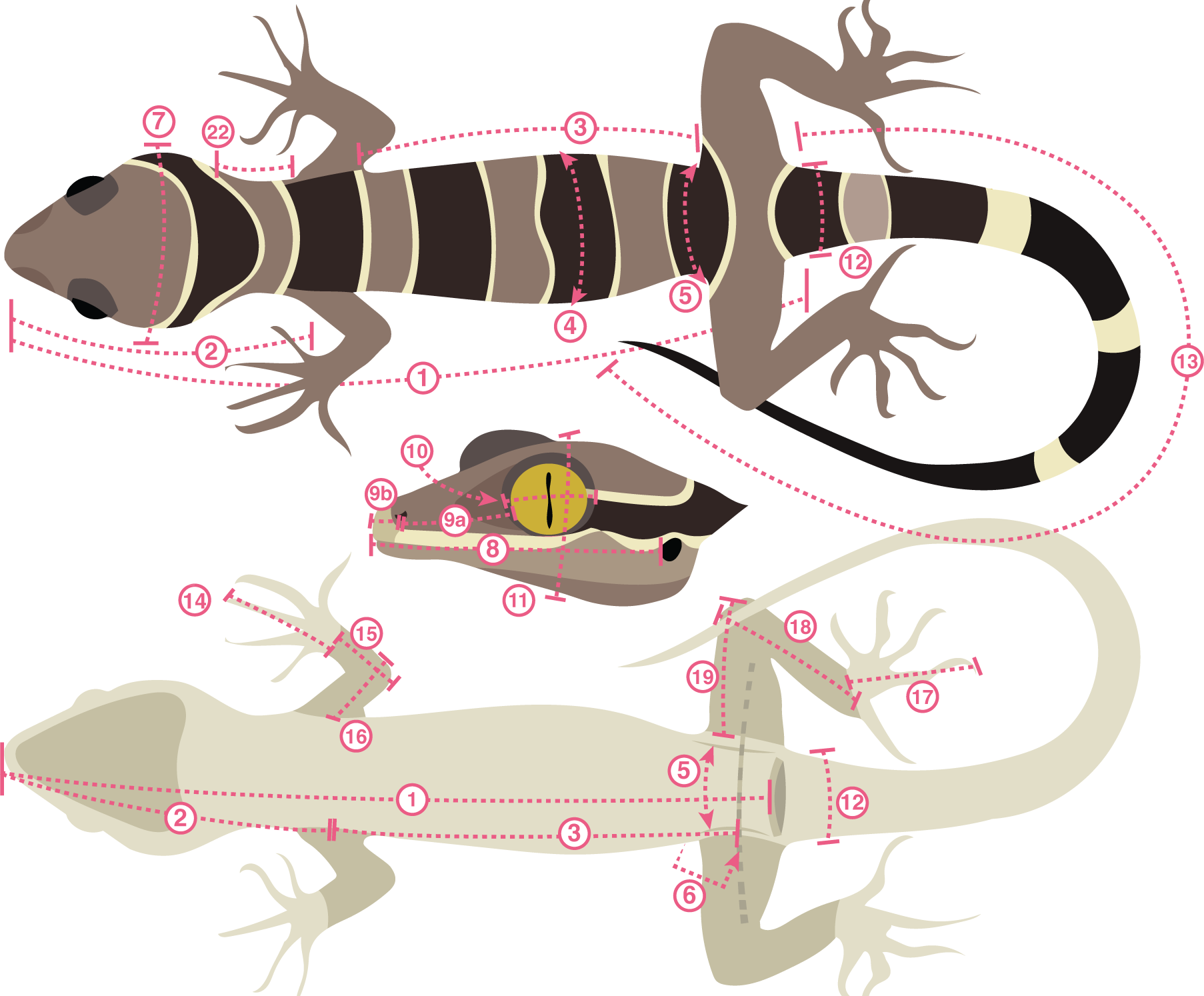
The 21 linear measurements collected.

**Table S3.**
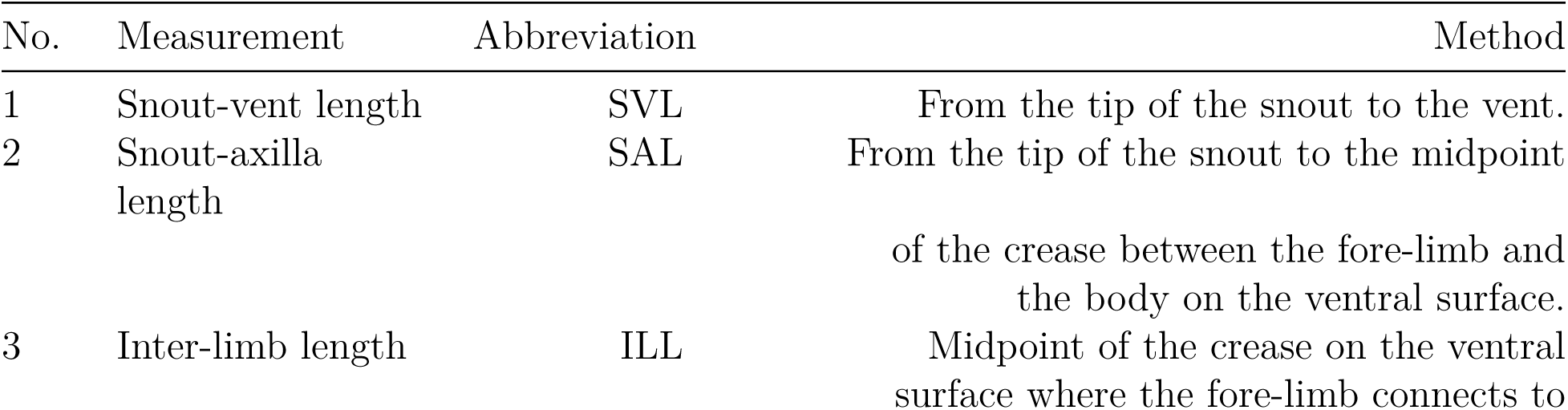
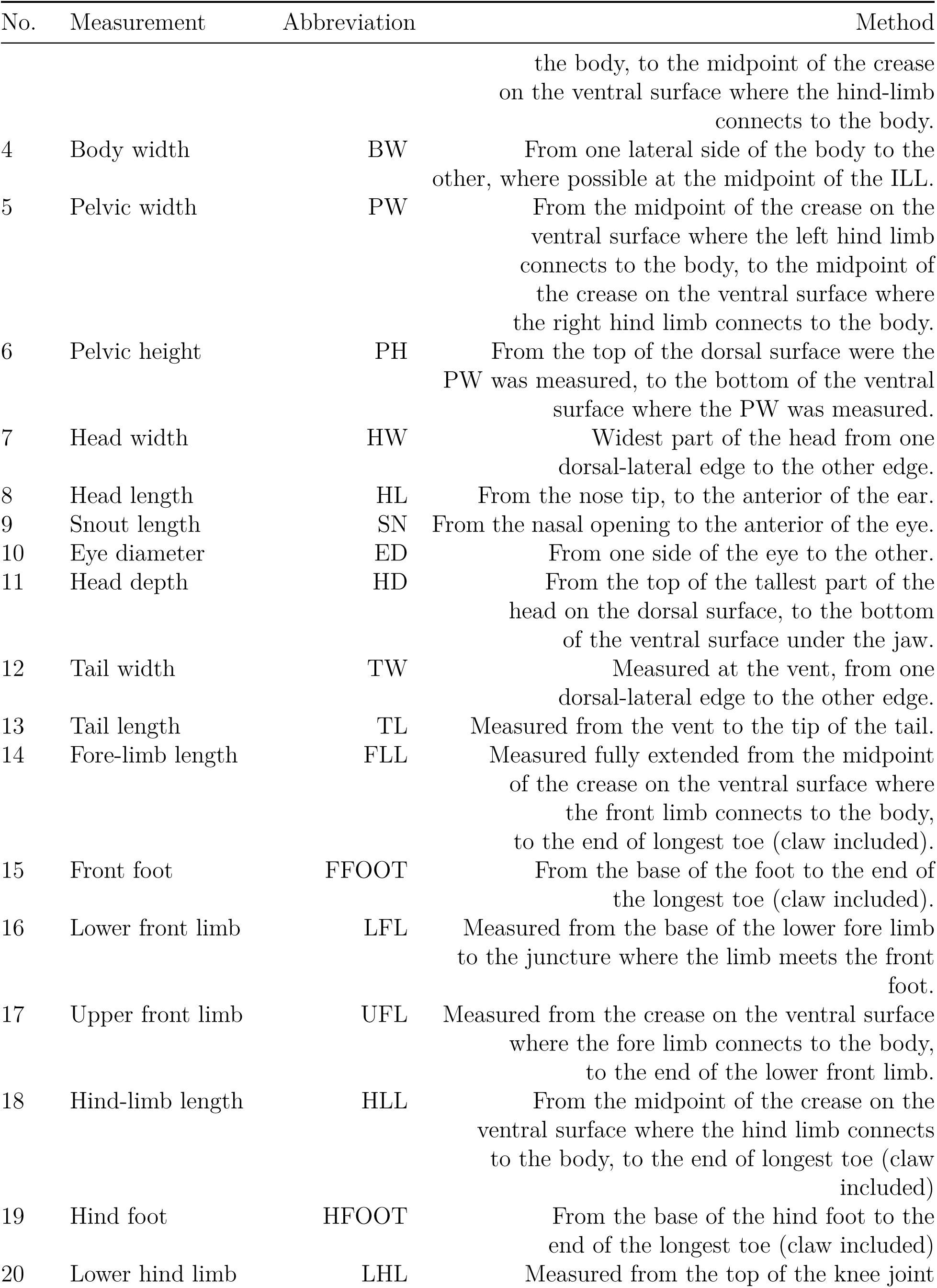
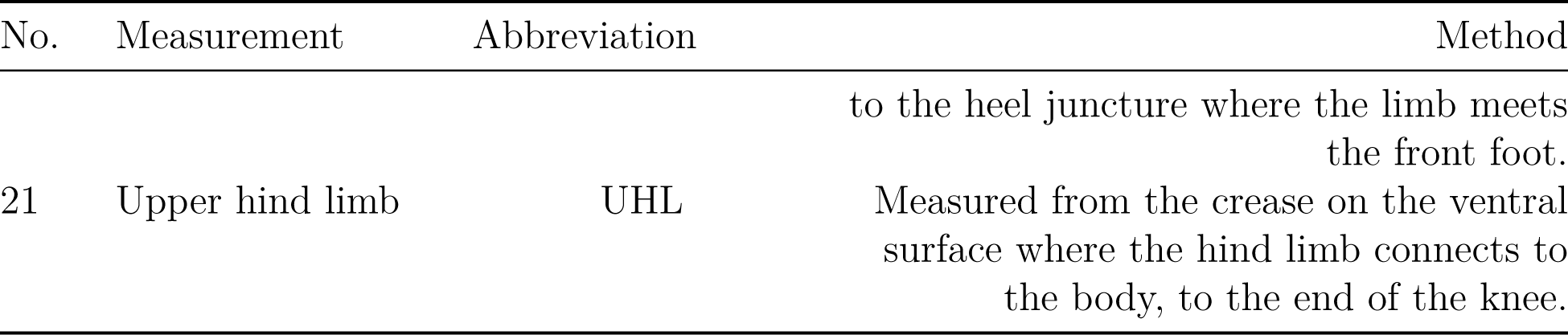
Initial morphological measurements used for data investigation:

**Table S4.** Final morphological traits used for phenotypic analyses:

| No. | Measurement | Shorthand | Method |
| --- | --- | --- | --- |
| 3 | Interlimb length | Interlimb | see above |
| 4 | Body width | Body_Width | see above |
| 5 | Pelvic width | Pelvic_Width | see above |
| 6 | Pelvic height | Pelvic_Height | see above |
| 7 | Head width | Head_Width | see above |
| 9 | Snout length | Snout_Eye | see above |
| 10 | Eye diameter | Eye_Diameter | see above |
| 11 | Head depth | Head_Depth | see above |
| 12 | Tail width | Tail_Width | see above |
| 13 | Tail length | Tail_Length | see above |
| 17 | Upper front limb | Upper_Arm | see above |
| 16 | Lower front limb | Lower_Arm | see above |
| 15 | Front foot | Hand | see above |
| 21 | Upper hind limb | Upper_Leg | see above |
| 20 | Lower hind limb | Lower_Leg | see above |
| 19 | Hind foot | Foot | see above |
| 22 | Neck length | Neck | Snout_Axilla - Head_Length |
| 23 | Posterior skull | Pos_Skull | Head_Length - (Snout_Eye + Eye_Diameter) |
| 24 | Pelvic gap | Pelvic_Gap | Snout_Vent - (Interlimb + Snout_Axilla) |

**Figure S6:**
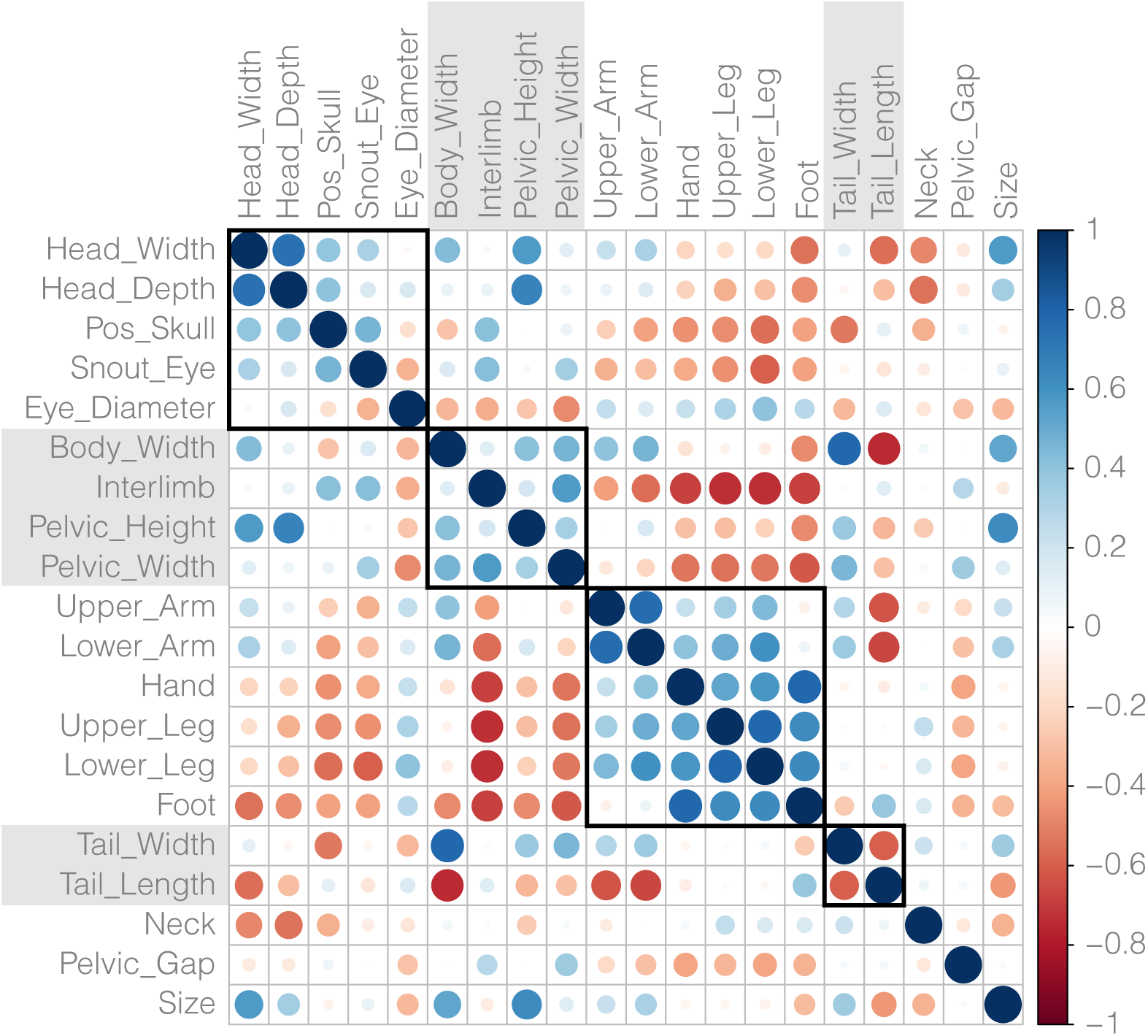
Correlation plot of all morphological traits after removing size via log-shape ratios. Some traits retain strong correlations despite removing the effect of size. Traits are organized according to the morphological module they belong to. Plot generated using *corrplot* [Wei & Simko 2021] using the Pearson (parametric) correlation.

### Phylogenetic Analyses and Timetree Estimation

Specifics of alignment are covered above.

**Figure S7:**
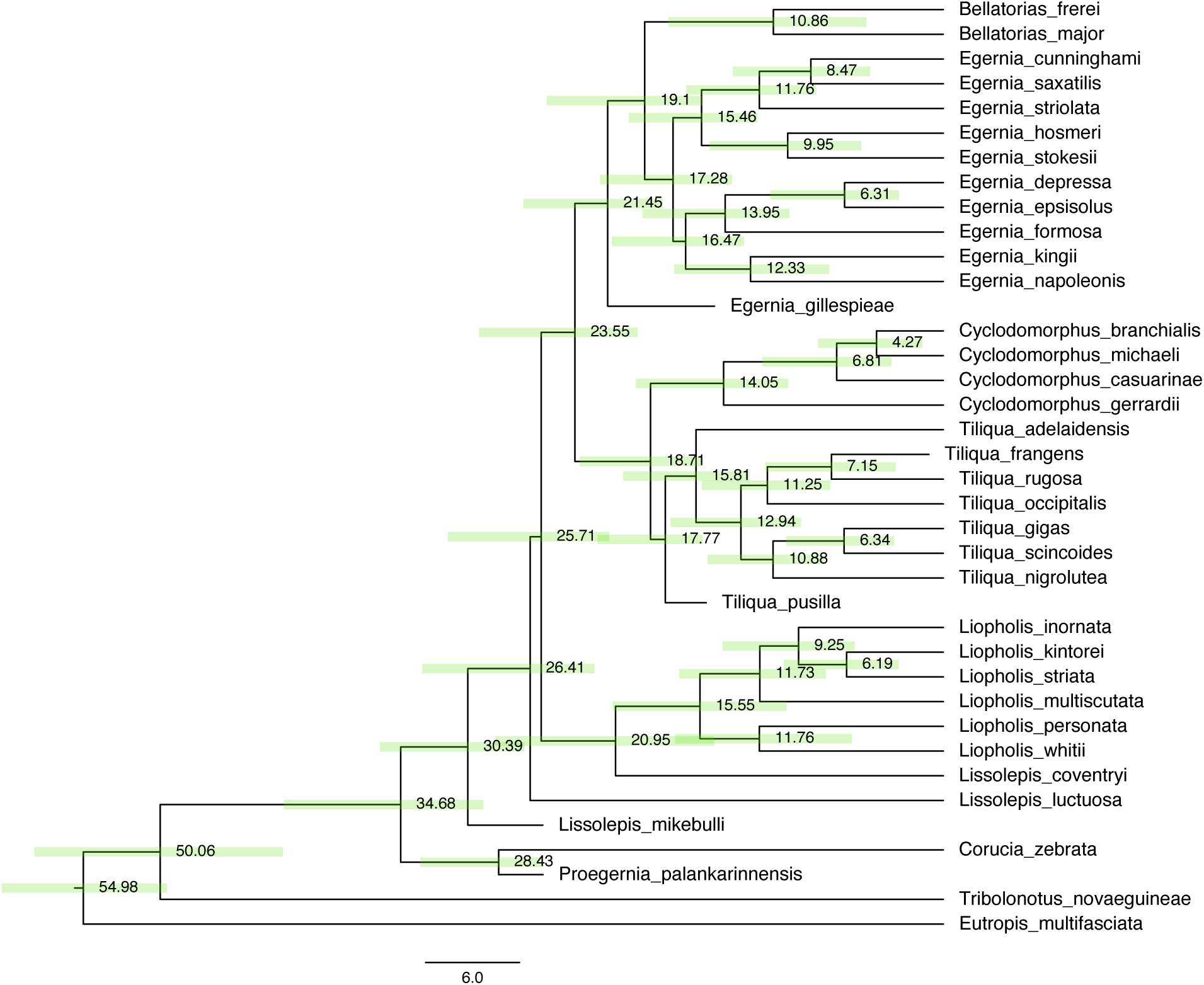
Combined evidence analysis of morphological and molecular data in BEAST 1.8.4, incorporating morphological data from Thorn et al. (2019), Thorn et al. (2021), Thorn et al. (2023). This analysis recovers a topology largely consistent with our molecular data, but provides insight into the phylogenetic placement of fossil taxa. Notably, we do not recover a monophyletic *Proegernia*, with *P.mikebulli* (in tree as *Lissolepis mikebulli*) placed as a stem Australian tiliquine, and we identify *Tiliqua pusilla* as a likely stem *Tiliqua*, and *Egernia gillespieae* as a stem lineage leading to *Bellatorias* and *Egernia*. As a result of the inclusion of morphological data, all *Cyclodomorphus* are drawn into a single clade, likely as a result of shared plesiomorphic characters. Estimated divergence dates are likely inflated by severe rate heterogeneity in morphological characters (see Fig.S8).

**Figure S8:**
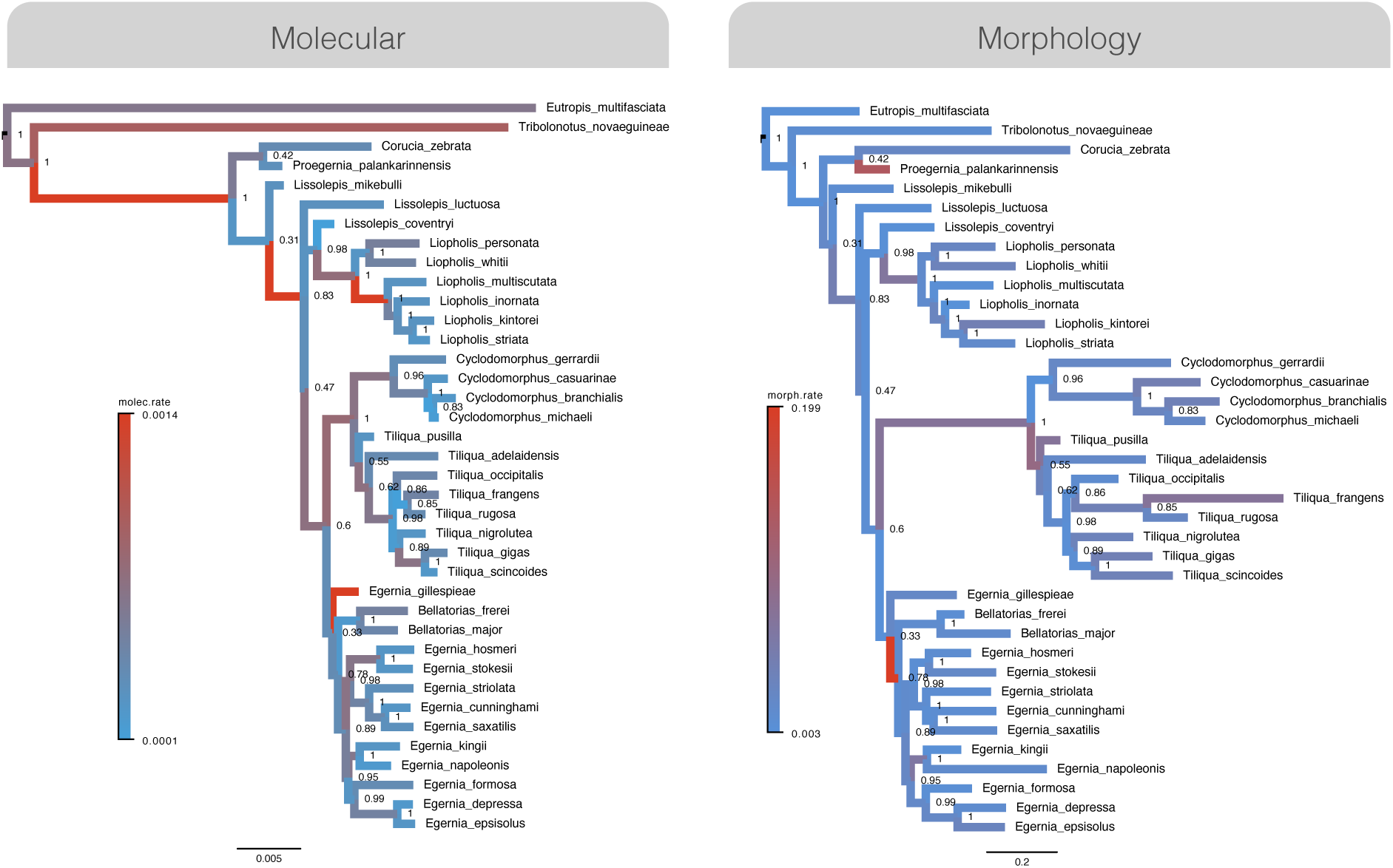
Maximum clade credibility trees based on molecular (left) and morphological (right) estimated in a combined-evidence BEAST analysis. The morphological tree shows strong rate heterogeneity in the lineage leading to *Cyclodomorphus/Tiliqua*.

**Figure S9:**
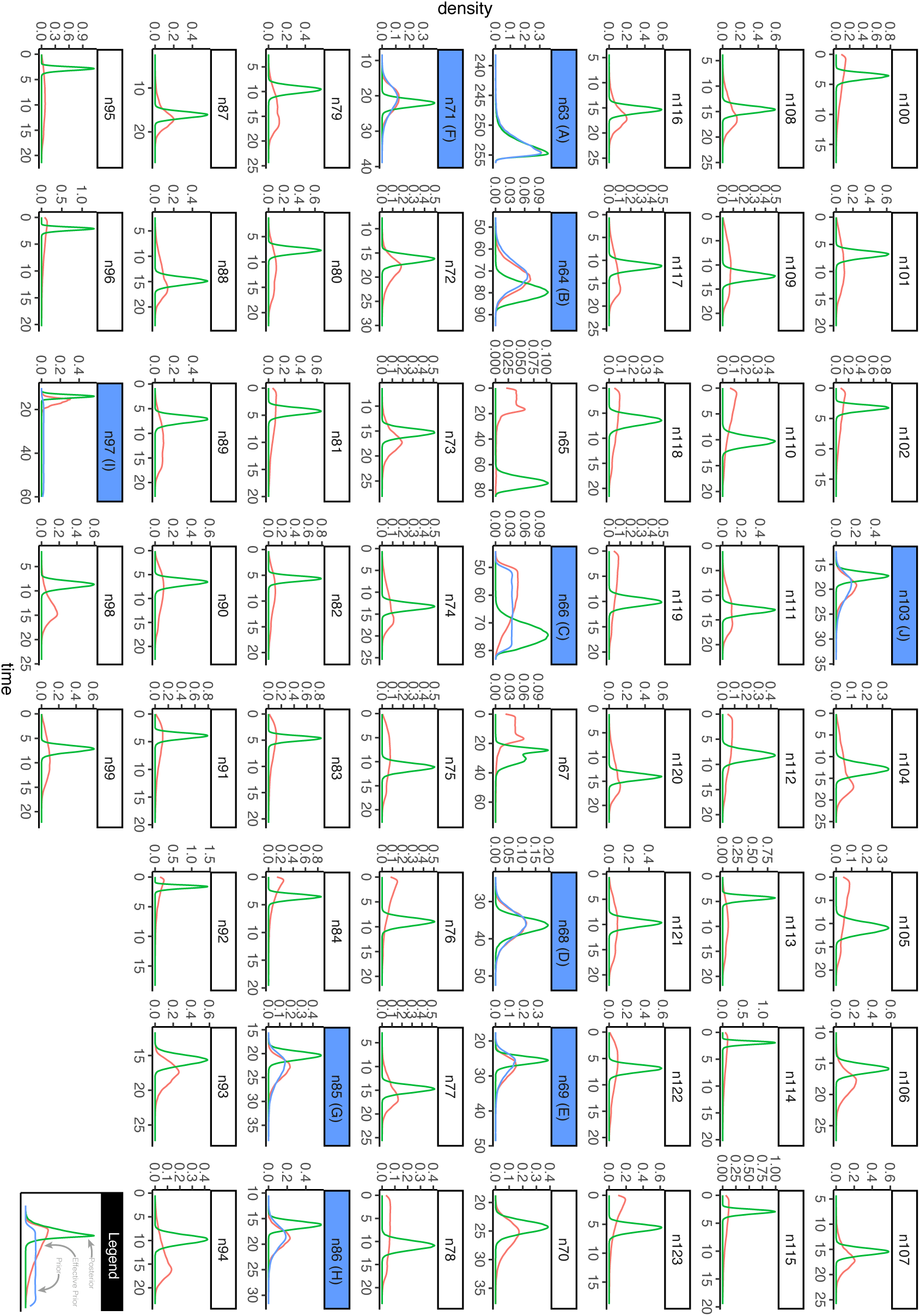
Plots of priors (blue), effective priors (pink), and posteriors (green) from MCMCtree analyses show reasonable behavior. Node numbers correspond to the phylogeny in *Trees/Tiliquini_MCMCTree.tre* and plotted with *ape*, where *n63* is the root node. Nodes with a blue header indicate those which included a prior.

**Figure S10:**
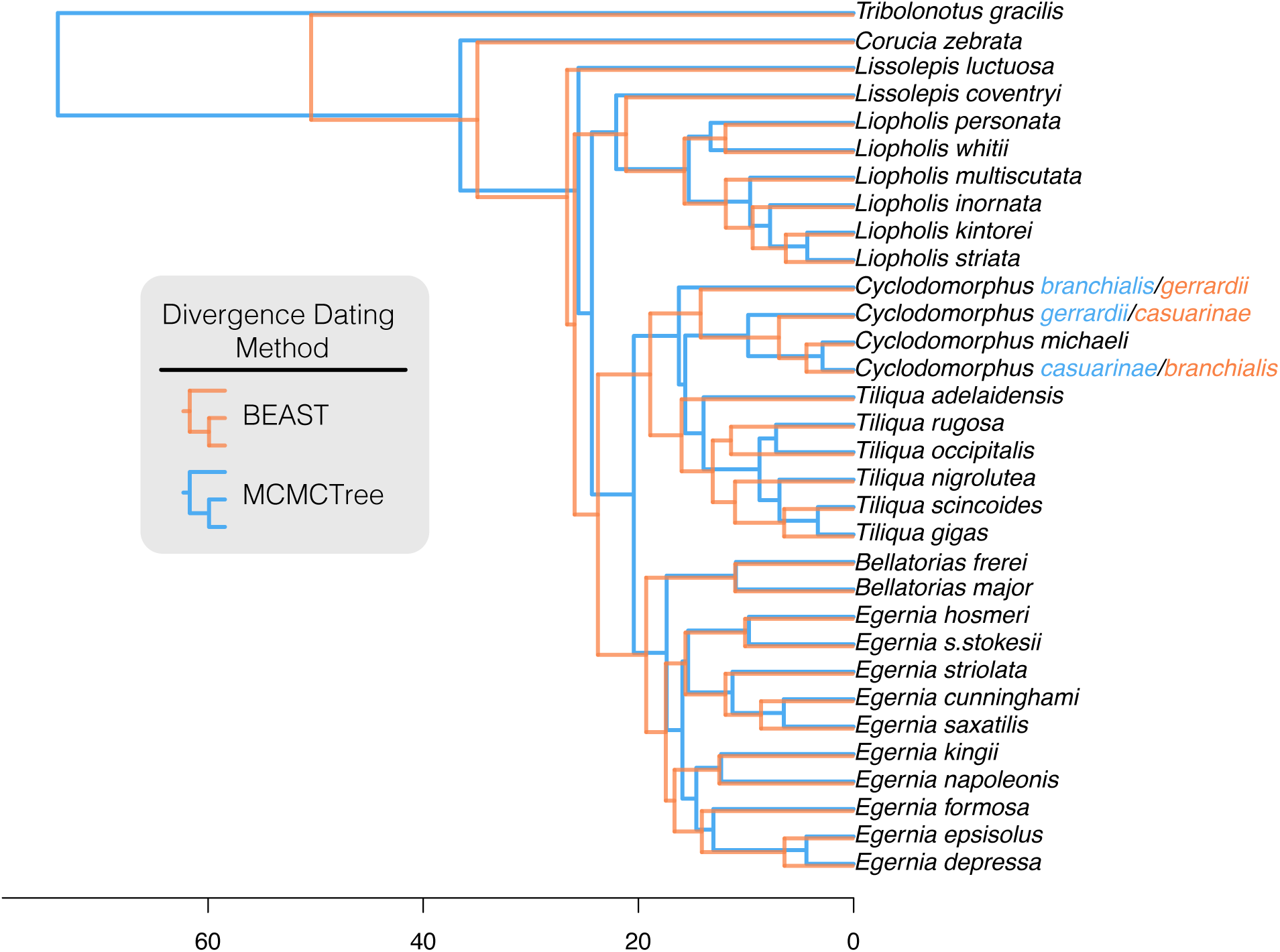
Comparison of divergence estimates under two different divergence dating methods. Trees were downsampled to maximize taxon overlap for visualization purposes. Blue phylogeny shows results of node-calibrated molecular analysis in MCMCTree and orange phylogeny shows results from combined evidence tip dating in BEAST. Node priors in MCMCTree analysis were informed by 95% confidence intervals estimated from the BEAST analysis, as shown in the calibration strategy in Table S2.

**Table S5:**
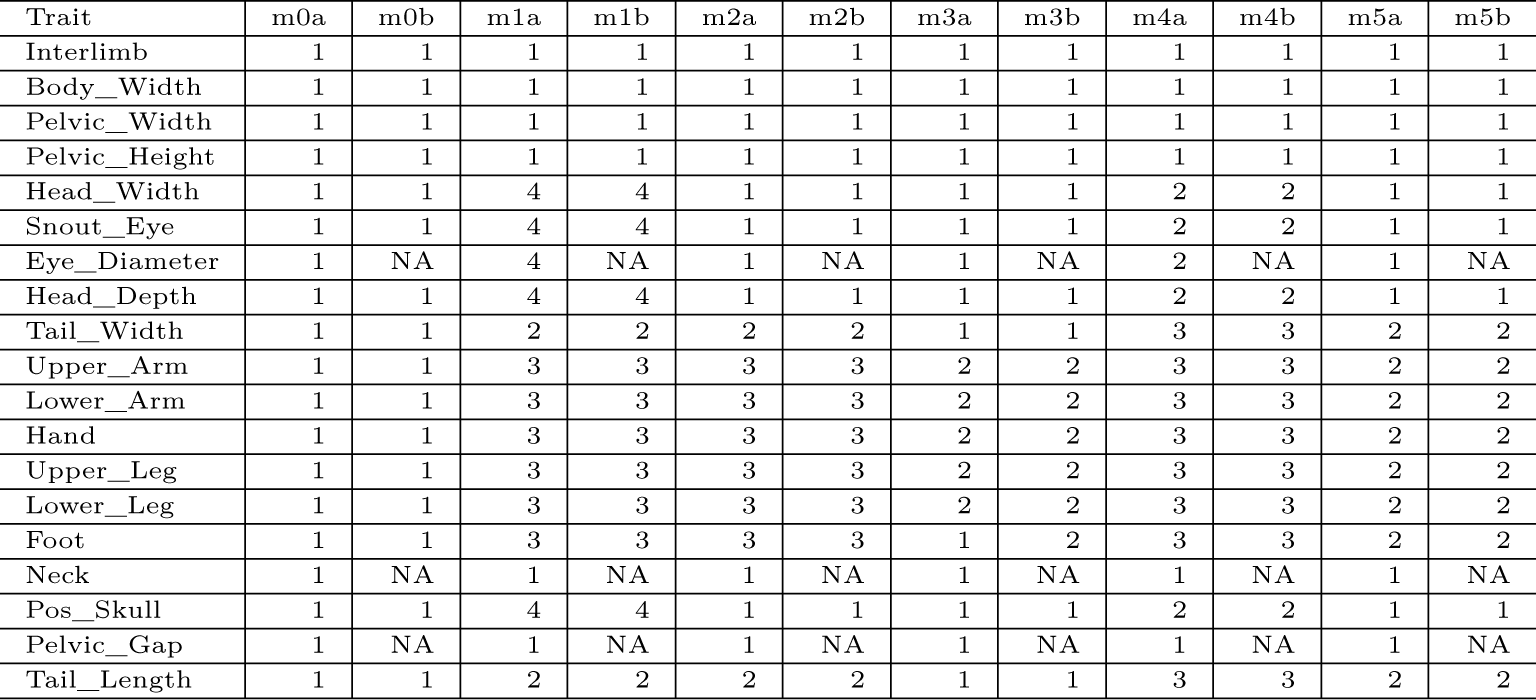
A set of morphological models for testing modularity with *EMMLi*. Each column represents a model and each row represents the occupancy of a given trait in that model. Module numbering scheme follows that required for *EMMLi*.

### Modelling Trait Evolution and Disparity

The below methodology is accompanied by a series of scripts found in /Scripts which will run on the included /Data files, These steps can be followed chronologically and are scripts are indicated as such `<Number>_<Process>.R e.g. 00_Data_Preparation.R.`

**Figure S11:**
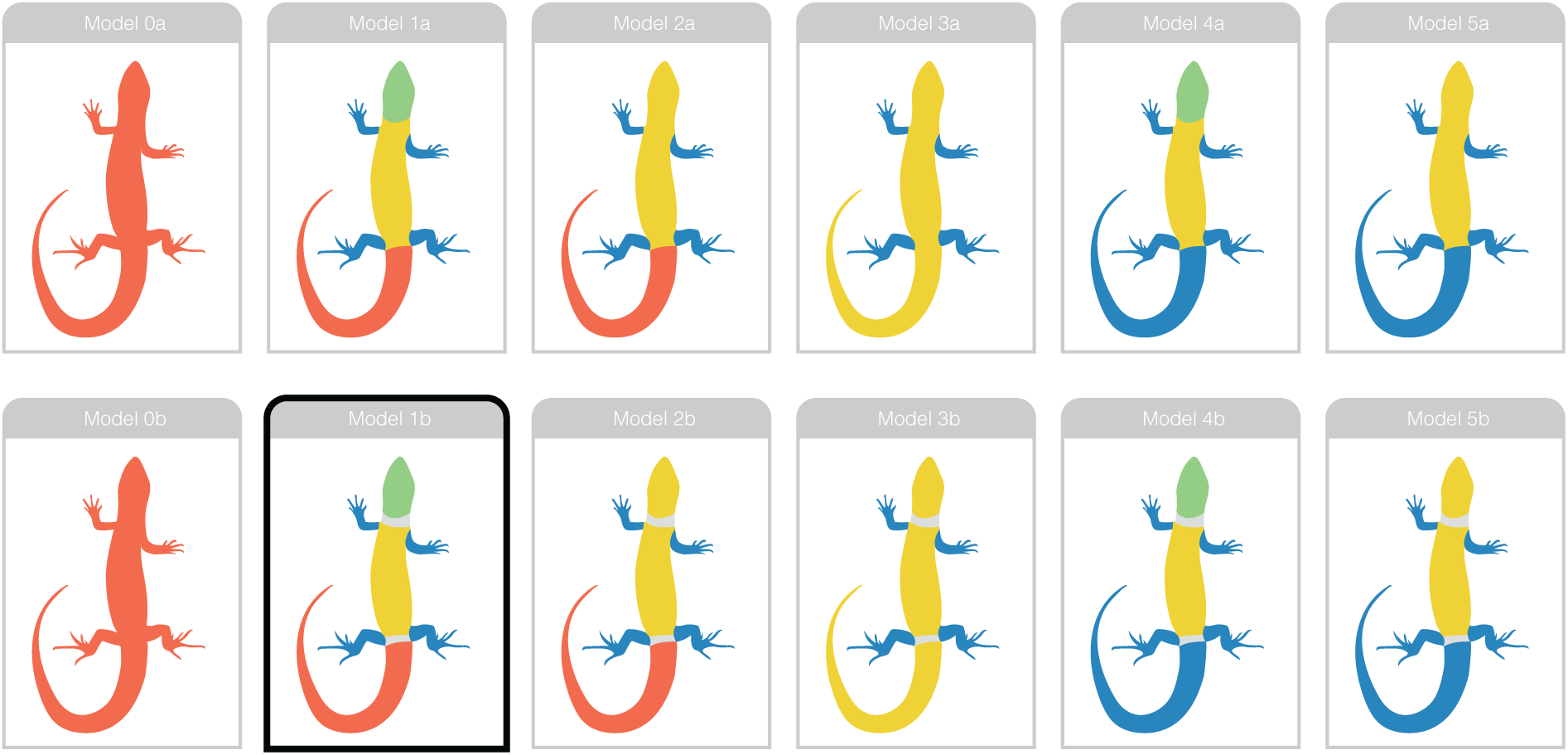
Diagram indicating the models for testing morphological modularity with *EMMLi*. Top and bottom rows indicate slight variations on the same basic model, with the bottom model (‘b’) treating three traits (neck length, pelvic gap length, eye diameter) that did not fit into modules neatly as ‘unintegrated’. The preferred model *Model 1b* is highlighted with a black outline. Trait occupancy in specific models are indicated in Table S5.

**Figure S12:**
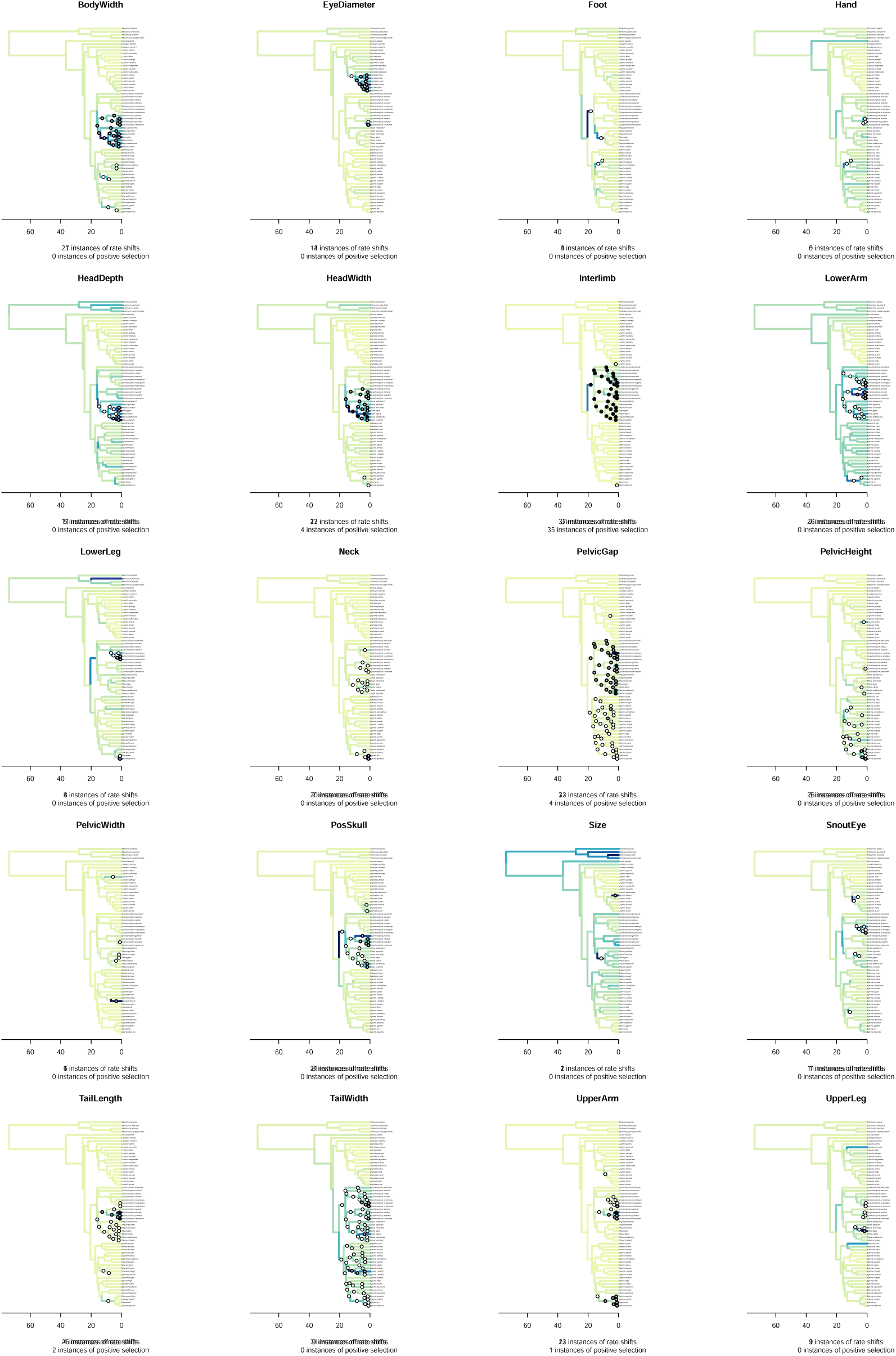
Pulses in rates of phenotypic evolution are common across tiliquine morphological traits. Species tree branches are colored according to mean estimated sigma value. Circles indicate significant rate shifts: empty circles represent shifts which appeared in > 70% of the posterior samples and in which the mean estimated scalar > 2; black circles represent shifts which appeared in >95% of the posterior samples and in which the mean estimated scalar > 2 corresponding to instances of “positive phenotypic selection” per _5_B_0_aker et al. 2017.

**Figure S13:**
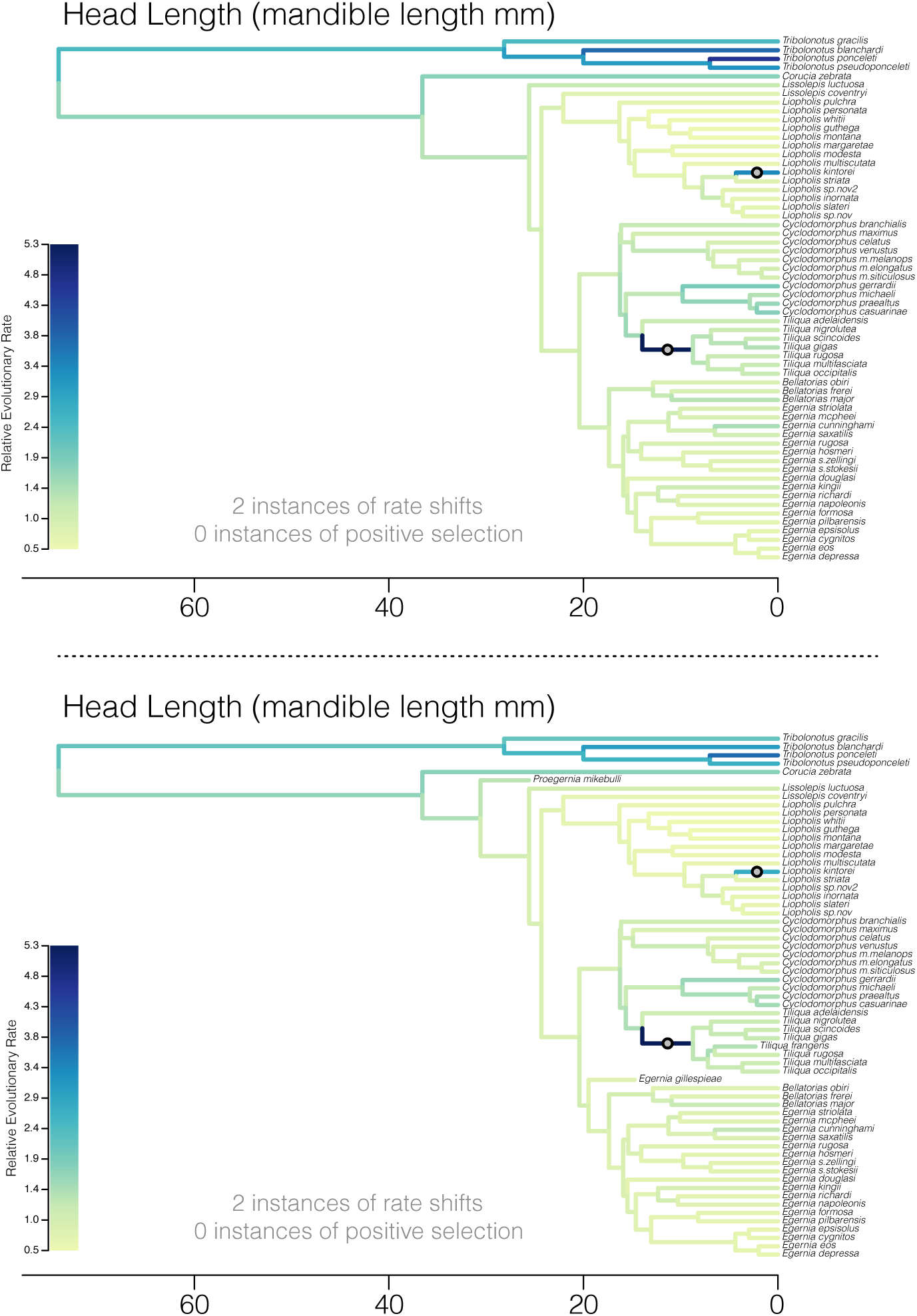
Incorporating extinct Tiliquini species does not change inferences of trait evolution. Trees show the estimated pattern of evolution of head length in Tiliquini skinks, with branches colored according to relative evolutionary rate and shifts indicated by circles on branches. (Top) Phylogenetic tree of extant Tiliquini and (bottom) phylogenetic tree of extinct and extant species return the same inferred shifts and qualitatively similar evolutionary rates. Additional fossil information may elucidate additional shifts in trait evolution, but are unlikely to erase inferred shifts. This is highlighted by the inclusion of *Tiliqua frangens* in the additional head length analysis. *T.frangens* has been estimated (Thorn et al. 2023) to have been more than twice the size of its close relative *T.rugosa* (SVL: 250 mm vs. 590 mm). In order to achieve this disparity in size, however, it is only necessary to increase the estimated evolutionary rate marginally. In comparison, a far higher evolutionary rate (5x the mean/background rate) is required to generate the comparatively huge heads of *Tiliqua* as shown on the dark blue branch in both trees.

**Figure S14:**
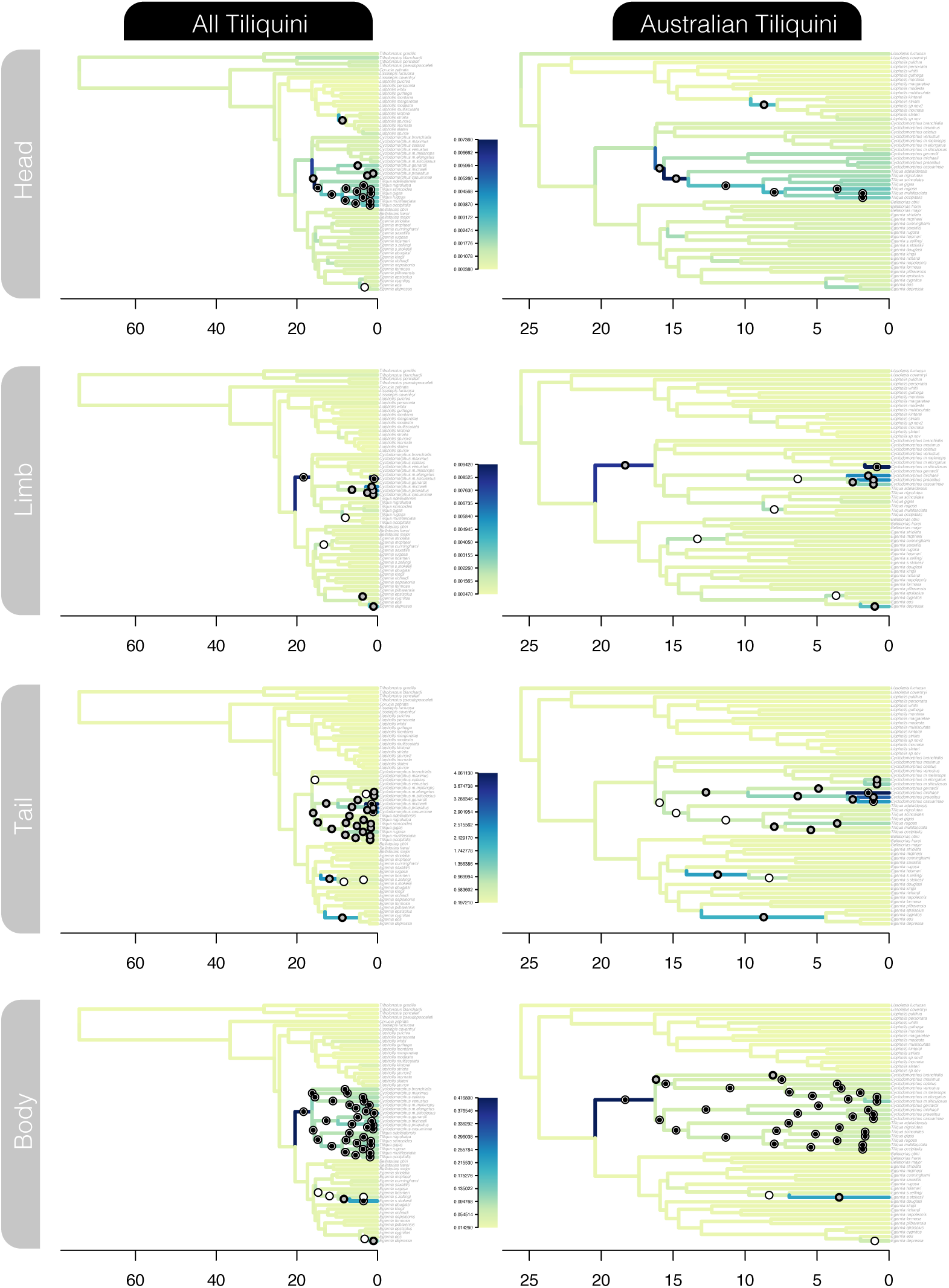
The inclusion of taxa on long branches (*Tribolonotus*, *Corucia*) does not bias the estimation of evolutionary rates across the rest of the tree. (left column) Analyses including non-Australian tiliquine taxa *Tribolonotus* and *Corucia*, and (right column) including ***only*** Australian tiliquine taxa. Trees show the output of the VR model applied to each labelled module (rows), with branches colored according to common color scales shown in the middle, plotted using *plot.VarRates.tree* in *Scripts/plotting_BayesTraits.R*. Estimated rates and inferred shifts are highly consistent across sampling strategies.

**Figure S15:**
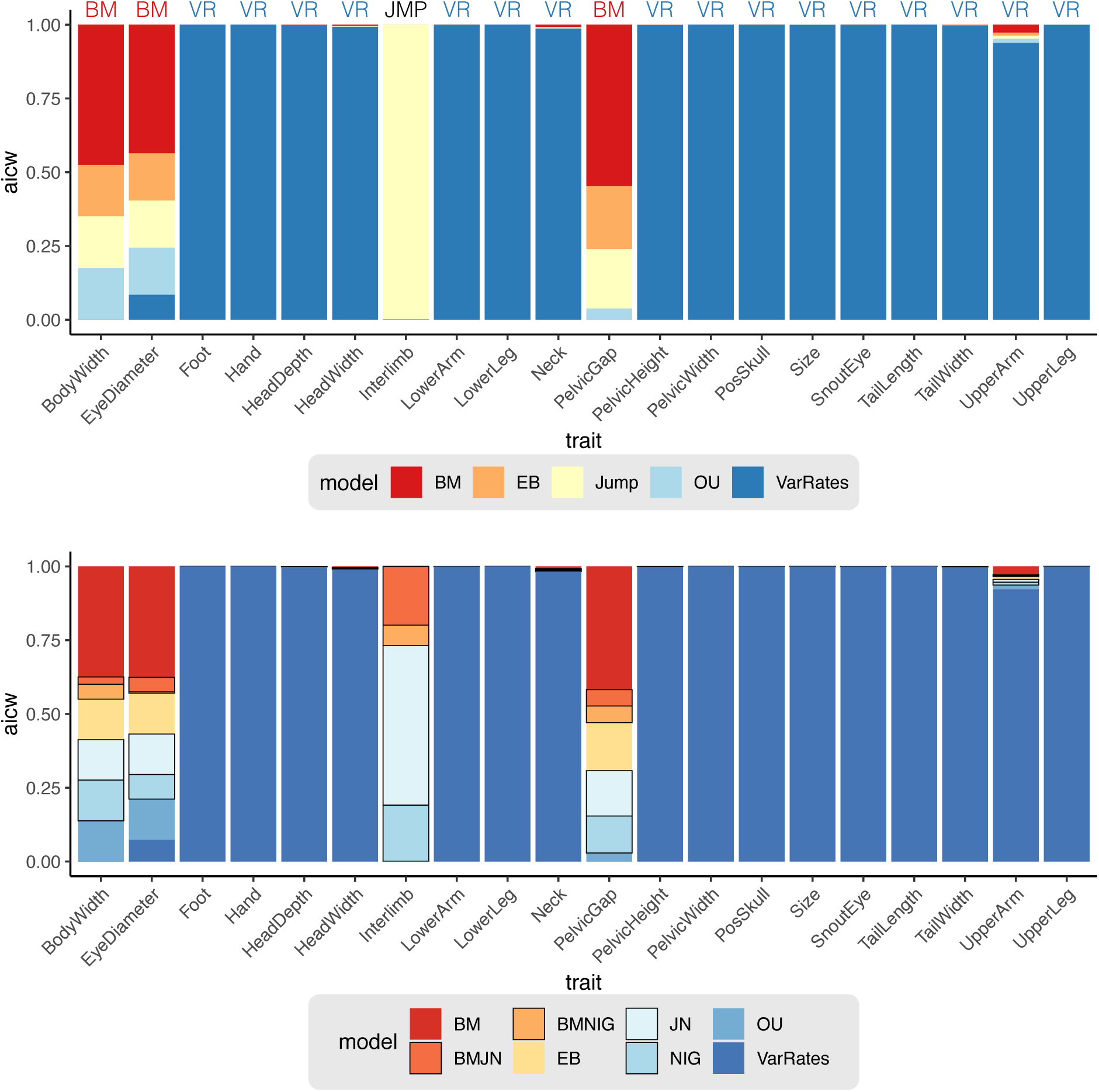
Comparative model fits for each of 19 morphological traits and the summary trait ‘Size’. Upper figure summarizes AICw by model type (BM, EB, ‘Jump’, OU, or VarRates), with the preferred model indicated above the column. Model preference required AICw of the best fitting model to be at least twice that of the next best fitting model. Bottom figure similarly shows model preference but with ‘Jump’ models expanded into their four alternative models (BMJN, BMNIG, JN, NIG) indicated by black outlines.

**Figure S16:**
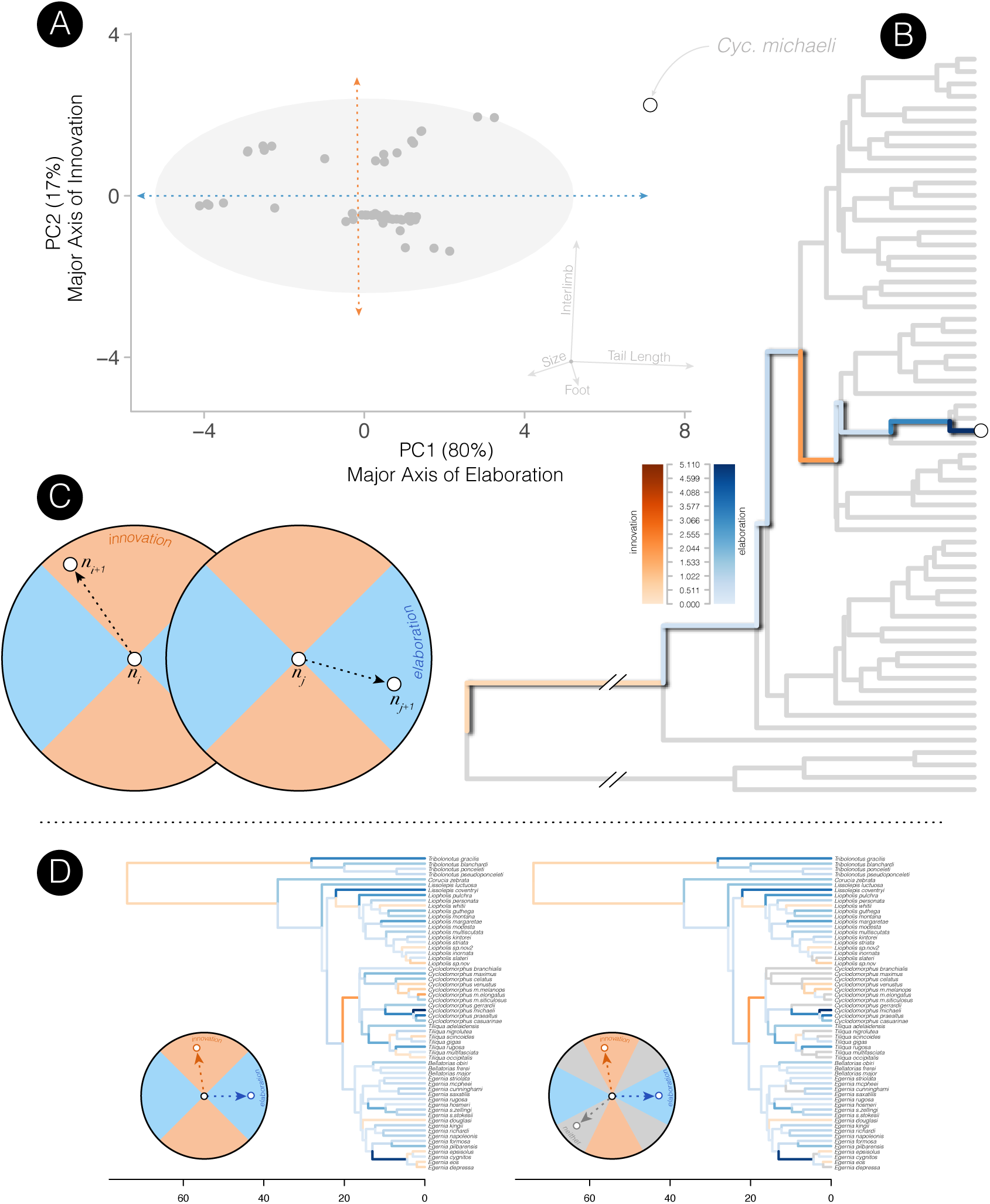
Diagram of method design and discretization of elaboration and innovation across Tiliquini skinks. (A) PCA biplot of first two axes indicating the path from root node to a specific tip, this path corresponds to the colored branches in the tree in (B). In (C) we show thresholds for designating elaborative and innovative trends, with parent node indicated at the center of the polar plot (*n_i_*, *n_j_*) and child nodes (*n_i+1_*, *n_j+1_*) plotted in either elaborative or innovative positions. Changing the threshold to a more conservative coding (D) does not change the overall inferences.

### Concatenation and the Anomaly Zone

To investigate topological differences between coalescent and concatented species trees we assessed the likelihood that discordant branches fall into the anomaly zone, where concatenation may be positively misled. To identify anomalous branches we used *Anomaly Finder* ^32^, using the script from Chafin et al. 2021.

**Figure S17:**
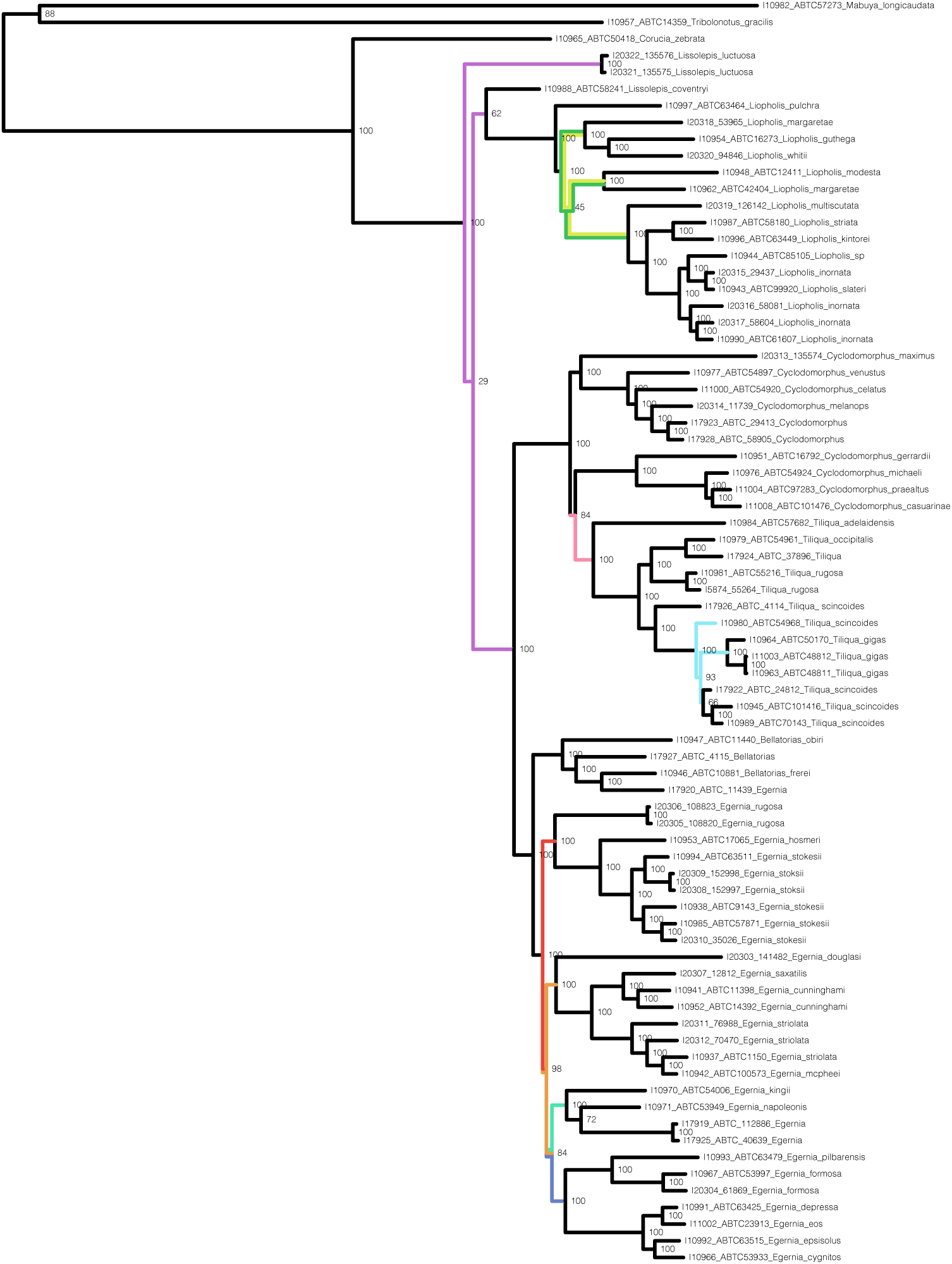
The Tiliquini phylogeny estimated from a concatenated and locus-partitioned alignment is likely misled by branches which fall into the anomaly zone. Colored branches correspond to the anomaly zone cladogram below and indicate edges which differ from the coalescent species tree. We ran *AnomalyFinder* following the scripts of Chafin et al. 2021.

**Figure S18:**
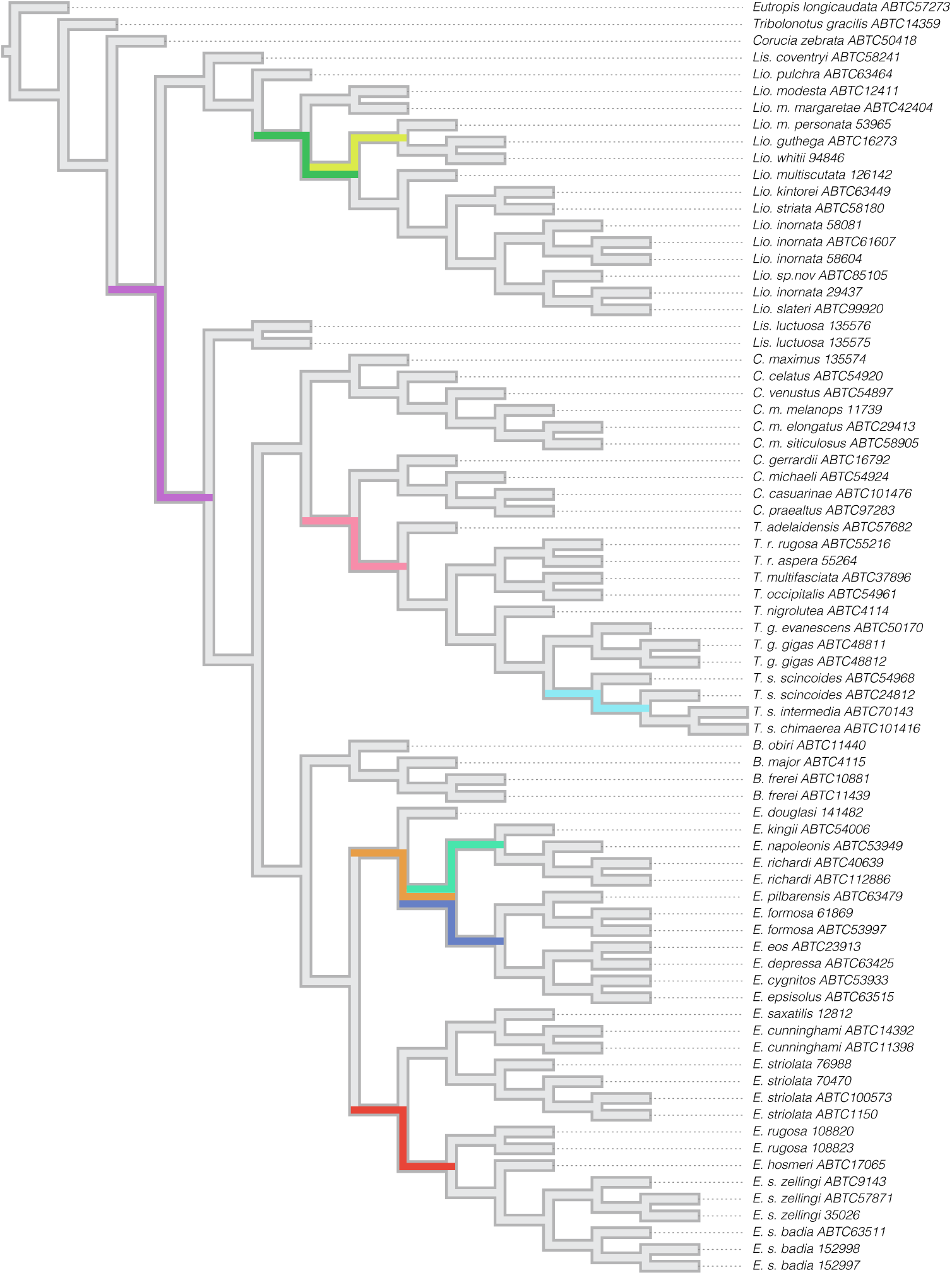
Colored branches indicate edges that fall within the anomaly zone and may provide misleading results under concatenation.

### Topology Tests

We used a series of topology tests in IQ-TREE to investigate two nodes which bear taxonomic implications. We used the concatenated alignment and simplified phylogenies as the input. In each case the preferred topology is presented first followed by the two alternative resolutions to the bipartition. Significant differences are denoted by (-), and show decisive preference for the coalescent species tree topology in comparison to alternative resolutions, reinforcing molecular phylogenetic evidence for the paraphyly of both *Cyclodomorphus* and *Lissolepis*.

Topology test for the paraphyly of *Cyclodomorphus* with regards to *Tiliqua*.

Tree topologies:

1. (Egernia,(C.maximus,(T.gigas,C.michaeli); : coalescent/concatenated species trees
2. (Egernia,(C.michaeli,(T.gigas,C.maximus); : alternative *Cyclodomorphus* paraphyly
3. (Egernia,(T.gigas,(C.maximus,C.michaeli); : *Cyclodomorphus* monophyly

| Tree | logL | deltaL | bp-RELL | p-KH | p-SH | p-WKH | p-WSH | c-ELW | p-AU |
| --- | --- | --- | --- | --- | --- | --- | --- | --- | --- |
| 1. | -1142007.338 | 0 | 0.97(+) | 0.98(+) | 1(+) | 0.98(+) | 0.993(+) | 0.971(+) | 0.981(+) |
| 2. | -1142067.499 | 60.161 | 0.03(-) | 0.02(-) | 0.026(-) | 0.02(-) | 0.039(-) | 0.0294(-) | 0.021(-) |
| 3. | -1142110.488 | 103.15 | 0(-) | 0(-) | 0(-) | 0(-) | 0(-) | 2.08e-10(-) | 1.79e-05(-) |

Topology tests for the paraphyly of *Lissolepis*.

Tree topologies:

1. (Outgroup,((Lis.luctuosa,E.striolata),(Lis.coventryi,Lio.inornata))); : coalescent species tree
2. (Outgroup,(Lis.luctuosa,(E.striolata,(Lis.coventryi,Lio.inornata)))); : concatenated species tree
3. (Outgroup,((Lis.coventryi,Lis.luctuosa),(Lio.inornata,E.striolata))); : *Lissolepis* monophyly

| Tree | logL | deltaL | bp-RELL | p-KH | p-SH | p-WKH | p-WSH | c-ELW | p-AU |
| --- | --- | --- | --- | --- | --- | --- | --- | --- | --- |
| 1. | -1326723.218 | 0 | 1(+) | 1(+) | 1(+) | 1(+) | 1(+) | 1(+) | 0.995(+) |
| 2. | -1326798.358 | 75.141 | 0(-) | 0(-) | 0.122(+) | 0(-) | 0(-) | 8.47e-07(-) | 0.00491(-) |
| 3. | -1327315.329 | 592.11 | 0(-) | 0(-) | 0(-) | 0(-) | 0(-) | 1.9e-174(-) | 3.58e-05(-) |

### Additional Supplementary Figures

**Figure S19:**
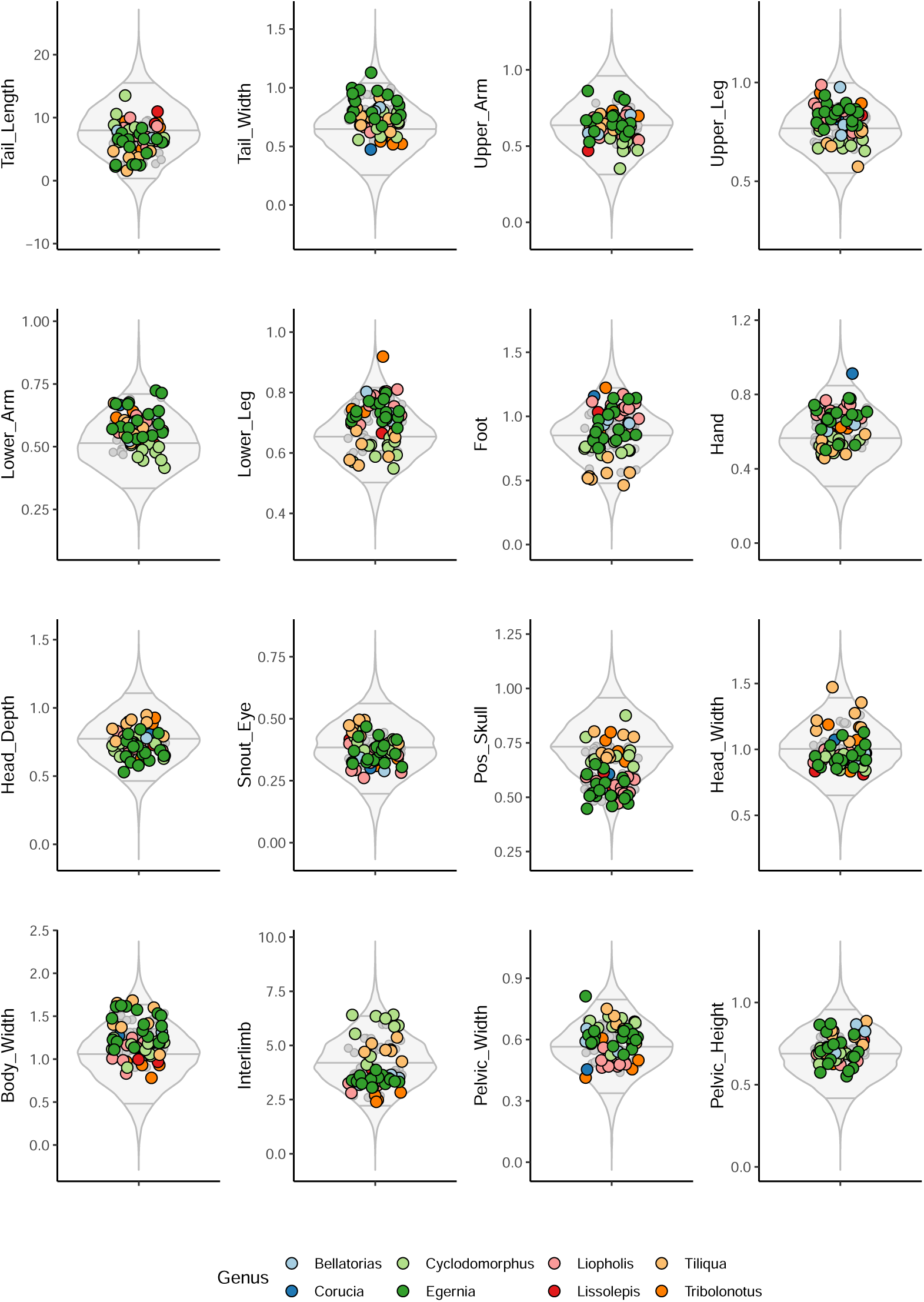
Empirical phenotypes generally conform to expected trait values of data simulated under uncorrelated evolution, showing limited ability of selection to exceed null expectations in Tiliquini skinks. Circles represent empirical trait values per species, colored by genus, with grey representing ancestral states. Transformed traits size corrected by log-shape ratios are plotted against 500 simulated datasets for each trait. Simulated traits are plotted as a grey violin plot summarizing the distribution of trait values, with595%, 50%, and 95% quantiles plotted as horizontal lines. Simulations were generated with MVMORPH using as input the theta–root and sigma values estimated by MVMORPH assuming a constant-rate Brownian Motion model.

**Figure S20:**
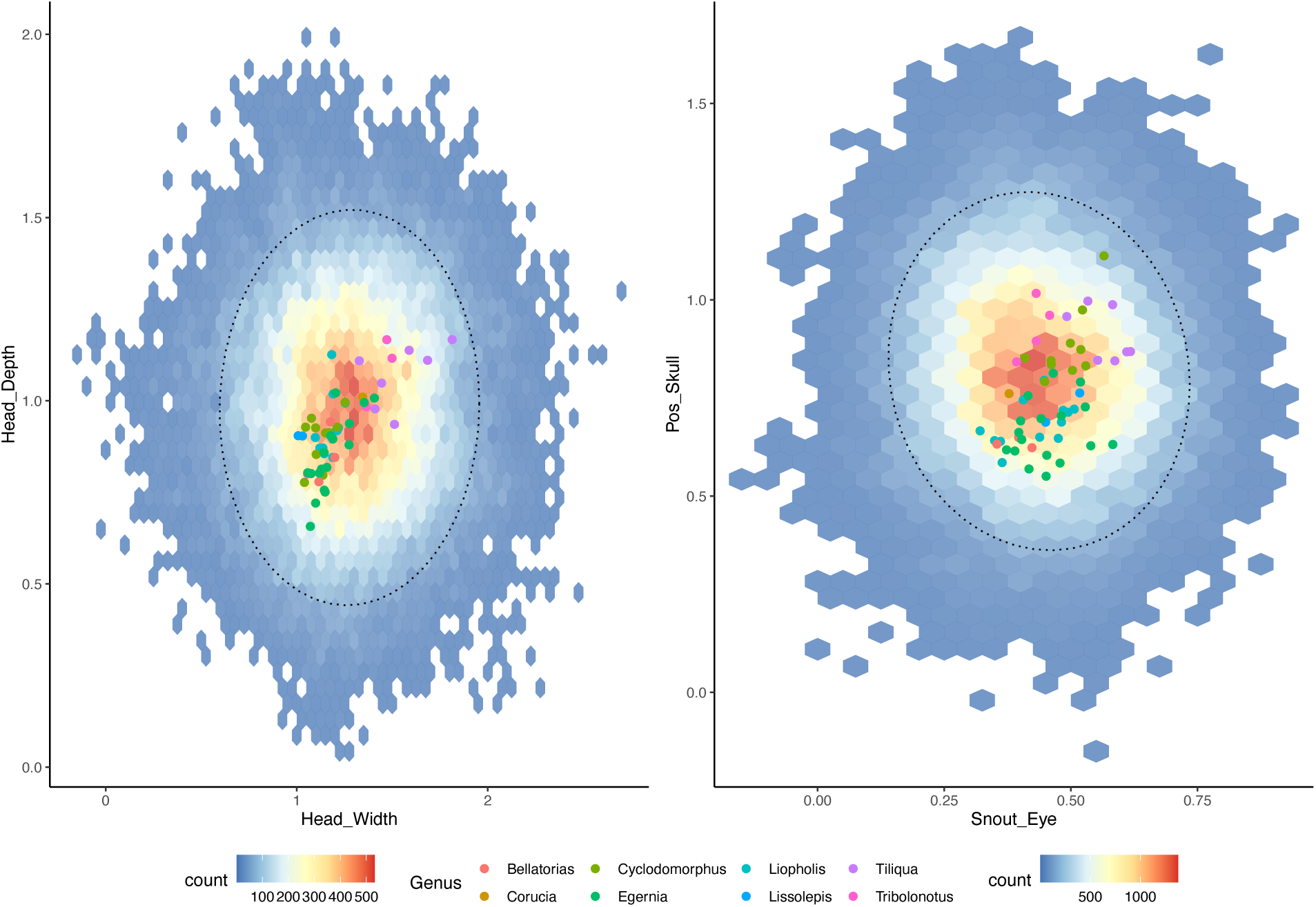
Empirical phenotypes generally conform to expected trait values of data simulated under uncorrelated evolution, showing limited ability of selection to exceed null expectations in Tiliquini skinks. Hexagonal heatplot in background shows the distribution of trait values from 500 datasets simulated under rates from empirical estimates. Warmer colors indicate greater density of simulated points. Circular points indicate empirical trait values and are colored according to the genus the belong to. The dotted ellipse contains 95% of the simulated data (95% confidence interval).

**Figure S21:**
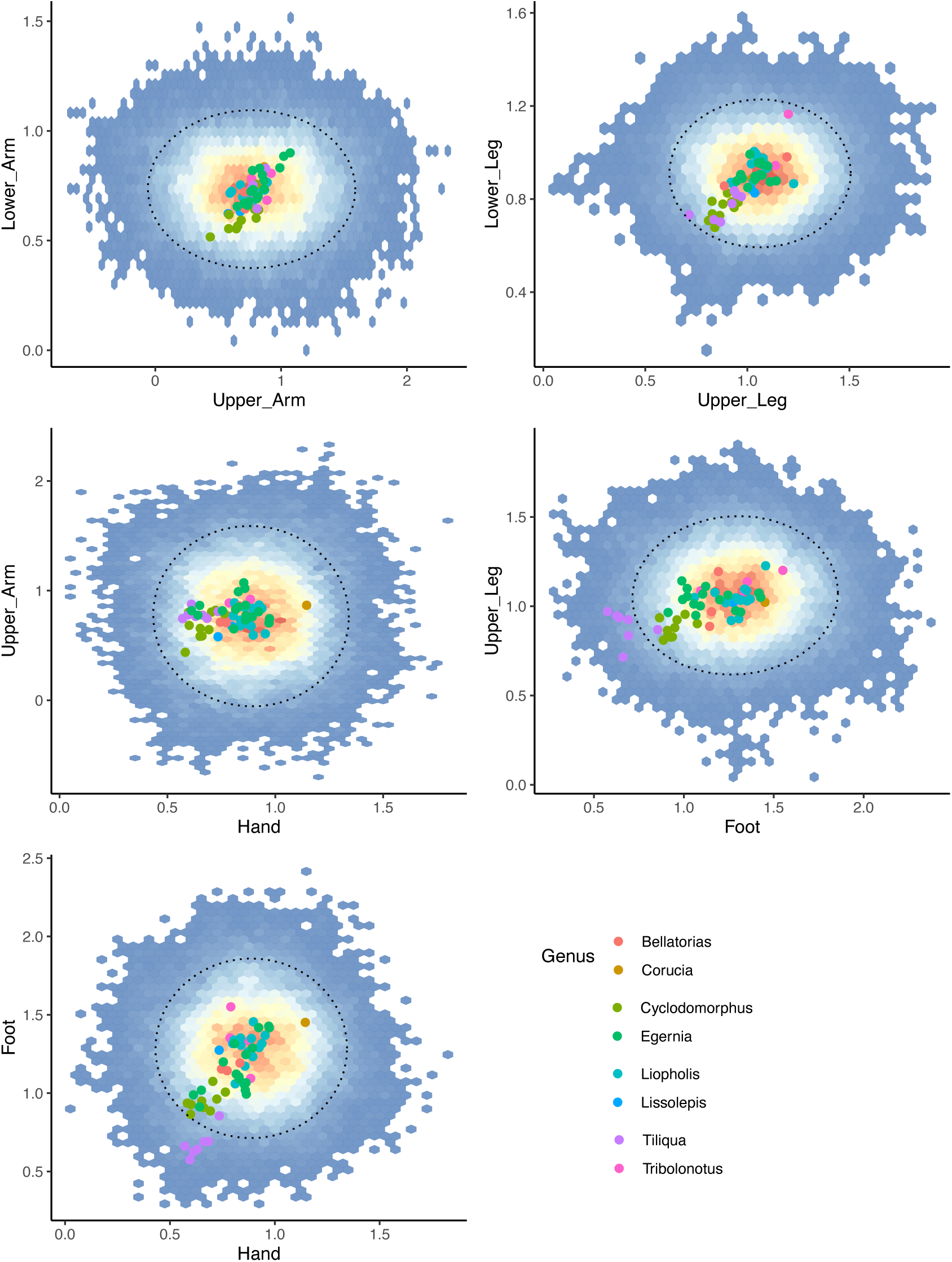
Empirical phenotypes generally conform to expected trait values of data simulated under uncorrelated evolution, showing limited ability of selection to exceed null expectations in Tiliquini skinks. Hexagonal heatplot in background shows the distribution of trait values from 500 datasets simulated under rates from empirical estimates. Warmer colors indicate greater density of simulated points. Circular points indicate empirical trait values and are colored according to the genus the belong to. The dotted ellipse contains 95% of the simulated data (95% confidence interval).

**Figure S22:**
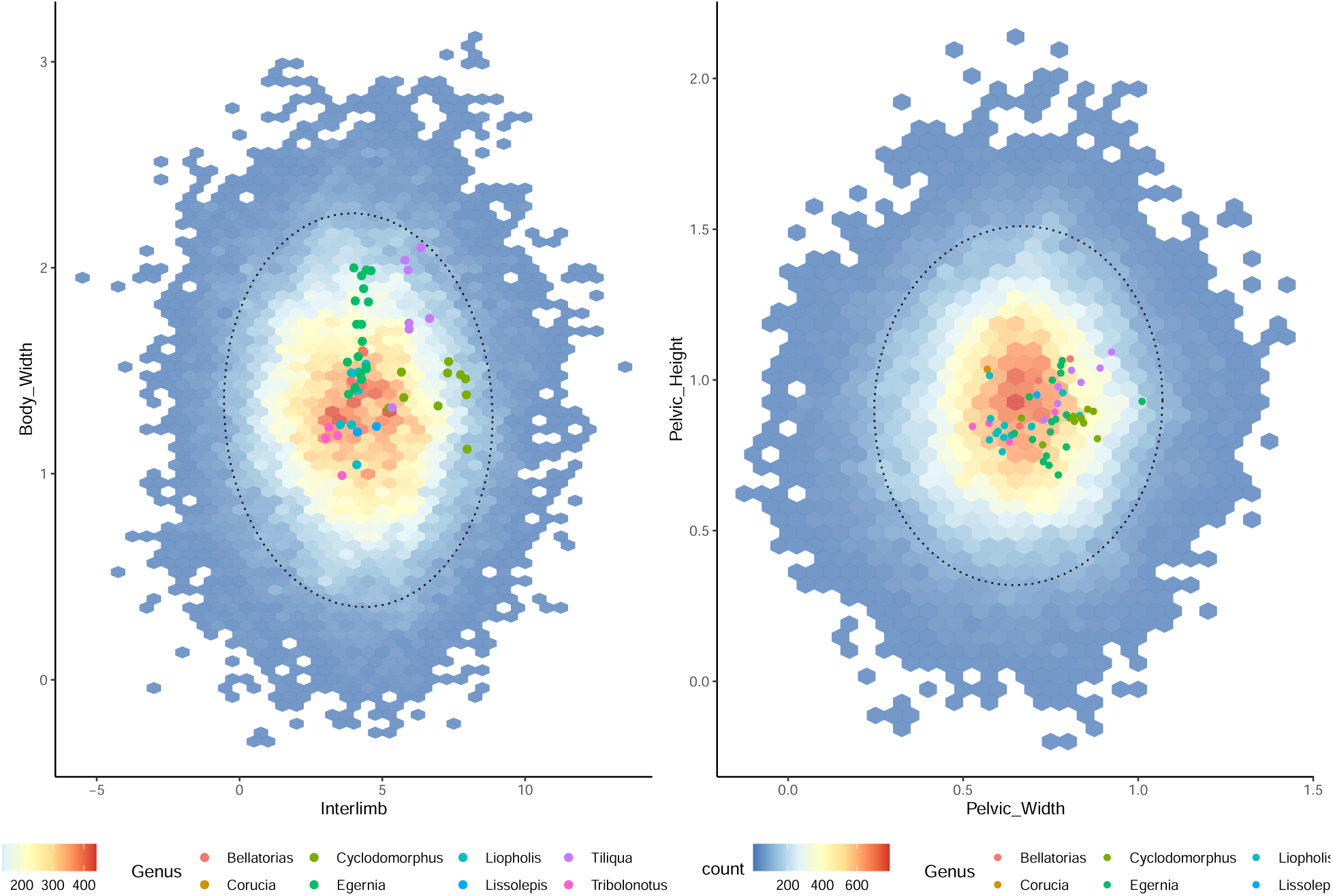
Empirical phenotypes generally conform to expected trait values of data simulated under uncorrelated evolution, showing limited ability of selection to exceed null expectations in Tiliquini skinks. Hexagonal heatplot in background shows the distribution of trait values from 500 datasets simulated under rates from empirical estimates. Warmer colors indicate greater density of simulated points. Circular points indicate empirical trait values and are colored according to the genus the belong to. The dotted ellipse contains 95% of the simulated data (95% confidence interval).

**Figure S23:**
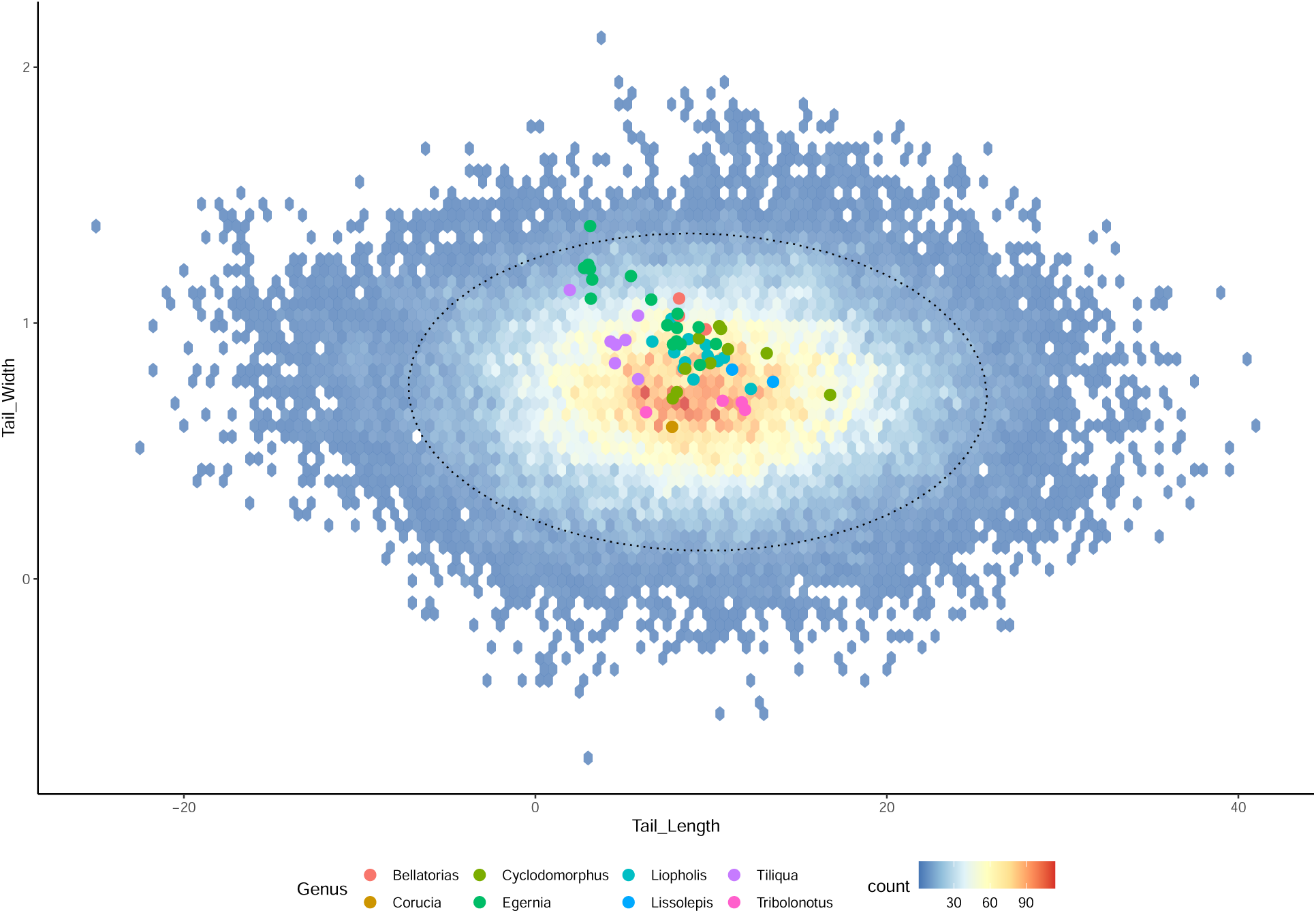
Empirical phenotypes generally conform to expected trait values of data simulated under uncorrelated evolution, showing limited ability of selection to exceed null expectations in Tiliquini skinks. Hexagonal heatplot in background shows the distribution of trait values from 500 datasets simulated under rates from empirical estimates. Warmer colors indicate greater density of simulated points. Circular points indicate empirical trait values and are colored according to the genus the belong to. The dotted ellipse contains 95% of the simulated data (95% confidence interval).

**Figure S24:**
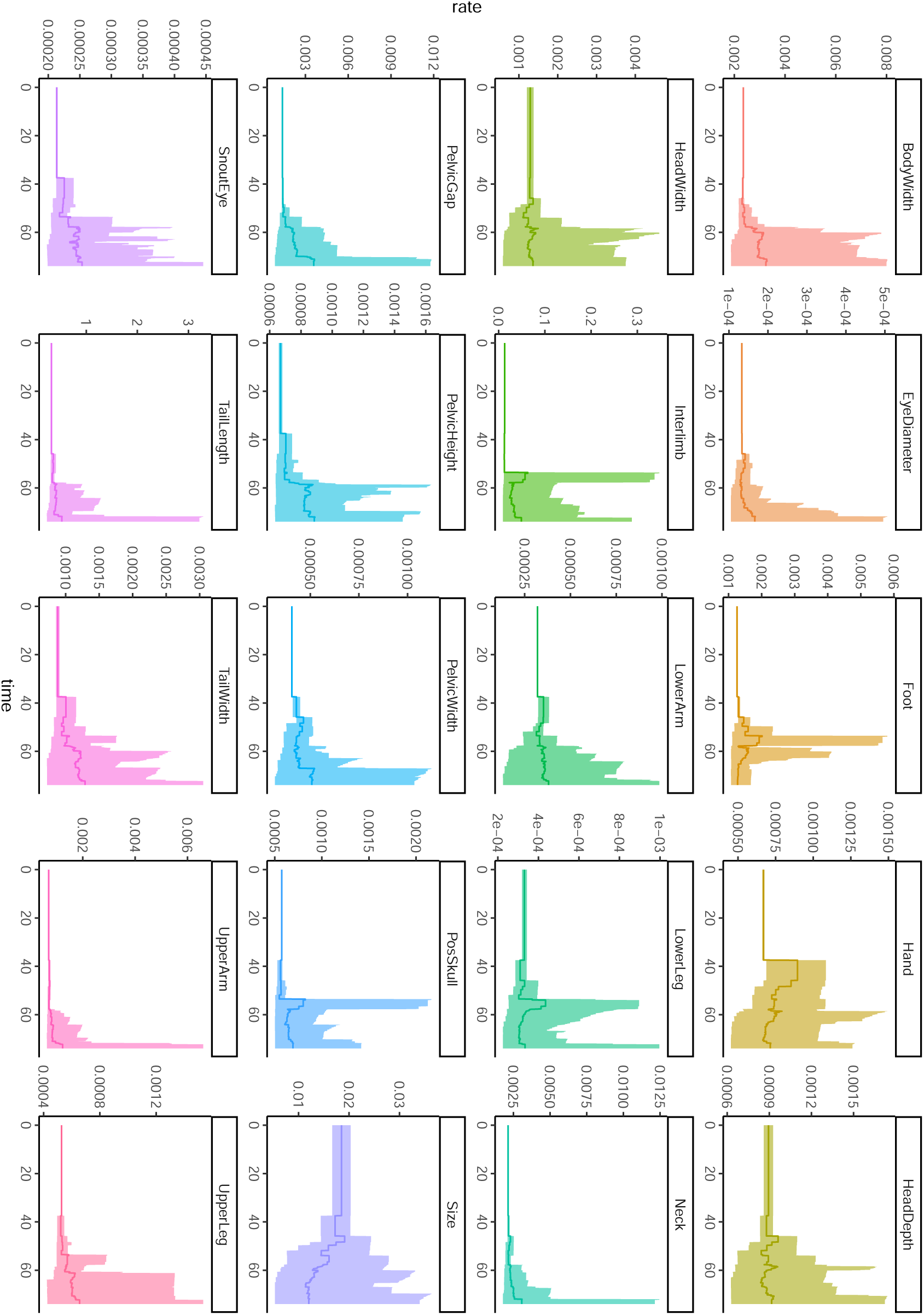
Mean evolutionary rates through time for all morphological traits with envelopes containing 95% quantiles.

**Figure S25:**
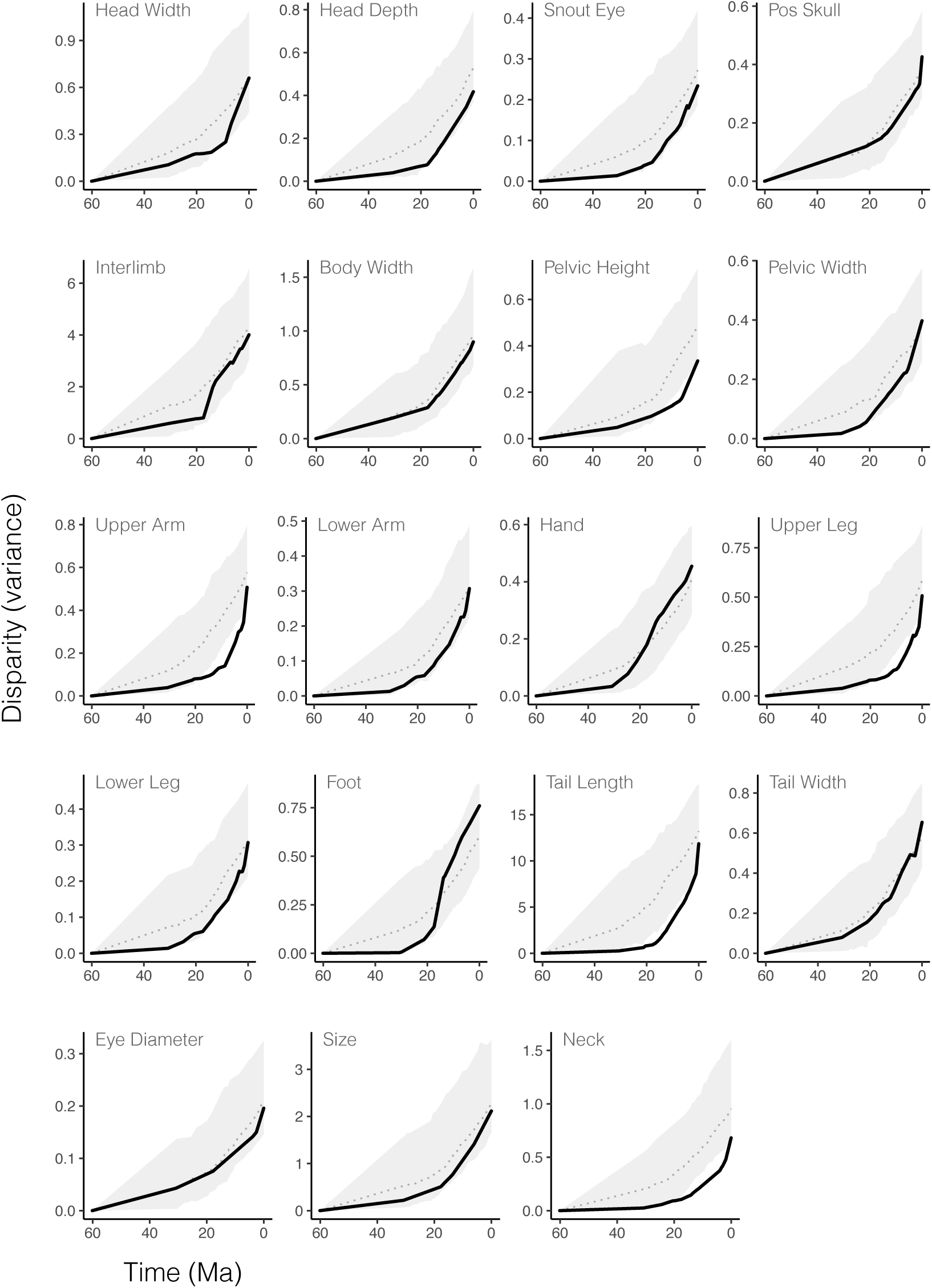
Disparity (variance) through time for morphological traits is often lower than expected under BM for long periods of time early in the evolution of this group. Empirical trends are shown in black with 95% quantiles of disparity for simulated datasets in grey.

**Figure S26:**
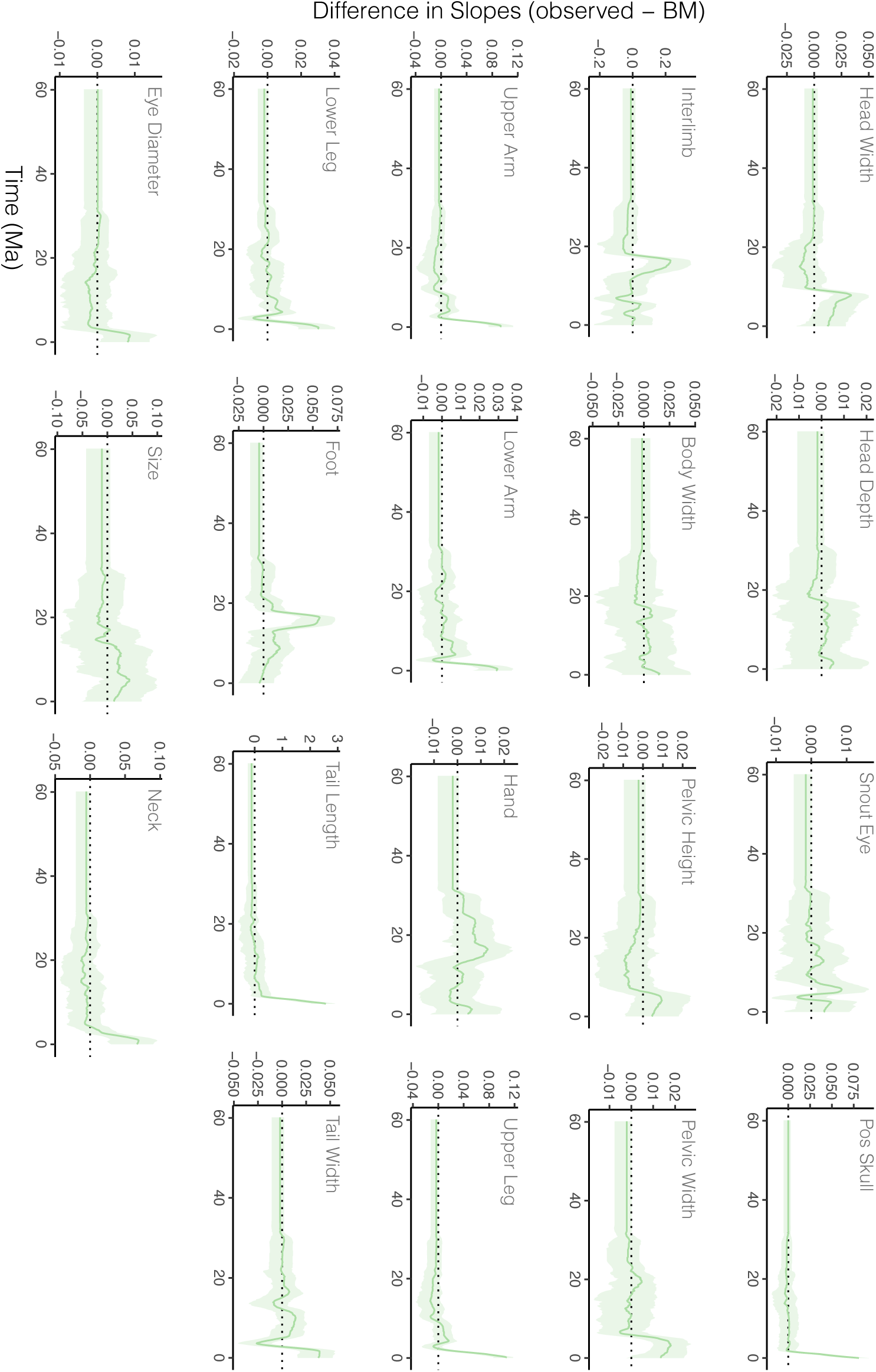
Individual traits show varied temporal patterns of morphological expansion and niche packing. Solid green line shows the mean trend and green envelope shows the 95% quantiles.

**Figure S27:**
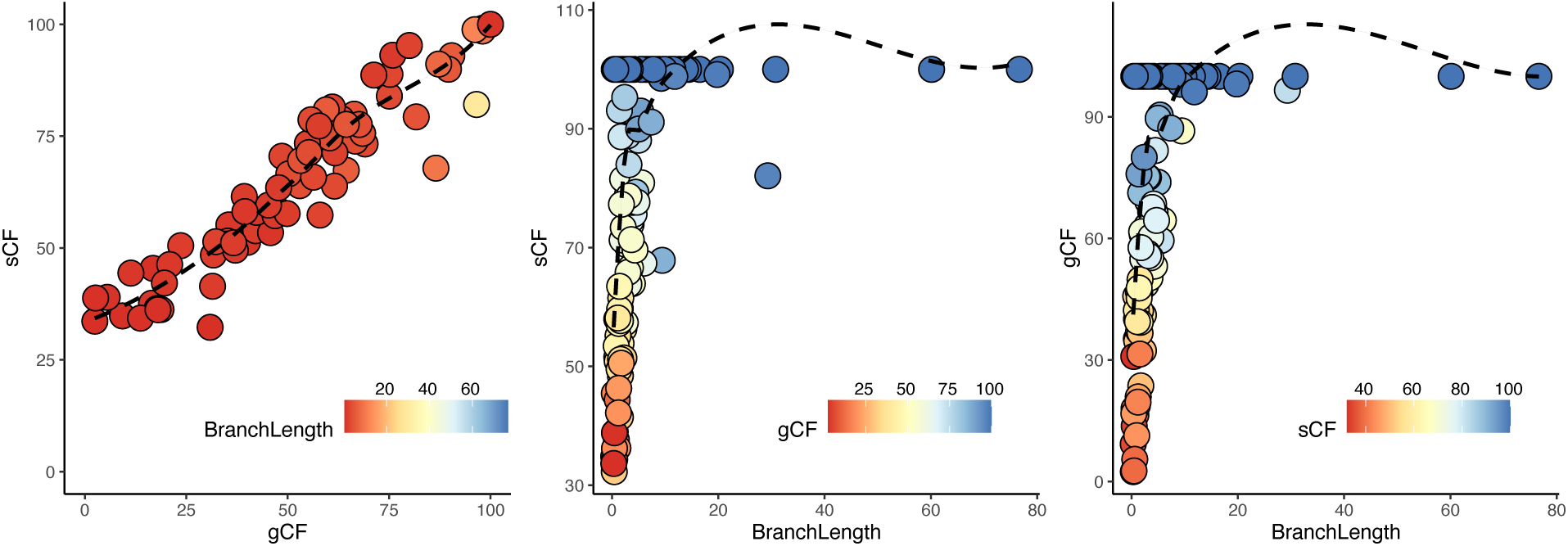
Concordance factors show a positive relationship with increasing branch length. Both site– and gene concordance factors increase support with increasing branch lengths (time), but show saturation (sCF=100, gCF=100) on branches >=10 million years long.

**Figure S28:**
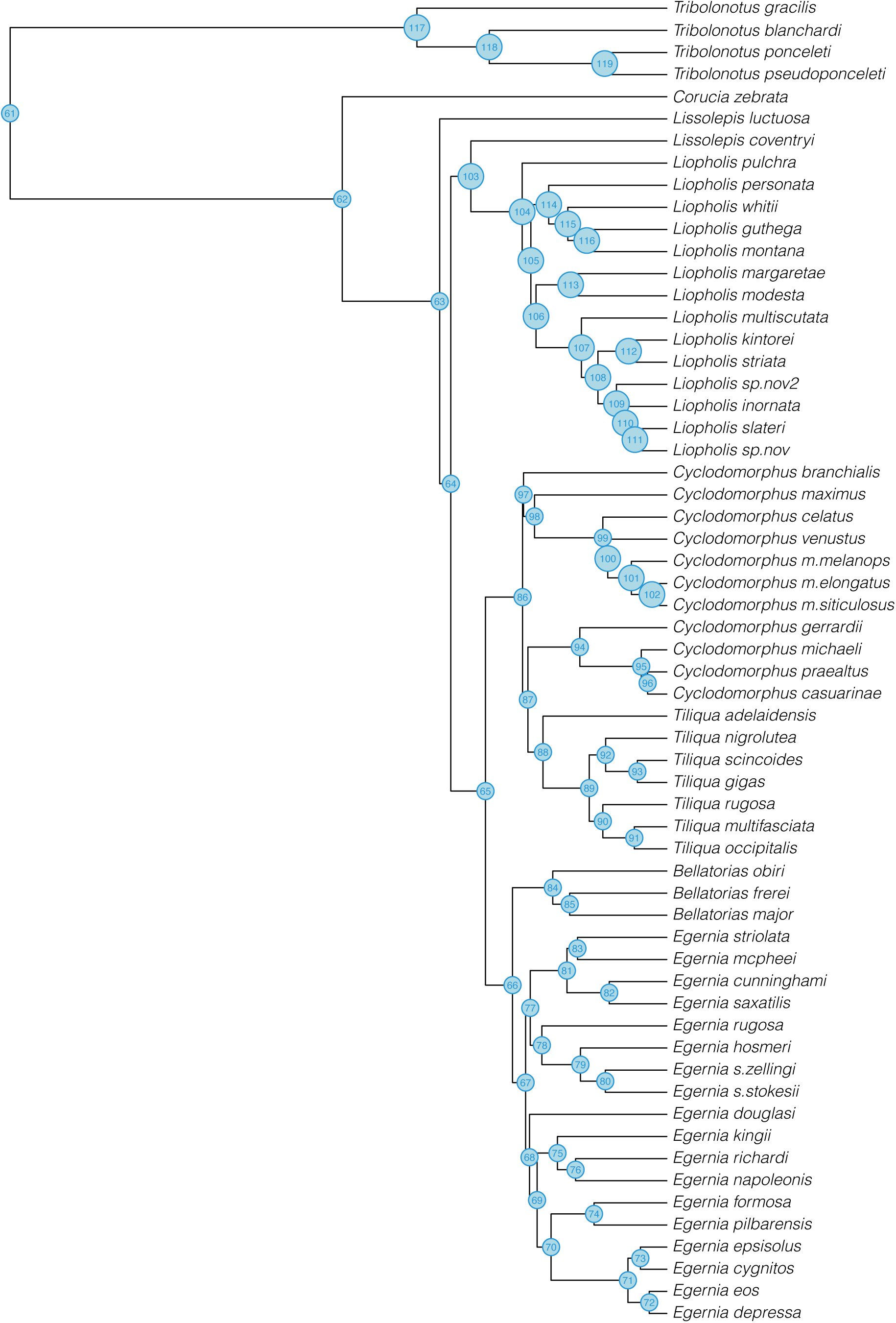
Node numbers of MCMCtree species tree used for analyses.

**Figure S29:**
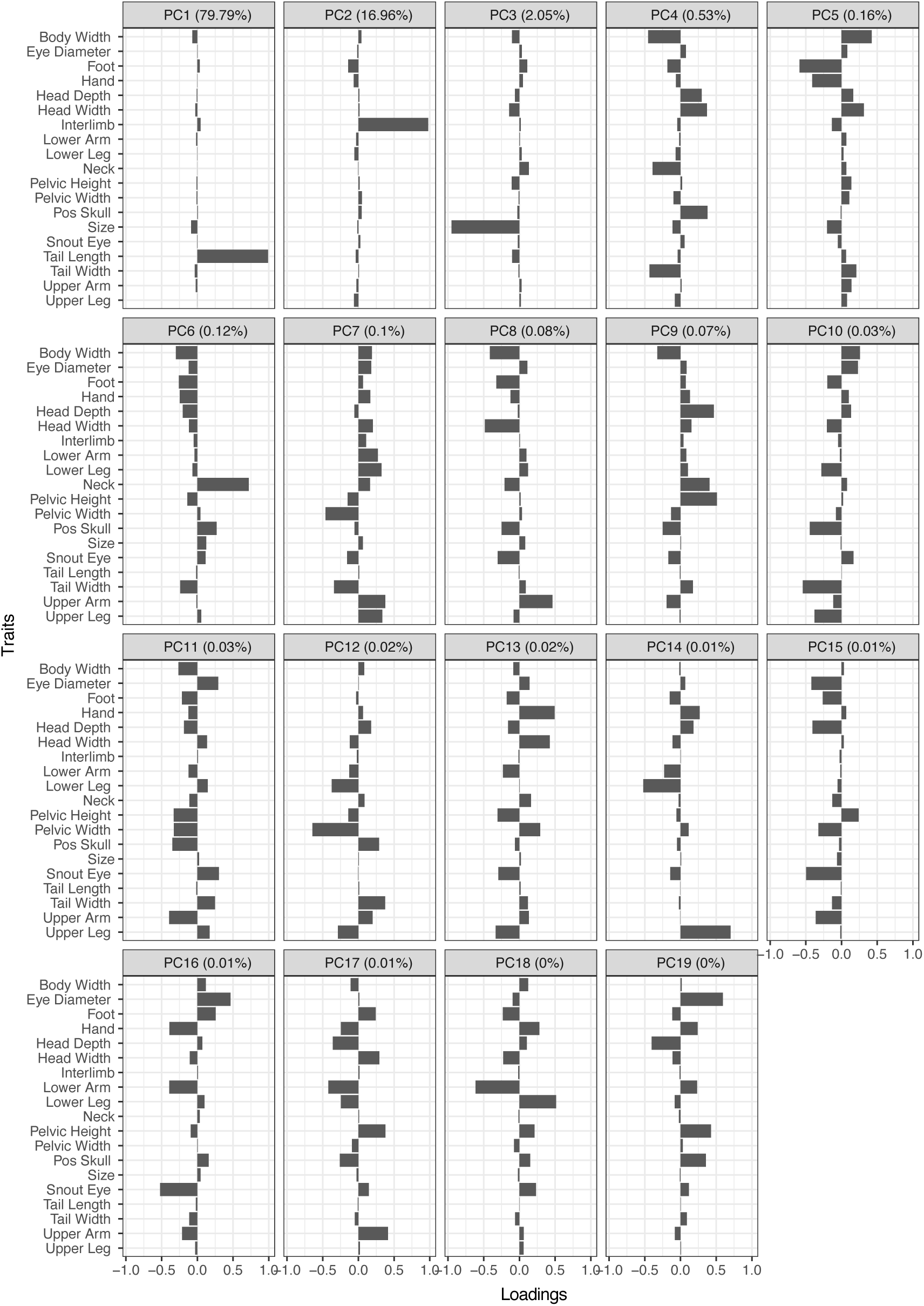
PCA loadings for 19 focal traits across all Tiliquini skinks. The first three principal components account for >98% of the variance, and are primarily explained by tail length, interlimb length, and size.

**Figure S30:**
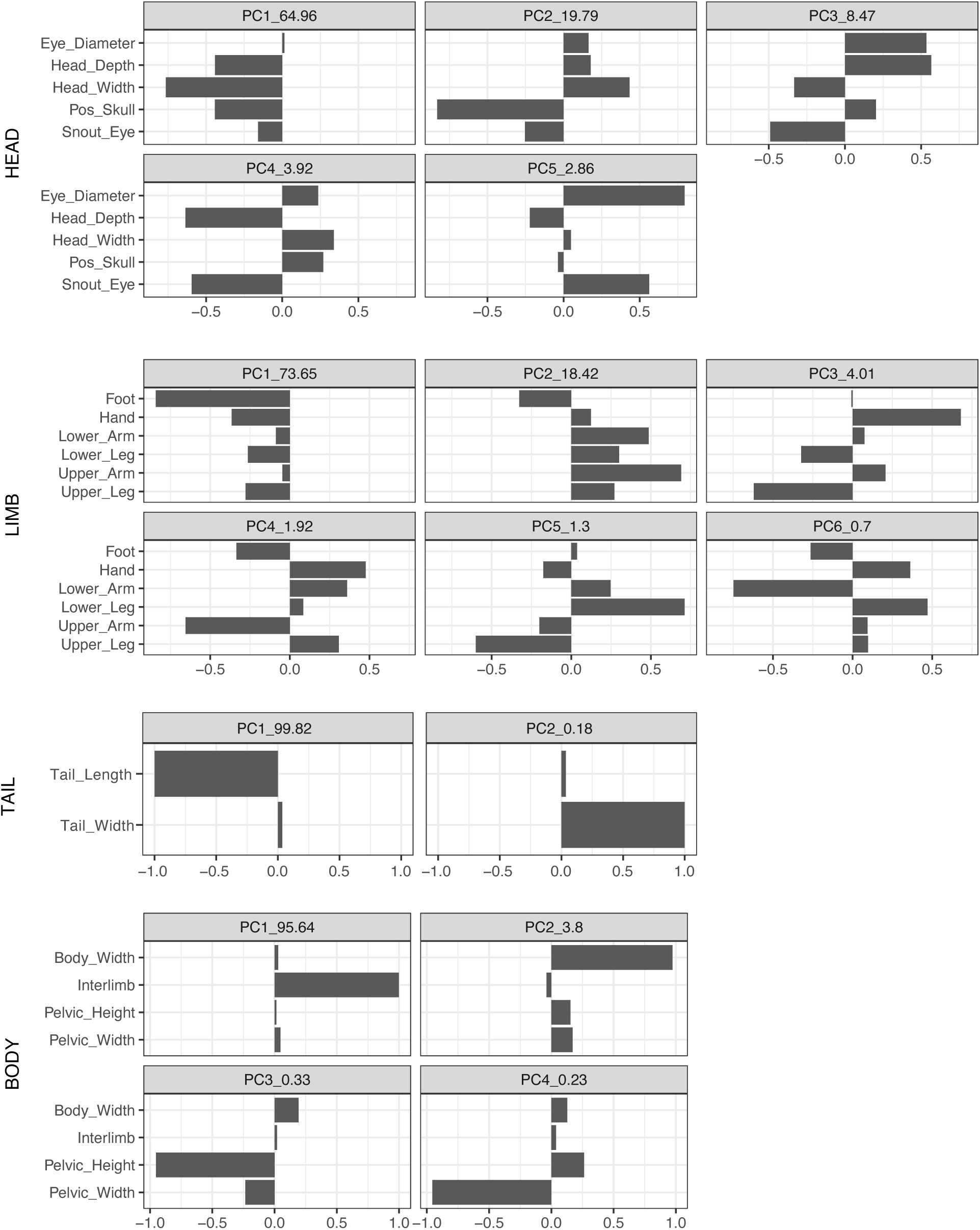
PCA loadings for 19 focal traits analyzed by module across all Tiliquini skinks.

**Figure S31:**
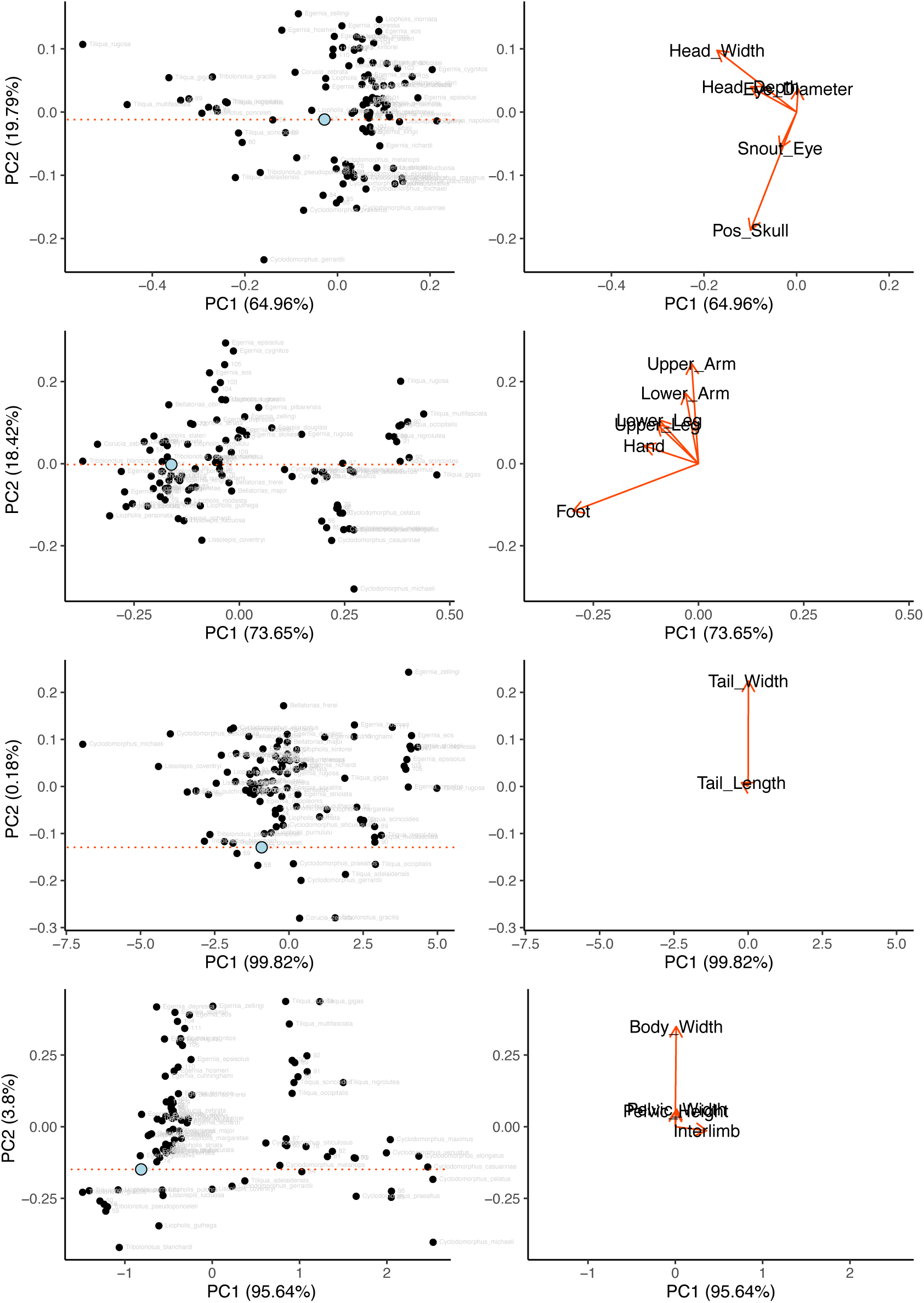
Biplots of the first two PC axes from analyses of modules.

**Figure S32:**
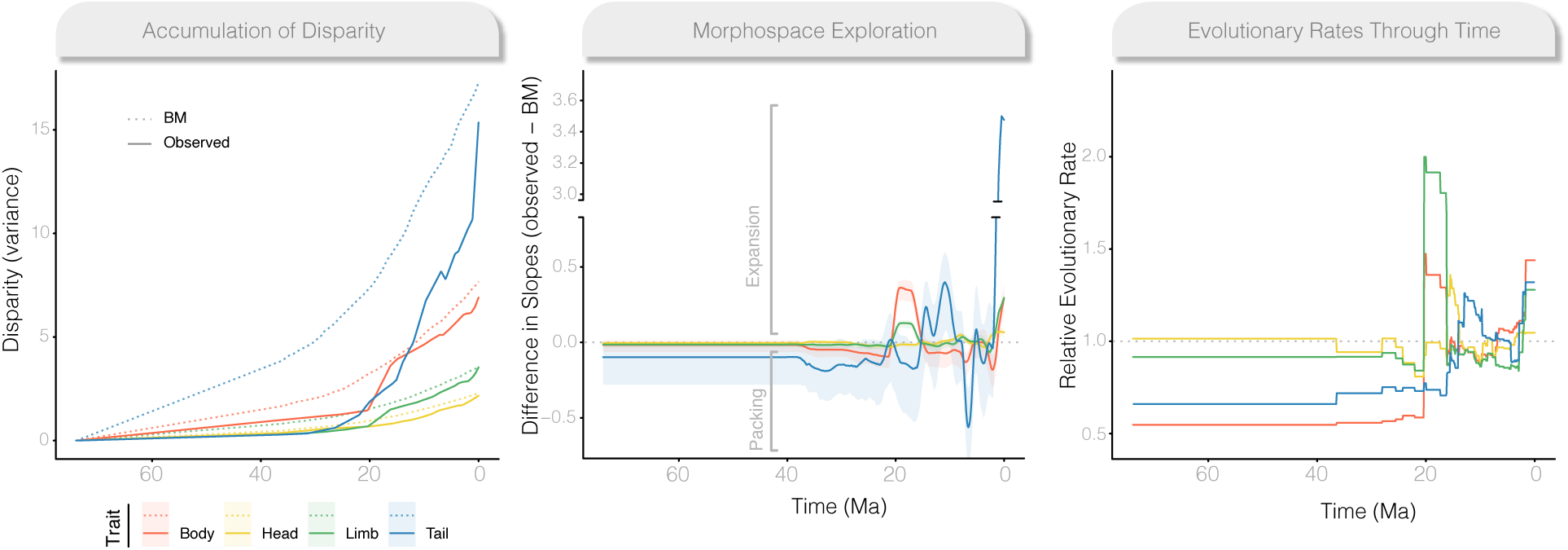
Evolutionary trajectories vary widely across modules and through time. (left) Accumulation of multivariate disparity (as variance) through time for each module (solid lines) compared to Brownian Motion (dotted lines). (center) The comparison of slopes of each module to BM highlights periods of morphological expansion (values greater than 0) and conservatism (values less than 0). (right) Evolutionary rates across modules are highly heterogeneous (see three different scales for y axis), showing periods of temporal variability, as well as high variances within modules and among traits (Fig.S24–S26). Figures represent analyses of all samples, and show some differences from Fig.3. Differences can likely be attributed to the underestimation of mean rates resulting from the long bare branches leading to *Corucia* and *Tribolonotus*.

## Notes

https://github.com/IanGBrennan/Tiliquini

### Competing Interest Statement

The authors have declared no competing interest.

